# Open-source cell culture automation system with integrated cell counting for passaging microplate cultures

**DOI:** 10.1101/2024.12.27.629034

**Authors:** Greg Courville, Shivanshi Vaid, Alexis Toruño, Paul Lebel, Joana P. Cabrera, Preethi Raghavan, Axel Jacobsen, George Bell, Manuel D. Leonetti, Rafael Gómez-Sjöberg

## Abstract

Tissue culture in 96-well microplates is conventionally a tedious, highly manual process sensitive to individual technique and experimenter error. Here, we describe the Automated Cell Culture Splitter (ACCS), a system for passaging plates of adherent or suspension cells, for routine culture maintenance or specialized applications such as seeding plates for microscopy. The system is built around the Opentrons OT-2 liquid handling robot and incorporates a novel on-deck imaging-based cell counter which allows it to compensate for density disparities across a source plate and control the number of cells seeded on a per-well basis. We find this solution can cut hands-on time by 61% and the results compare favorably to our existing manual cell culture processes in terms of both seeding density precision and biological outcomes, achieving a control of seeding density with a well-to-well coefficient of variation (CV) under 11%. The system is designed to be adaptable and an accessible entry point into automation for high-throughput cell culture; to that end, all of the source code and hardware designs are released under open source licenses.

## Introduction

Cell culture is a fundamental task in vast areas of biological research but is notoriously labor-intensive, particularly in the case of high-throughput methodologies where large numbers of samples are cultured in parallel. Reliance on manual methods can be a throughput bottleneck, a resource drain and a source of error and variability.

For instance, when manually passaging cells in 96-well plates, it is common practice to visually estimate cell density in each well based on the apparent degree of confluency of the cell layer, in order to group wells of similar density together and choose appropriate dilution factors for seeding. However, the results of such visual methods are often inconsistent – even for the same experimenter viewing the same image twice [1, 2]. In addition, subtle differences in individual technique at various steps can have marked effects on downstream results.

Meanwhile, many experiments depend on tight control over parameters such as cell density. For example, differences in seeding density can affect overall growth and confound drug sensitivity assays [3]. Another example, and the original driver of the work described here, are high-throughput imaging campaigns such as OpenCell[4], which includes 3D live fluorescence imaging of thousands of endogenously tagged cell lines. In such campaigns, reproducible imaging results are contingent on achieving an optimal and uniform cell density in each well[5].

Laboratory automation is increasingly used to address the reproducibility needs of cell culture assays[6], including robotics-assisted workflows for the culture of cells and organoids[7–9]. Increasingly, laboratory robotics is supported by a growing ecosystem of open-source hardware platforms that facilitate access and simplify integration with other workflows[10–14].

Here we present the Automated Cell Culture Splitter (ACCS), an open-source benchtop system for automating the passaging of adherent or suspension cells in 96-well plates. ACCS leverages a liquid handling robot (Opentrons OT-2) and a custom cell counting instrument which allows it to adjust seeding volumes according to cell density on a per-well basis. A single installation can passage 2-3 source plates in a working day, and the process requires minimal intervention once a run is started. The system fits inside a biosafety cabinet, and can be adapted to other uses as needs arise.

We are releasing the hardware designs and software source code for ACCS along with under open-source licenses, along with additional documentation to guide prospective builders, for others in the research community to leverage and build upon.

## Results

### System overview

Fig. 1 gives a high-level overview of the major components of the ACCS system. The design and function of each of the components described below is elaborated upon further in “Materials and Methods”; hardware components are shown in more detail in Fig. 4.

**Fig. 1.**
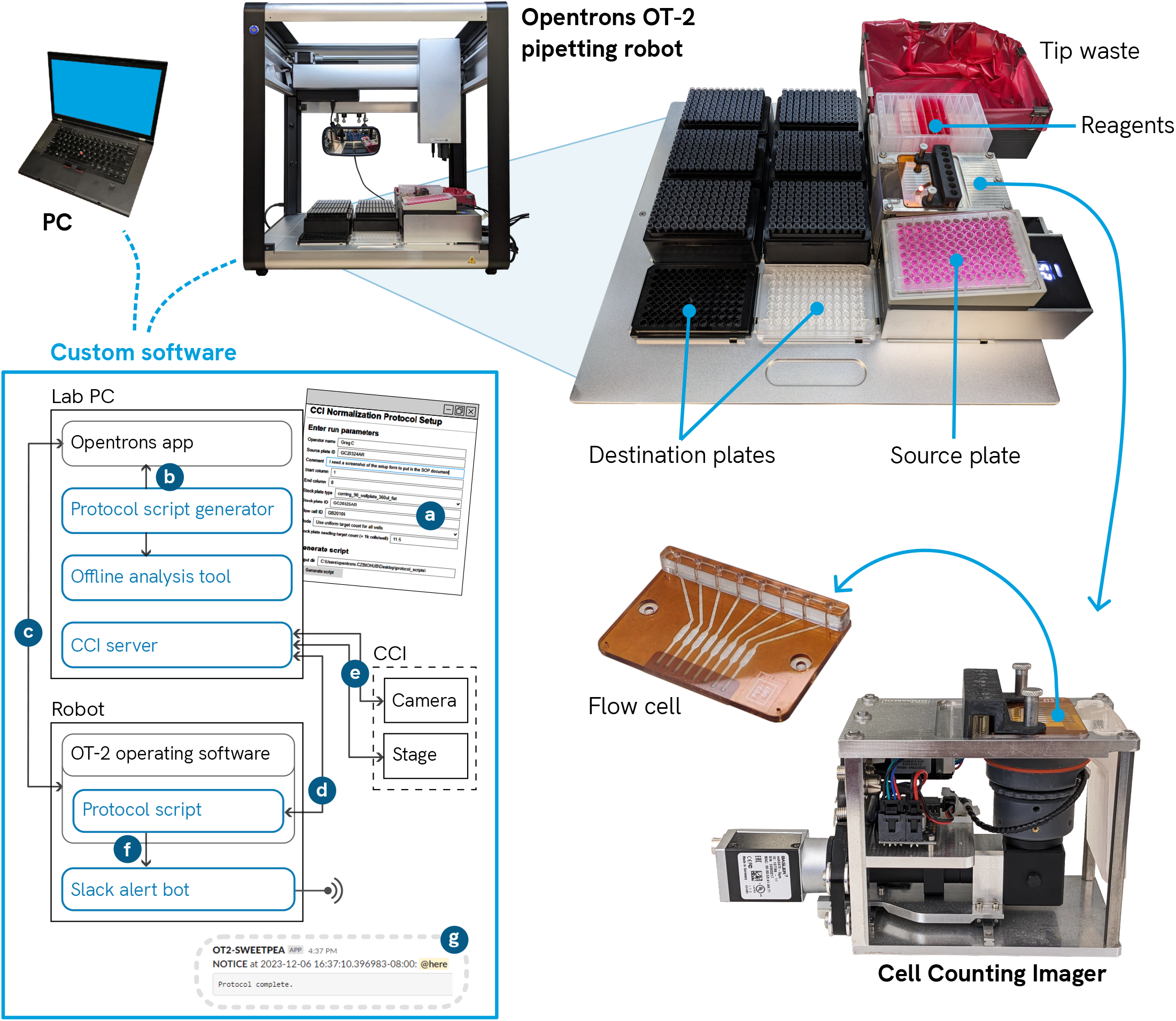
Major components. The system is built around an Opentrons OT-2 pipetting robot. The Cell Counting Imager (CCI) occupies a slot on the OT-2 deck and counts cells in samples from the source plate. A custom flow cell attaches to the top of the CCI and is loaded by the multichannel pipette. Additional custom hardware components are detailed in Fig. 4. **Inset: software overview**. Blue outlines denote ACCS custom software components. Key interactions: (a) Operator fills out the protocol setup form to generate a protocol script; (b) Operator loads the protocol script into the Opentrons app, then (c) sends it to the robot and starts the run; (d) The protocol script running on the robot sends requests to the CCI server, which in turn (e) controls the CCI hardware, captures and analyzes images, and returns cell counts; (f) The protocol script emits log events which are received by the alert bot and (g) broadcast to operators via Slack as appropriate.

#### Hardware

The major hardware components of ACCS include a minimally modified pipetting robot (Opentrons OT-2), a temperature-controlled heat block (Opentrons Temperature Module, or “Tempdeck”), and a custom instrument for measuring cell concentrations (Cell Counting Imager, or CCI). Additional custom hardware components are: An adapter to hold the source plate at an incline, a high capacity bin for tip waste, a reagent trough riser, and a pipette position calibration tool.

The CCI is placed in one of the labware “slots” on the OT-2 deck. It measures cell concentrations in 8 samples at once, to calculate the appropriate volumes of cell suspension to seed a specified number of cells in each well. Essentially, it is an inverted darkfield microscope featuring a custom flow cell with eight chambers and a motorized linear stage that positions the optical system to image each chamber. The flow cell is mounted on top of the CCI and has a row of ports to allow the multichannel pipette on the robot to inject samples from one column of the source plate at a time. Software on a host computer controls the CCI and processes the resulting images to produce cell concentration measurements.

The ACCS system requires less than 3 feet of lateral bench space and can be placed on a standard lab bench or inside a biosafety cabinet with a typical >24” deep work area. Operation is managed via a desktop computer.

#### Software

ACCS includes a suite of custom software components that work together as a system. A protocol setup software tool takes run parameters from the user via an interactive form and generates a unique script file to run on the OT-2. An “alert bot” program we add to the OT-2 captures important messages from the protocol script and alerts the operator(s) on a dedicated Slack channel. Finally, the CCI is managed by a dedicated control program running on the same computer used for protocol setup. This program directs the stage movement, captures images, processes them to count cells, and reports cell concentration calculations back to the protocol script running on the robot.

For protocol development, a custom offline analysis tool simulates protocols and checks key results such as final liquid volumes and mixtures, pipette tip usage, etc.

### Operator workflow

Two of the most common day-to-day use cases for ACCS are routine culture maintenance and seeding of microscopy plates. This section focuses on a master protocol (named `cci_normalization`) which covers these uses. Other ACCS protocols offer alternative functionalities, such as seeding by fixed volumes instead of using the CCI, and ad hoc modified protocols are sometimes made for specific needs.

To start a typical passaging run the operator follows these steps: (1) inspect, clean and install the CCI flow cell; (2) prepare the trash bin liner and install all labware (tipracks, plates, reagent reservoir) on the OT-2 deck; (3) start the CCI software, (4) set up the protocol script as described below; and finally (5) start the run via the Opentrons app.

Preparation of the CCI flow cell includes checking for damage, cleaning the top and bottom surfaces, and removing the liquid from the channels. The reagent reservoir is filled with necessary reagents (DPBS, trypsin, growth media, water, cleaning detergent) in a standardized format. Reservoirs can be filled in bulk ahead of time, sealed, and stored under refrigeration to reduce setup time.

The protocol script setup process starts with the operator filling out a browser-based setup form, then the ACCS software assembles a unique script file with the instructions for the robot to carry out the protocol as well as metadata about the run for later reference. The Opentrons desktop app is used as normal to send the script to the robot and start the run. While individual protocol scripts can be reused, in practice a new script is usually generated each time to capture and save specific run details, such as identities of the plates, etc. Once started, the process requires minimal intervention, allowing the operator to work on other tasks. Alerts are sent to a Slack channel for protocol start, completion, and any situations requiring operator attention.

Supplement D is an example SOP document explaining the operator steps to run the `cci_normalization` protocol in detail.

### Cell density-related metrics

In many cases the quality of downstream results may depend on consistently seeding cells at a particular density. A key feature of the system is the ability to compensate for variation in cell density between wells on the source plate by measuring the concentration of resuspended cells in each source well and using that information to transfer the appropriate volume of media and cell suspension to the destination well. Here we examine aspects of the system’s quantitative performance with respect to this task.

#### Basic CCI cell counting performance

The system’s ability to seed a controlled number of cells in each output well is dependent on being able to accurately measure the concentration of cells in suspension in the source well after dissociation.

The basic counting performance of the CCI, in particular the linearity and dispersion of measurement values, was characterized by measuring a series of prepared HEK293T stocks in parallel on both the CCI and a flow cytometer, with the latter being used as a ground truth for cell concentrations. Results are illustrated in Fig. 2(A).

**Fig. 2.**
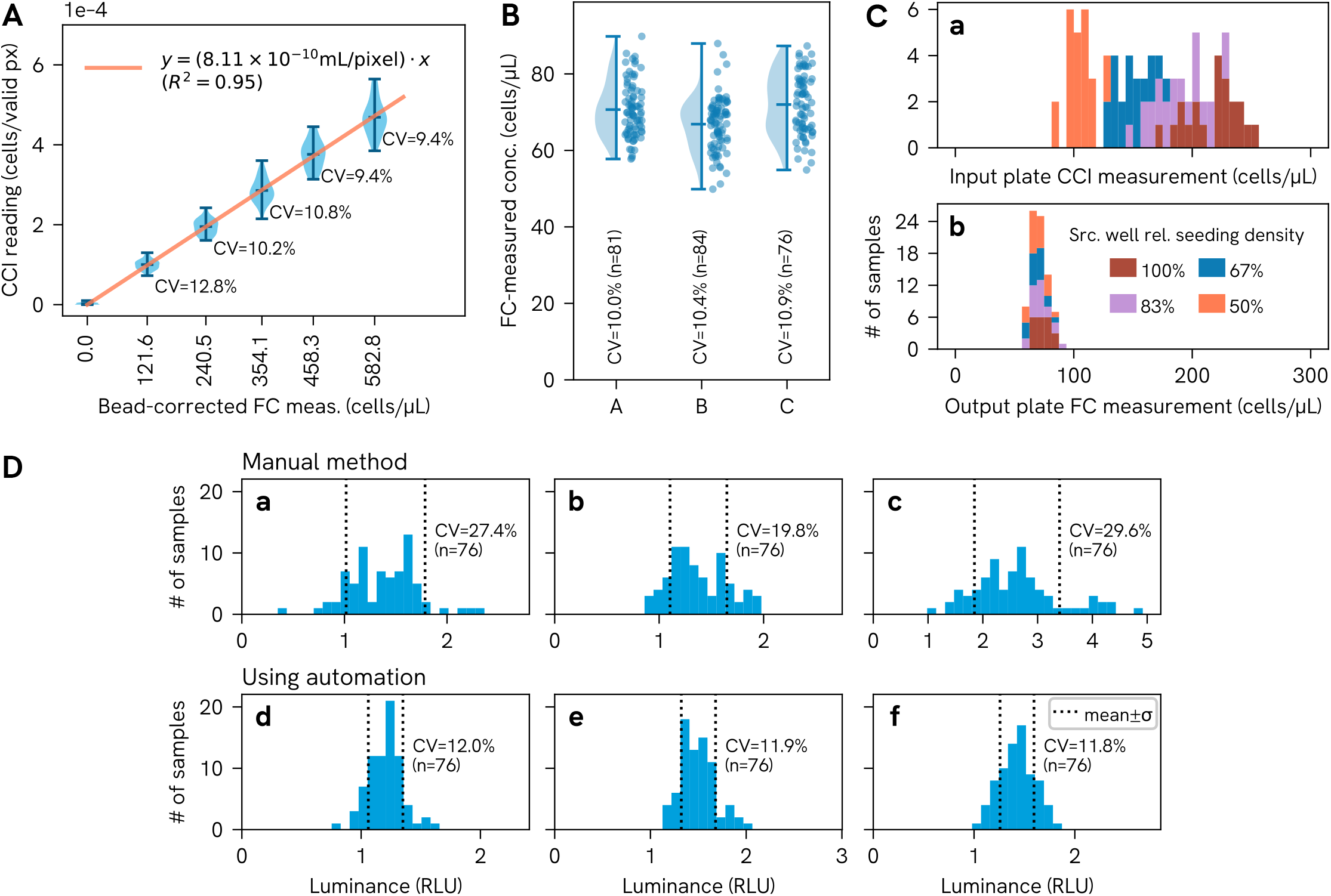
(A) CCI counting performance. Distribution of CCI readings for each dilution plotted against cell concentration values obtained by bead-corrected flow cytometry measurements, with a linear regression through the origin used to generate the “calibration factor” for the CCI (the slope of the fit line); **(B) Cell density target performance**. Distribution of cell concentrations measured in three test plates seeded by ACCS, using an artificially diverse source plate, as measured by flow cytometry (“FC”) immediately after seeding; **(C) Input plate vs output plate cell concentration**. Distribution of source plate cell concentration readings from the CCI compared to concentration measurements taken from the output plate by flow cytometry (“FC”), for the dataset corresponding to sample A in subfigure B; **(D) Seeding density consistency for ACCS vs manual process**. Distribution of raw luminance values from an ATP assay (CellTiter-Glo) used as a proxy for relative cell density for three plates seeded manually based on visual assessment of the confluency of the source wells ((a)-(c)) and three plates seeded automatically by ACCS ((d)-(f)).

It should be noted that the variance in CCI readings here is potentially exaggerated by the fact that measurements were aggregated over multiple cycles of sampling with significant time in between; the measurement procedure is described in greater detail in “Materials and Methods”.

Although in principle we can calculate the volume “seen” by the imager from design parameters, in practice we use an empirical calibration against a chosen reference instrument to compensate for non-ideal factors (software over-/undercounting, flow cell thickness variation, etc); the slope of the fit line through the data in Fig. 2(A) is equal to this “calibration factor,” which represents the ratio of physical volume to image pixels in the analyzed portion of the image. In practice we currently do not find per-flow-cell or per-channel calibration worthwhile as counting variation is dominated by other influences (sampling error, biological variation, etc.).

#### Control of seeding density

To examine how consistently the system can actually seed cells at a specified density, we first seeded a test plate with an artificially large spread of cell densities (which we call “Challenge Plate”), incubated it overnight, then used ACCS to passage that plate into a U-bottom plate with a fixed seeding target, and finally counted a sample of cells in each output plate well by flow cytometry to determine the cell concentration. The seeding target was set at 12.5k cells/well in 200 µL total media volume, similar to values we use when seeding microscopy plates for OpenCell.

Fig. 2(B) shows the results from three such experiments; Fig. 2(C) further illustrates the ability of the system to produce well controlled seeding densities from highly disparate input, comparing the cell concentrations measured by flow cytometry in one of the output plates to those measured in the corresponding input plate by the CCI during the harvesting and passaging process.

#### Seeding density consistency compared to manual method

For the system to be useful it should perform at least as well as the manual method in seeding cells at a consistent density, so we directly compared plates seeded manually to plates seeded using ACCS on the basis of viable cell density.

Multiple plates were seeded both by hand and using ACCS, using the same Challenge Plate configuration and seeding target as in the previous experiment, and incubated overnight. The plates were then analyzed using a CellTiter-Glo ATP assay the following day to assess the relative density of viable cells in each well. As illustrated in Fig. 2(D), the resulting data shows that the plates processed by the automated system had tighter distributions (54% lower coefficient of variation on average) compared to those produced by visually estimating confluency for each well and passaging by hand.

### Throughput and hands-on time

For many pipetting tasks, a skilled human experimenter can move more quickly than the OT-2. However, the throughput benefit of the automated system lies in the reduced active engagement time for the operator. Manually passaging a single plate may take an hour or more of continuous work, while using ACCS can reduce this to under 30 minutes of hands-on time (depending on experience level and circumstances), with a typical end-to-end time of under 3 hours.

The tradeoff is illustrated in Fig. 3(A), which shows the range in hands-on and end-to-end time for two subjects using ACCS to passage a plate, compared to three subjects passaging by hand. While the end-to-end time for a run, including the setup, teardown and robot operation, is significantly longer than for the manual process, there is a substantial period of “walk-away” time during which the operator is able to attend to other tasks (potentially including setting up parallel runs on additional robots).

**Fig. 3.**
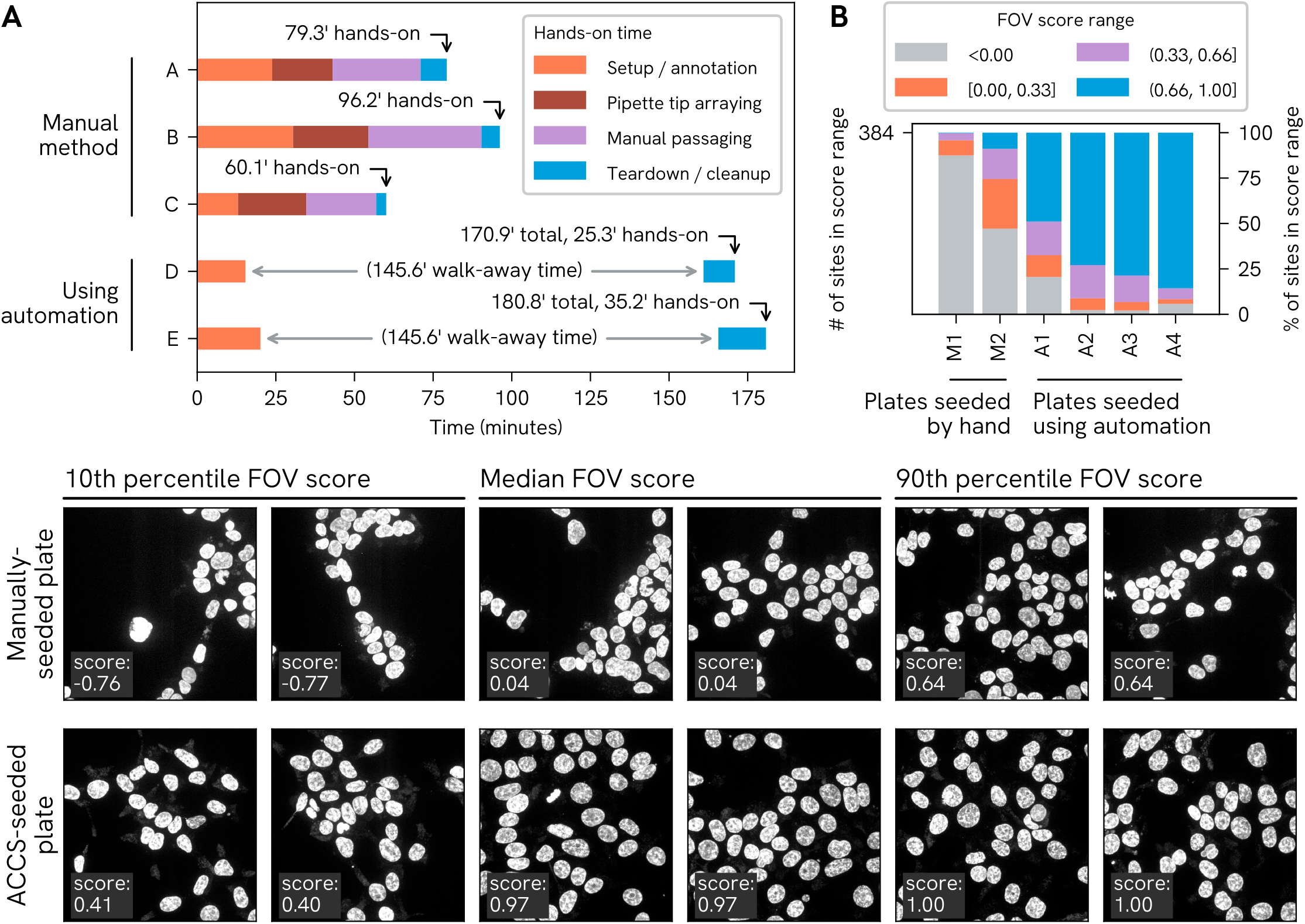
(A) Total duration and walk-away time. Illustration of the tradeoff between end-to-end duration and hands-on time for passaging of a 96-well plate by the conventional method, with split factors determined by visual confluency estimation (samples A-C, corresponding to three different testees), compared to performing the same task using ACCS (samples D-E, corresponding to two different ACCS operators); **(B) OpenCell imaging FOV score performance**. Comparison of FOV score distributions obtained from two imaging runs where the imaging plate was seeded manually, followed by four consecutive and most recent runs (with the same plate layout) where the plate was seeded using ACCS; **(C) Example imaging results for manually-seeded and ACCS-seeded wells**. A selection of Z-projections generated from the DAPI channel data for image stacks acquired from a manually-seeded plate (left, corresponding to “M2” in subfigure B) and an ACCS-seeded plate (right, corresponding to “A4”). To illustrate the range of results obtained for each condition, as well as the relationship between image features and FOV score, images are programmatically selected based on their corresponding FOV scores to represent “high” (90th percentile), median and “low” (10th percentile) scores for each dataset.

### Outcomes in a practical scenario

One of the original driving use cases for ACCS was preparing plates for the high throughput imaging pipeline in OpenCell. At the start of the imaging process, custom microscope automation software takes a set of images of the DAPI-stained nuclei over a grid of candidate sites in the middle of each well, in order to assess local cell density. A supervised machine learning model, trained on representative images of desired density, assigns an “FOV score” to each field of view (FOV) (see [4] for further detail) which is used to select the most favorable sites at which to run a full multidimensional acquisition. Scores range from –1.0 to 1.0, and in practice only FOVs with positive scores are considered usable. Typically, the four highest-scoring sites in each well are selected; therefore, the following analyses consider the aggregate of all FOV scores from each plate which fell in the four highest in their corresponding well. Sites which could not be scored are treated here as having a score of –1.

Fig. 3(B) illustrates the distribution of FOV scores for selected imaging sites (4 highest-scoring sites per well) for the four most recent imaging runs on ACCS-seeded copies of a particular OpenCell plate, compared to historical data from two copies of the same plate (only two such datasets exist for comparison) that were seeded manually. Meanwhile Fig. 3(C) visually illustrates the relationship between FOV scores and the spatial distribution of nuclei as well as the range of quality observed on the best ACCS-seeded copy vs the best manually-seeded copy. On the whole, the datasets from ACCS-seeded plates collectively had a 92% “keeper rate” (score > 0) compared to 52% for the most successful hand-seeded plate.

## Materials and Methods

### Overview of system components

The ACCS system comprises several major hardware and software components: the robot protocol suite and underlying software framework, the Cell Counting Imager (CCI) with its associated flow cell and control software, and a small number of bespoke hardware accessories to adapt the OT-2 to the task of passaging 96-well plates.

The following subsections give a brief, high-level description of the aforementioned components. Additional technical details and practical guidance for assembling a complete system are provided in the *ACCS Integrator’s Manual* (supplement B). The design, construction and operation of the CCI is described in the *Cell Counting Imager Technical Manual* (supplement C).

### ACCS automated passaging protocol

In brief, a standard ACCS passaging protocol consists of similar steps to what would be done by hand. The robot uses the multi-channel pipette to wash the cells, trypsinizes the cells, quenches the trypsin and resuspends the cells, then for each column on the source plate it takes cell concentration measurements using the CCI and transfers the appropriate amount of cell suspension into the destination plate with the single-channel pipette according to the specified seeding target.

See the Supplemental Materials and Methods (supplement A) for an in-depth description of typical steps in an ACCS protocol.

### Protocol run-time framework and tooling

The OT-2 features a Python scripting API as the primary means to develop custom protocols. The ACCS protocol framework wraps and extends the Opentrons API, abstracts certain common operations, and adds various extra facilities for configuration, logging, and offline analysis. It also includes special features such as a mechanism for allowing tipracks to be replenished during a run and a mode that allows moving the Gen2 pipette plungers at dramatically higher speeds than normally possible. The latter was crucial for reaching the flow rates necessary for effective dissociation of cells.

ACCS protocol scripts automatically generate a “paper trail” on the robot’s filesystem for every run, including a record of log messages, a JSON file with summary information, and a copy of the script itself. The logging system also feeds messages to a Slack bot that alerts users to critical events.

The user-facing protocol setup tool presents a browser-based form for data entry and outputs a self-contained protocol script file which is unique to a given run and contains all of the necessary framework code as well as user-entered parameters and the code for the protocol itself.

For offline development, a custom analysis tool extends the simulation functionality of the Opentrons software and allows tracking the movement and mixture of liquids as well as pipette tip usage and can be used to “sanity check” protocols or, for example, to predict the amount of reagents and pipette tips needed for a given run.

### The Cell Counting Imager

#### High-level description

Fig. 4(A) identifies the major components of the Cell Counting Imager. The CCI is essentially an inverted darkfield microscope with 4X magnification and a motorized stage, built in a compact form factor which mounts in an SBS compatible slot on the deck of the OT-2. Cell samples are loaded in a purpose-made flow cell (see “CCI flow cell“) by the robot’s multichannel pipette, images are taken of each chamber, then the fluid channels are flushed out by the pipette before loading the next sample. Images are processed to produce a cell density measurement for each channel.

**Fig. 4.**
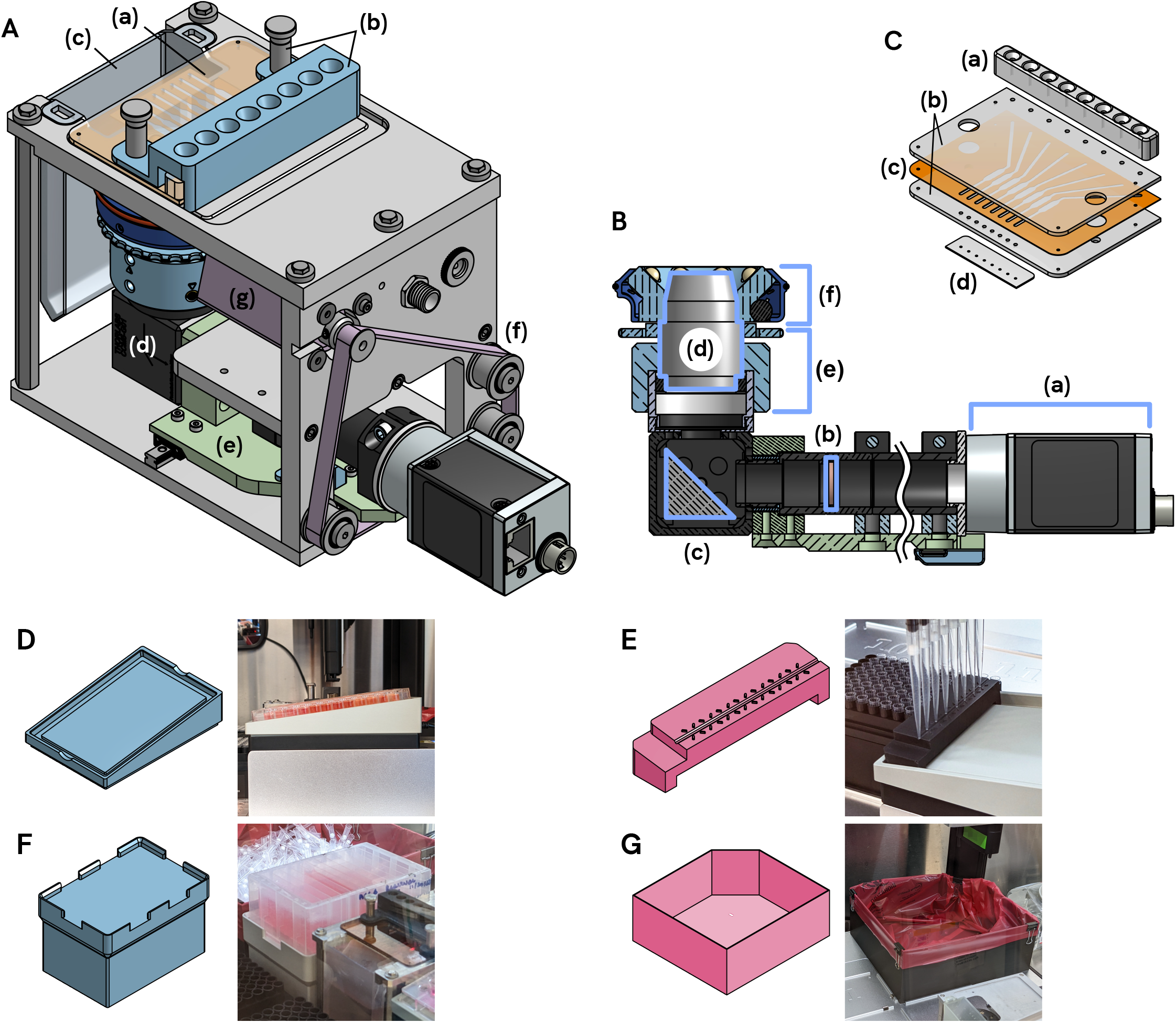
(A) CCI key mechanical components: (a) flow cell, detailed in panel C, (b) pipette tip guide and flow cell hold down fixture, (c) waste trough, (d) optical system - detailed in panel B, (e) linear stage carries the optical system, (f) timing belt system moves the stage, (g) stepper motor drives the timing belt; **(C) CCI flow cell construction:** (a) Pipette tip interface is made of elastomer material and mates with the tips on the multichannel pipette for injection of samples, (b) Top and bottom slides form the outer structure and optical windows of the flow cell; (c) Spacer layer forms the flow channels and imaging chambers, (d) Drain port strip prevents “wicking” of fluid out of the flow cell; **(D) Inclined thermal block adapter** holds the source plate at a fixed angle on the Tempdeck; **(E) Pipette position calibration tool** facilitates calibrating the coordinates of the source plate in the Opentrons labware calibration flow; Reagent trough riser eases access to the reagent trough by locating it in an elevated position and allowing it to simply drop into place rather than interacting with spring clips on the OT-2 deck; **(G) High capacity replacement tip waste bin** is required in order to accommodate the large amount of tips used in passaging protocols.

#### Optical system

Fig. 4(B) shows the main components of the CCI optical system. The CCI uses a commercial 4X plan objective to image cells resting on the bottom of the flow cell chambers. The optical system is L-shaped to satisfy mechanical constraints, with a first surface mirror used to orient the objective vertically. Samples are illuminated by a ring of eight 740 nm near-IR LEDs and a 700 nm longpass filter in front of the camera is used to minimize the influence of ambient light. Images are captured by a Basler acA5472-5gm monochrome CMOS camera. Focus adjustment is provided by a 3D-printed knob and collar.

#### CCI flow cell

The CCI flow cell is a hemocytometer-like device with a row of 8 imaging chambers and channels allowing samples to be injected and flushed out. The camera’s nominal FOV is a ∼2.4 mm × 3.6 mm rectangle within each 6 mm × 2.7 mm chamber. After masking, CCI images effectively cover about 1 µL of sample volume.

An exploded view of the flow cell is shown in Fig. 4(C). The construction consists of two poly(methyl methacrylate) (PMMA) plates sandwiching a ∼0.24 mm thick adhesive spacer layer that forms the fluid paths. The flow cell was designed to be manufacturable using a consumer-level CO2 laser engraver, however the PMMA plates are currently produced by CNC machining.

A “pipette interface” device molded from elastomeric material is bonded to the top side of the flow cell, and is used to form a seal with the pipette tips for the robot to inject liquid into the flow cell. A “nozzle strip” device is attached to the drain ports in order to prevent the liquid in the flow cell channels from wetting the outside surface of the flow cell and seeping out. This is accomplished using small exit holes surrounded by a super-hydrophobic surface produced using a laser in a manner similar to that described in [15].

#### Mechanical design and electronics

The frame of the CCI is made up of machined aluminum plates. The mechanical structure of the optical system is a combination of ThorLabs components and custom machined parts. A pair of linear rails support a single axis positioning stage which is driven by a small (NEMA 8) stepper motor via a timing belt. A Pololu Tic T249 stepper motor controller powers the motor.

The darkfield illuminator contains eight LEDs and a current regulator circuit in a 3D printed frame which attaches to the nose of the objective.

The flow cell is attached to the top of the CCI using thumbscrews and a 3D printed fixture which also protects the pipette interface from misaligned pipette tips. A removable 3D-printed waste trough mounts beneath the flow cell to collect fluid from the drain ports.

#### Control software and image processing

The control of the CCI stage and processing of images is handled by a Python program on the same computer used to set up the OT-2. The software exposes a simple HTTP API that services requests from the protocol script running on the robot. After priming the CCI flow cell with DPBS, the protocol script commands the CCI to perform a “background scan”, which takes an image of each channel with no cells present. Then, after loading each sample, the protocol script commands the CCI to perform a “counting scan,” after which it receives back the calculated cell concentration values for each channel. All images captured are stored on disk for potential later review, alongside diagnostic plots and measurement values.

Fig. 5(A) illustrates what the CCI camera “sees” when imaging a sample. The image processing steps used to measure cell concentration are outlined in Fig. 5(B). Each sample image is pre-processed to remove artifacts using a series of masking strategies to exclude problematic areas or features, then a cell detection routine based on cross-correlation and local peak finding is used to identify the locations of cells in the “valid” parts of the image, and finally a cell concentration value is calculated taking into account the volume represented by the “valid” areas of the image.

**Fig. 5.**
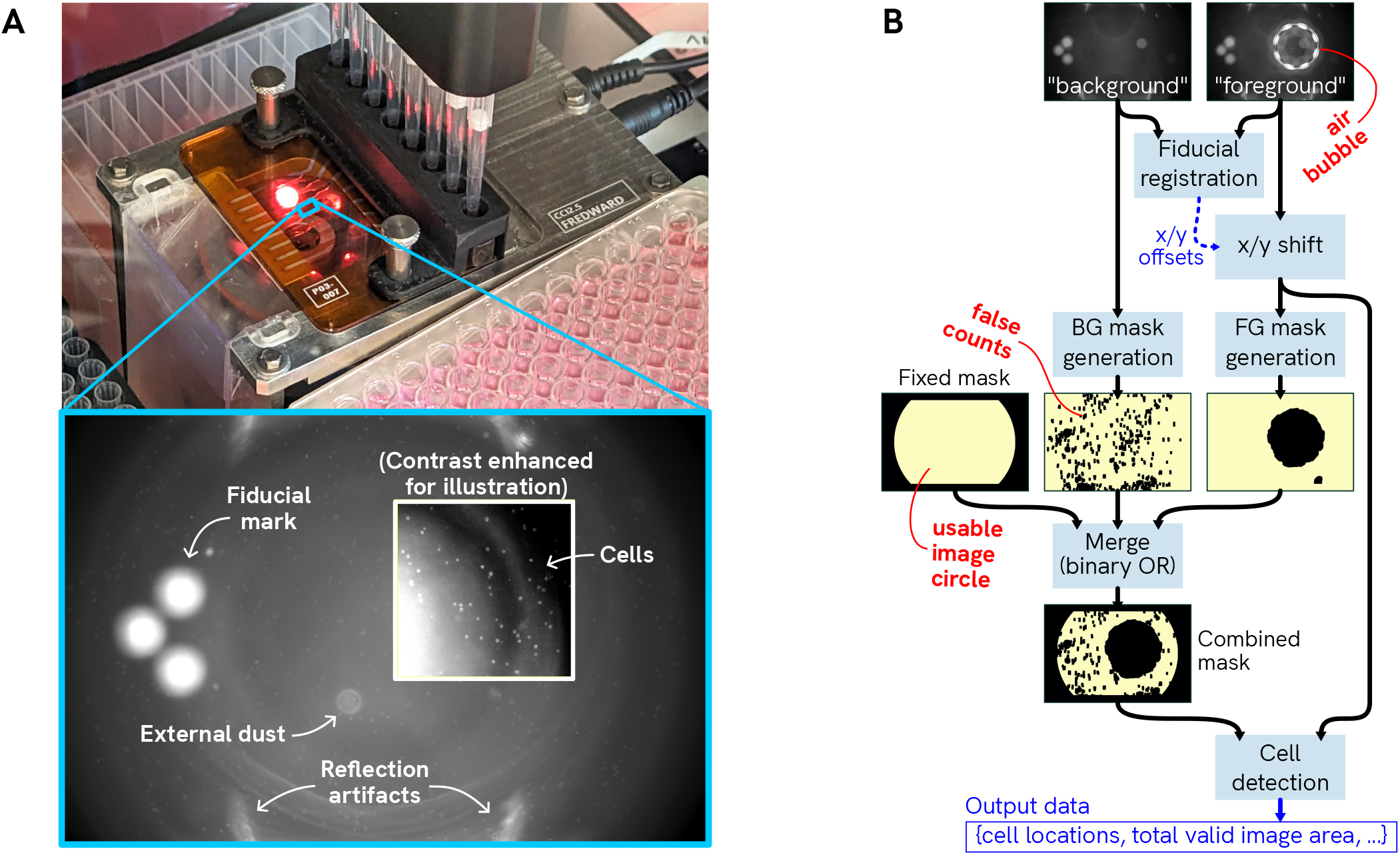
CCI counting process. **(A) CCI performing an imaging cycle**, with inset showing an example of what the CCI camera sees; **(B) CCI image processing overview**. Simplified high-level outline of processing steps. Each “foreground” image (sample to be measured) is aligned against a stored “background” image (cell-free reference) and then a mask is generated to exclude problematic fixed (e.g. flow cell debris) or transient (e.g. air bubbles) features from the analyzed volume. An additional fixed pattern is added to the mask to exclude areas outside the image circle or close to the channel walls. Finally the cell detection routine processes the foreground image to count cells within the image regions deemed “valid” per the mask.

Most parameters for the various image processing steps are controlled by configuration files, allowing for per-instrument tuning or switching between cell types. Generally, objects with a similar or larger diameter to HEK293 cells (∼15 µm)[16] and a sufficient reflective signature should be suitable. The CCI is typically used with cell concentrations between roughly 50k and 500k cells/mL. The useful lower bound is dictated by the acceptable error for the application; for example, the best-case statistically achievable CV is ∼10 % at a sample concentration around 100k cells/mL and gets worse with decreasing concentration. The upper limit of cell concentration that the software can reliably process has not been explored. For a more thorough discussion of CCI image processing see the *CCI Technical Manual*.

### OT-2 robot configuration and modifications

The liquid handling platform for ACCS consists of an Opentrons OT-2 liquid pipetting robot with one Opentrons P300 Gen2 8-channel pipette and one P300 Gen2 single-channel pipette.

A small set of passive add-on hardware components are used in addition to the CCI (see “Add-on hardware“). These items are simply placed in position and can be easily removed in order to use the OT-2 for other purposes. The only physical modification made to the robot, which is easily reversible, is the removal of the built-in trash bin.

The Opentrons robot software and desktop app version used for ACCS are version 4.7 (newer versions have not yet been tested). One file is added to the root filesystem on the robot to enable the ACCS Slack alert bot to run as a system service (optional). Otherwise all ACCS related files are contained to the user-writeable area used by the onboard Jupyter Notebook environment.

### Add-on hardware

The ACCS system also includes a few additional custom hardware components as seen in seen in Fig. 4(D)-(G): An adapter block (Fig. 4(D)) goes between the source plate and the Opentrons Temperature Module in order to hold the source plate at an 8° tilt while maintaining thermal contact. The tilt allows the pipettes to fully aspirate the well contents and to dispense PBS down the sidewall for the washing steps. The geometry of 96 well plate wells allows us to avoid the complexity of a variable tilt mechanism as commonly seen in automated cell culture systems [7, 17], since the well bottoms remain covered by media despite tilting.

A calibration tool (Fig. 4(E)) is used during the Opentrons labware calibration flow to help position the pipette correctly with respect to the source plate location.

The reagent trough riser (Fig. 4(F)) holds the reagent trough in an elevated position and makes it much easier to install and access compared to mounting it directly on the robot deck.

The extended tip waste bin (Fig. 4(G)) replaces the OT-2’s original waste bin in order to accommodate the large amount of tips used in the passaging protocols.

### Instruments and third party software

Cell concentration measurements for preparing dilutions were performed using a Invitrogen Countess II FL automated cell counter (AMQAF1000) and disposable counting chamber slides (C10228).

Flow cytometry measurements were performed using a Becton Dickinson FACSymphony A1 flow cytometer with High Throughput Sampler, using FACSDiva software for data acquisition. Analysis was done with Python and the FlowKit library[18].

Luminescence measurements were performed using a Molecular Devices Spectramax i3x multi-mode microplate reader and SoftMax Pro 7 software.

Live cell imaging is carried out on an Andor Dragonfly high speed confocal microscope system using the microscopy automation software developed for OpenCell (https://github.com/czbiohub-sf/2021-opencell-microscopy-automation).

Data visualizations including those included in this manuscript were generated using Python and matplotlib[19].

### Reagents and cultures

Live cells used for ACCS quantitative performance testing were wild type HEK293T cells, except that one run of the seeding density control experiment (corresponding to dataset “A” in Fig. 2(B)) was carried out with a HEK293T cell line expressing the split fluorescent protein mNG1-10. For both manual and automated protocols, cells are washed with DPBS without calcium and magnesium (Gibco, 14190-144); the dissociation agent is 0.25 % Trypsin-EDTA (Gibco, 25200-056).

Growth media for ACCS operations (except where otherwise noted) consists of DMEM, high glucose with GlutaMAX (Gibco, 10566-016); plus 10% v/v fetal bovine serum (FBS); plus 25 mM HEPES (Gibco, 15630-080) plus 100 U/mL penicillin-streptomycin (Gibco, 15140-122). Media used to seed imaging plates contains a greater portion of FBS at 20% v/v. Media used during manual passaging omits HEPES. Media used for Hoechst staining and live cell imaging substitutes DMEM without phenol red.

For nuclear staining prior to live cell imaging, Hoechst 33342 (Thermo Fisher, H3570) is used at 1 ng/µL in phenol red-free medium as described above.

Fibronectin used in imaging plate treatment is Corning 356008.

The internal cleaning agent for the CCI flow cell is Contrad 70, 10 % v/v in deionized water.

For counting using the Countess II FL, cells were stained with 0.4 % trypan blue (Invitrogen, included with counting slides C10228)

### Consumable labware

Routine culture maintenance uses BioLite 96-well plastic microwell plates (Thermo-Fisher, 130188). Imaging plates are Cellview 96-well glass-bottom plate (Greiner Bio-One, 655891).

Plates for ATP assays used opaque white tissue culture-treated polystyrene plates (Falcon, 353296). Plate seals used during measurement were Bio-Rad Microseal ‘B’ optically clear PCR plate sealing film (MSB1001).

### Manual cell culture protocols

#### Passaging of adherent HEK cells in 96 well plates

All incubation steps are at nominally 37°C and 5 % CO2.

The source plate is inspected in brightfield at 4X magnification, the confluency of cells in each well is estimated, wells with similar confluency are grouped together (typically no more than 4 or 5 groups), and a split factor is assigned to each group. The plate is returned to the incubator and pipette tips are arrayed according to the well positions for each group.

The old media is removed from the wells and the cells are washed with 100 µL of DPBS. To avoid dislodging the cell layer, washing is done by holding the plate at an incline, gently dispensing DPBS onto the sidewall of the well, and aspirating from the bottom corner.

50 µL of dissociation reagent is added and the plate is incubated for 5 minutes. Then 200 µL of growth media is added to quench the dissociation reagent and the well is mixed through several pipette strokes to help dissociate the cells. Finally, working in rows, the pre-arrayed tips are used to transfer a volume of media to the destination plate followed by a volume of cell suspension from the source plate.

#### Live cell imaging preparation

Glass-bottom plates intended for imaging are treated before seeding by filling with 70 µL of fibronectin solution at 50 µg/mL and incubating at ambient temperature for 45 minutes, then washing with 100 µL of DPBS. Plates prepared well ahead of the time of use are filled with media and stored at 4°C.

Prior to imaging, cells undergo nuclear staining and media replacement as follows: Media is removed from the plate and replaced with a staining medium containing Hoechst 33342 nucleic acid stain, the plate is incubated for 30 minutes at 37°C, then the staining medium is removed and replaced with phenol red-free imaging medium (see “Reagents and cultures”). The plate is incubated for at least 3 hours before starting imaging.

### ACCS performance testing procedures

#### “Challenge Plate”: spatially randomized test plate

For experiments related to seeding density control we used a “Challenge Plate” which is a plate seeded with a randomized arrangement of different cell densities. A Challenge Plate is prepared using a purpose-made ACCS protocol script which seeds from a reservoir of prepared cell stock using the single-channel pipette. The plate map consists of four well groups arranged in a pseudo-random pattern across the plate, with each being seeded at a different relative density ranging linearly from 50% to 100%. Supplemental Figure S1 illustrates the layout of the Challenge Plate.

Cells are manually harvested from a flask and diluted to prepare a 20 mL cell stock at nominally 200k cells/mL, from which the robot seeds between 100 and 200 µL into each well (corresponding to a range of 20k to 40k cells), topping up with media to 200 µL total volume. The plate is incubated for ∼24 hours before use.

#### Procedure for hands-on time study

For the time study comparing the overall duration and “hands-on” time for manual passaging compared to using ACCS, three different individuals passaged a plate manually and two different individuals (at the time of writing this was everyone available and proficient in operating the system) used ACCS to do the same. Both processes were carried out in the same way as for the seeding accuracy comparison experiments as described in “Procedure for comparative seeding accuracy tests by ATP assay“, except that a normal culture plate was used for the output plate and there were no special reserved wells.

Starting conditions were as follows: biosafety cabinet open, wiped down and “warmed up” ahead of time; reagents ready at appropriate temperatures (media and dissociation reagent warmed to 37°C, all others RT); all necessary equipment and consumables staged within reach.

An observer marked the time at transition points between tasks in each process. Timing for both conditions started when the operator first touched any piece of equipment and ended when the biosafety cabinet sash was finally shut after cleanup. Durations between these time points were grouped into four categories of activities, as enumerated below:

“Setup / annotation” includes loading and arranging labware in the cabinet; inspecting the plate and planning split factors (“Manual” condition); and all steps to set up the robot, Cell Counting Imager and software for the run (“ACCS” condition). “Teardown / cleanup” encompasses all tasks following the final liquid transfers into the destination plate, including the final inspection of the plate in the case of the “Manual” condition. For the “Manual” condition only, “Pipette tip arraying” represents the process of pre-arranging pipette tips in racks to facilitate multichannel pipetting, and “Manual passaging” encompasses the pipetting and other manual steps following tip arraying and prior to final inspection of the plate.

#### Procedure for CCI counting performance measurements

For the CCI counting characterization experiment, a master stock of HEK293T cells was harvested from a flask and cell concentration was measured using a Countess II automated cell counter. This stock was then used to produce a set of test samples of 15 mL each with nominal concentrations linearly covering a range from 100k to 400k cells/mL, along with a negative control sample consisting of pure media. From each stock a 2 mL aliquot was separated for measurement by the flow cytometer and the remainder was loaded into a well in a 12-well reservoir for measurement by the CCI.

For the flow cytometry measurements, 100 µL of Count-bright Absolute Counting Beads was added to each sample, then the samples were distributed into columns of 8 × 250 µL on a U-bottom plate (Corning, 3799) for loading by the High-Throughput Sampler of a BD FACSymphony A1 flow cytometer. From the first 3 rows measurements were taken with a sample volume of 150 µL at a rate of 3.0 µL/s.

For the CCI measurements, a purpose-made ACCS protocol script was used to cycle through the samples a total of 4 times, mixing each well and then loading the CCI, to produce 32 (8 flow cell channels × 4 repeats) measurements for each sample. Results are summarized in Fig. 2(A).

#### Procedure for flow cytometry-based seeding accuracy tests

ACCS was used to passage cells from a Challenge Plate, using CCI measurements to control seeding density, into a U-bottom plate for counting by flow cytometry, with a target of 12.5k cells/well. The ACCS run was configured as normal except that Gibco Anti-Clumping Agent (00-10057AE) was added at 1% v/v to the media in the reservoir in an effort to minimize cell aggregation and adhesion. An example of the distribution of cell concentration values reported by the CCI during this process is illustrated in Fig. 2(C)(a). Upon completion of the run, 20 µL of Countbright Absolute Counting Beads (Invitrogen, C36950) was added to each well in column 4 of the plate for QC purposes, then all wells were resuspended by triturating with a 12-channel manual pipette immediately prior to transferring to the flow cytometer. Measurement of the cell concentration in each well on the output plate was performed using a BD FACSymphony A1 flow cytometer with High-Throughput Sampler. 60 µL was sampled from each well at a rate of 3.0 µL/s – see Supplemental Materials and Methods (Supplement A) for full instrument configuration. The distribution of cell concentration values calculated from flow cytometry data for columns 1-3 and 5-12 is illustrated in Fig. 2(B). The analysis automatically excluded wells for which either the cell concentration in the source plate was too low (requiring an aliquot greater than 180 µL, the approximate practical limit we have found on the amount of cell suspension the system can reliably aspirate from the source plate without creating air bubbles), or for which the flow cytometer failed to load the sample properly (we detect this particular failure mode by an anomalous range of event timestamps in the output data).

#### Procedure for comparative seeding accuracy tests by ATP assay

For these tests the “subject” (ACCS or human volunteer) was directed to passage a provided Challenge Plate into an empty assay plate with a target of 12.5k cells/well, except for designated reference/control wells as described below. The plate was then incubated overnight and an ATP assay (CellTiter-Glo) was used to measure variation in seeding density across the plate. This measurement strategy was chosen rather than measuring concentration in suspension immediately after seeding in order to focus on the number of live cells delivered and to account for the common shortcut of dispensing the same amount of media to all wells when passaging by hand.

Each plate contained 8 “ref hi” wells and 8 “ref lo” wells seeded from prepared stocks with 20k and 10k cells, respectively, 2 “no reagent” control wells, 2 “no cells” wells and 76 sample wells for the performance measurement. For the “no cells” condition the wells were filled with 200 µL of media instead of seeding with cells. For the “no reagent” condition the wells were seeded normally but no media was removed or reagent added for the assay. Supplemental Figure S2 illustrates the layout of these assay plates.

CellTiter-Glo reagent was prepared in advance per the manufacturer’s instructions[20] and frozen in 10 mL aliquots at −20 °C. Ahead of each test, these were placed in a 37 °C bead bath just long enough to thaw, then allowed to naturally equilibrate to ambient temperature for at least 1 hour before use.

100 µL of media was removed from each well using the OT-2, excluding wells C3 and F7 by removing the corresponding tips from the tiprack, then 100 µL of CellTiter-Glo reagent was gently added to the plate manually, working column-wise and again excluding C3 and F7. A transparent plate seal was applied and the plate was moved to the plate reader as quickly as practical.

The plate reader was programmed to first perform 2 minutes of orbital shaking at “high” intensity and then take measurements of the plate every 2 minutes for 60 minutes total. We use the measurement at the +60 minute time point for analysis as past experience with this assay has shown that results at the +10 minute time point (per the vendor’s SOP) can be more sensitive to effects of pipetting technique, physical plate handling, temperature variation, etc. Measurements were carried out in top read mode with an integration time of 500 ms and a read height of 6.51 mm.

## Discussion

### Cell viability

A key area of concern when developing the system was any potential impacts on cell viability. As illustrated in the Results section, in the present design, cells spend considerably more time in ambient atmosphere when using ACCS compared to when splitting by hand. For this reason we proactively include HEPES as a pH buffer in the growth media (see “Reagents and cultures“).

In our routine usage of the system with HEK293T cells we have not observed any obvious defects in morphology or growth attributable to this difference in environmental exposure. In an informal blind comparison where lab members and colleagues viewed random pairs of microscopy images of cells from one plate seeded using ACCS and a second plate seeded by hand, neither set was consistently identified as subjectively better or worse in terms of morphological features. The assay results used to compare seeding accuracy (see Fig. 2(D)) suggest that that there is no significant deficit in overnight recovery and growth for cells passaged by the robot. In Supplement E we explore viability over longer culture times and observe no detriment attributable to robotic passaging.

### Other use cases and cell types

While our routine usage of the system has been limited to HEK293 cells thus far, we approached the system design with consideration of potential future application to other cell types. Broadly, the current embodiment is suited to adherent or suspension cells cultured in liquid media which can tolerate being outside of the incubator for 2 to 3 hours. The handling steps in each protocol are driven by an extensive set of configuration values (for example, trypsin volume, incubation temperature, number of mixing cycles, etc.) which can be adjusted for best results with a given cell type. Image processing parameters for the CCI are similarly configurable. A single exploratory test with HuH-7 cells showed that they could be counted by the CCI and survived automated passaging using standard settings, but further work to qualify and optimize the system for this application remains to be done.

### Practical impact

Although this article has focused primarily on potential gains in efficiency and throughput, increased automation in cell culture has occupational health implications as well. Intensive manual pipetting work has long been established as a significant risk factor for work-related musculoskeletal disorders (WMSD) in laboratory workers[21–23]. Further compounding the issue, most cell culture operations are carried out in a hood or biosafety cabinet; this imposes additional constraints on users’ posture and movements and is itself implicated as a risk factor for WMSD[24]. There is therefore an obvious benefit to handing off the most repetitive pipetting tasks in cell culture work to robotics.

ACCS is part of a growing field of open-source, “DIY” automation solutions[13, 25] aimed at bringing throughput gains to biology research in a collaborative, accessible and adaptable way, in contrast to high-end proprietary solutions offered by industry. For maximum reach, we have attempted to minimize hardware complexity as much as possible. The cost of the ACCS hardware is not trivial (at time of writing, the list price for the OT-2 and pipettes is $18,250[26]), but an automated liquid handler would have a great deal of practical value in the lab aside from running ACCS protocols, so the cost can be amortized over many different applications (alternatively, emerging DIY solutions for automated pipetting[14] could provide a future path to reduce the overall hardware cost). The CCI flow cell can be reused for at least several weeks and tens of cycles, and can be fabricated for under $50 per unit. The system can also be used without a CCI for passaging at fixed split ratios.

We find that training new operators can typically be done in one or two hands-on practice runs. Upkeep primarily consists of the normal maintenance requirements for the Opentrons equipment, occasional fabrication of a new CCI flow cell and waste trough, and any spot cleaning needed due to spills.

In our own usage of the system we find that not only does does the automated method offer a significant benefit in time savings, the consistency and parametric control it enables is instrumental in optimizing parameters for sensitive downstream applications. By making the system open-source we hope to share these potential benefits as well as promote further interest and collaboration in small-scale “DIY” automation solutions not only in cell culture but more broadly in life sciences research.

## Supporting information

B. ACCS Integrator's Manual

C. ACCS Cell Counting Imager Technical Manual

D. Sample CCI Normalization SOP

E. Growth and Viability

## Acknowledgments

The authors thank Diane Wiener for feedback on the manuscript, and Opentrons for technical assistance during early development and for their commitment to a fully open-source software stack for the OT-2, which was crucial to the implementation of ACCS.

## Supplementary Material

The following supplementary PDF documents accompany this article:

A. Supplemental Materials and Methods, incl. Figures S1-S2
B. ACCS Integrator’s Manual
C. ACCS Cell Counting Imager Technical Manual
D. Sample SOP for “CCI Normalization” protocol
E. Further investigation of effects on growth and viability

## Data availability

All raw data and analysis code underlying the findings and figures presented here, source code for the ACCS software components and links to documentation and CAD models can be found at https://github.com/czbiohub-sf/2024-accs-pub.

## References

1. Gil Topman, Orna Sharabani-Yosef, and Amit Gefen. A method for quick, low-cost automated confluency measurements. Microscopy and Microanalysis, 17(6):915–922, 2011. ISSN 1431-9276, 1435-8115. doi: 10.1017/S1431927611012153.

2. Thrive Bioscience. Variability in cell confluency: Comparison of human and CellAssist assessments, 2020. URL https://www1.thrivebio.com/wp-content/uploads/2020/12/Cell-Confluence-Assessment.pdf. Online; accessed 20 March 2024.

3. Marc Hafner, Mario Niepel, and Peter K Sorger. Alternative drug sensitivity metrics improve preclinical cancer pharmacogenomics. Nat. Biotechnol., 35(6):500–502, June 2017. doi: 10.1038/nbt.3882.

4. Nathan H. Cho, Keith C. Cheveralls, Andreas-David Brunner, Kibeom Kim, André C. Michaelis, Preethi Raghavan, Hirofumi Kobayashi, Laura Savy, Jason Y. Li, Hera Canaj, James Y. S. Kim, Edna M. Stewart, Christian Gnann, Frank McCarthy, Joana P. Cabrera, Rachel M. Brunetti, Bryant B. Chhun, Greg Dingle, Marco Y. Hein, Bo Huang, Shalin B. Mehta, Jonathan S. Weissman, Rafael Gómez-Sjöberg, Daniel N. Itzhak, Loïc A. Royer, Matthias Mann, and Manuel D. Leonetti. OpenCell: Endogenous tagging for the cartography of human cellular organization. Science, 375 (6585):eabi6983, 2022. doi: 10.1126/science.abi6983.

5. Mark-Anthony Bray, Shantanu Singh, Han Han, Chadwick T Davis, Blake Borgeson, Cathy Hartland, Maria Kost-Alimova, Sigrun M Gustafsdottir, Christopher C Gibson, and Anne E Carpenter. Cell painting, a high-content image-based assay for morphological profiling using multiplexed fluorescent dyes. Nat. Protoc., 11(9):1757–1774, September 2016. doi: 10.1038/nprot.2016.105.

6. M Kempner. A review of cell culture automation. J. Lab. Autom., 7(2):56–62, April 2002. doi: 10.1016/S1535-5535-04-00183-2.

7. Benjamin W. Gregor, Mackenzie E. Coston, Ellen M. Adams, Joy Arakaki, Antoine Borensztejn, Thao P. Do, Margaret A. Fuqua, Amanda Haupt, Melissa C. Hendershott, Winnie Leung, Irina A. Mueller, Aditya Nath, Angelique M. Nelson, Susanne M. Rafelski, Emmanuel E. Sanchez, Madison J. Swain-Bowden, W. Joyce Tang, Derek J. Thirstrup, Winfried Wiegraebe, Brian P. Whitney, Calysta Yan, Ruwanthi N. Gunawardane, and Nathalie Gaudreault. Automated human induced pluripotent stem cell culture and sample preparation for 3D live-cell microscopy. Nature Protocols, 19(2):565–594, 2024. ISSN 1754-2189, 1750-2799. doi: 10.1038/s41596-023-00912-w.

8. Alexandra Louey, Damián Hernández, Alice Pébay, and Maciej Daniszewski. Automation of organoid cultures: Current protocols and applications. SLAS Discov., 26(9):1138–1147, October 2021. doi: 10.1177/24725552211024547.

9. Julia Tischler, Zoe Swank, Hao-An Hsiung, Stefano Vianello, Matthias P Lutolf, and Sebastian J Maerkl. An automated do-it-yourself system for dynamic stem cell and organoid culture in standard multi-well plates. Cell Rep. Methods, 2(7):100244, July 2022. doi: 10.1016/j.crmeth.2022.100244.

10. Sebastian Eggert, Pawel Mieszczanek, Christoph Meinert, and Dietmar W Hutmacher. Open-Workstation: A modular open-source technology for automated in vitro workflows. HardwareX, 8 (e00152):e00152, October 2020. doi: 10.5281/zenodo.3986643.

11. Kevin Angers, Kourosh Darvish, Naruki Yoshikawa, Sargol Okhovatian, Dawn Bannerman, Ilya Yakavets, Florian Shkurti, Alán Aspuru-Guzik, and Milica Radisic. RoboCulture: A robotics platform for automated biological experimentation. May 2025. doi: 10.48550/arXiv.2505.14941. URL https://arxiv.org/abs/2505.14941.

12. Rick P Wierenga, Stefan M Golas, Wilson Ho, Connor W Coley, and Kevin M Esvelt. PyLabRobot: An open-source, hardware-agnostic interface for liquid-handling robots and accessories. Device, 1 (4):100111, October 2023. doi: 10.1016/j.device.2023.100111.

13. Philip Dettinger, Tobias Kull, Geethika Arekatla, Nouraiz Ahmed, Yang Zhang, Florin Schneiter, Arne Wehling, Daniel Schirmacher, Shunsuke Kawamura, Dirk Loeffler, and Timm Schroeder. Open-source personal pipetting robots with live-cell incubation and microscopy compatibility. Nature Communications, 13(1):2999, May 2022. ISSN 2041-1723. doi: 10.1038/s41467-022-30643-7.

14. Dulguunnaran Naranbat, Benjamin Phelps, John Murphy, and Anubhav Tripathi. How to convert a 3d printer to a personal automated liquid handler for life science workflows. SLAS Technology, 30: 100239, 2025. ISSN 2472-6303. doi: 10.1016/j.slast.2024.100239.

15. A. Riveiro, T. Abalde, P. Pou, R. Soto, J. del Val, R. Comesaña, A. Badaoui, M. Boutinguiza, and J. Pou. Influence of laser texturing on the wettability of PTFE. Applied Surface Science, 515: 145984, 2020. ISSN 0169-4332. doi: 10.1016/j.apsusc.2020.145984.

16. Alice Blumlein, Noel Williams, and Jennifer J. McManus. The mechanical properties of individual cell spheroids. Scientific Reports, 7(2045-2322):7346, 2022. doi: 10.1038/s41598-017-07813-5.

17. Stefanie Terstegge, Iris Laufenberg, Jörg Pochert, Sabine Schenk, Joseph Itskovitz-Eldor, Elmar Endl, and Oliver Brüstle. Automated maintenance of embryonic stem cell cultures. Biotechnology and Bioengineering, 96(1):195–201, 2007. ISSN 0006-3592, 1097-0290. doi: 10.1002/bit.21061.

18. Scott White, John Quinn, Jennifer Enzor, Janet Staats, Sarah M. Mosier, James Almarode, Thomas N. Denny, Kent J. Weinhold, Guido Ferrari, and Cliburn Chan. FlowKit: A python toolkit for integrated manual and automated cytometry analysis workflows. Frontiers in Immunology, 12: 768541, 2021. ISSN 1664-3224. doi: 10.3389/fimmu.2021.768541.

19. Thomas A Caswell, Elliott Sales De Andrade, Antony Lee, Michael Droettboom, Tim Hoffmann, Jody Klymak, John Hunter, Eric Firing, David Stansby, Nelle Varoquaux, Jens Hedegaard Nielsen, Oscar Gustafsson, Benjamin Root, Ryan May, Kyle Sunden, Phil Elson, Jouni K. Seppänen, Jae-Joon Lee, Darren Dale, Hannah, Damon McDougall, Andrew Straw, Paul Hobson, Greg Lucas, Ruth Comer, Christoph Gohlke, Adrien F. Vincent, Tony S Yu, Eric Ma, and Steven Silvester. matplotlib/matplotlib: REL: v3.7.3, 2023-09-12.

20. Promega Inc. CellTiter-Glo luminescent cell viability assay technical bulletin, 2023. URL https://www.promega.com/-/media/files/resources/protocols/technical-bulletins/0/celltiter-glo-luminescent-cell-viability-assay-protocol.pdf. Literature # TB288. Online; accessed 21 December 2023.

21. Marianne Gerner Björksten, Bo Almby, and Ejvor Sassarinis Jansson. Hand and shoulder ailments among laboratory technicians using modern plunger-operated pipettes. Applied Ergonomics, 25 (2):88–94, 1994. ISSN 00036870. doi: 10.1016/0003-6870(94)90069-8.

22. G. David and P. Buckle. A questionnaire survey of the ergonomic problems associated with pipettes and their usage with specific reference to work-related upper limb disorders. Applied Ergonomics, 28(4):257–262, 1997. ISSN 00036870. doi: 10.1016/S0003-6870(97)00002-1.

23. Jonas Winkel Holm, Ole Steen Mortensen, and Finn Gyntelberg. Upper limb disorders among biomedical laboratory workers using pipettes. Cogent Medicine, 3(1), 2016. ISSN 2331-205X. doi: 10.1080/2331205X.2016.1256849.

24. Shreya Maulik, Amitabha De, and Rauf Iqbal. Work related musculoskeletal disorders among medical laboratory technicians. In 2012 Southeast Asian Network of Ergonomics Societies Conference (SEANES), pages 1–6, Langkawi, Kedah, 2012. IEEE. ISBN 978-1-4673-1734-4 978-1-4673-1732-0 978-1-4673-1733-7. doi: 10.1109/SEANES.2012.6299585.

25. Zachary J. Heins, Christopher P. Mancuso, Szilvia Kiriakov, Brandon G. Wong, Caleb J. Bashor, and Ahmad S. Khalil. Designing automated, high-throughput, continuous cell growth experiments using eVOLVER. Journal of Visualized Experiments, (147):59652, 2019. ISSN 1940-087X. doi: 10.3791/59652.

26. Opentrons Labworks Inc. OT-2 Robot, 2024. URL https://opentrons.com/products/ot-2-robot?sku=999-00111,%20999-00003,%20999-00006. Online; accessed 17 November 2024.

