## Supplementary material for "Open-source cell culture automation system with integrated cell counting for passaging microplate cultures": A. Supplementary Materials & Methods

### Supplemental Materials and Methods

#### Contents

ACCS passaging protocol
Flow cytometer settings
Supplemental figures

#### ACCS passaging protocol

This section expands on the steps of the automated passaging process in more detail than could be accommodated in the main article.

Growth media and dissociation reagent are warmed to 37°C prior to loading on the robot; any other reagents are loaded at room temperature. HEPES is used in the media as a precaution to ensure stable pH since the plates spend more time out of the $CO_2$ incubator atmosphere than they typically would when passaging by hand. See the Reagents and Cultures subsection of the main article for a full enumeration of the reagents used for ACCS protocols.

Harvesting of cells from the source plate is carried out columnwise with the 8-channel pipette. The harvesting phase consists of aspirating and discarding the old media, gently washing the adhered cells with 100 uL of DPBS, adding 60 µL of dissociation reagent, incubating for approximately 8 minutes, quenching with 200 µL of growth media to neutralize trypsin, then mixing vigorously to dissociate the cells into singlets as effectively as possible. From the start of the protocol, the block heater is held at a setpoint of 41°C which we have empirically found to maintain the bottom of the plate as close as practical to 37°C; after all wells have been quenched the block is allowed to coast back to room temperature.

Following the dissociation step, if seeding according to cell density targets, the CCI is used to measure the concentration of harvested cells in each well. The process consists of taking a 30 µL sample from a column using the 8-channel pipette, injecting the samples into the CCI, waiting for 1 minute for the cells to settle in the chambers, then running a CCI counting cycle. The CCI takes about 100 seconds to acquire the images and perform analysis for all 8 channels, then the channels are immediately flushed with DPBS. In the case of a "blind split", the CCI is not involved, no sample is taken for counting, and seeding volumes are calculated directly from dilution ratios specified by the user.

The cell concentration values returned by the CCI software are used to calculate the appropriate volume to transfer for each well. Seeding volumes are constrained to a range of 20 µL to 200 µL, dictated at the upper end by the amount of cell suspension that can be reliably recovered from the source well and at the lower end by the volume range rating of the pipette. If the calculated volume is outside this range, a warning is issued and the value is clipped to the top or bottom of the range as appropriate. Practically speaking we find that aliquots are generally reliable up to 180 µL, above which there is greater likelihood of air bubbles being drawn into the pipette tip.

Seeding of the output plates is carried out with the single-channel pipette. For each well, first the appropriate quantity of media for dilution is added to the output well, then the cell suspension in the source well is mixed once more to ensure homogeneity before finally transferring the desired number of cells to the output well.

When seeding a duplicate plate, each column of the first plate is seeded with the sum of the cells and media for both copies, then the 8-channel pipette is used to mix the contents and transfer a portion of the total mixture to the duplicate. The amount transferred is controlled by a user-specified ratio, typically between 1:3 and 1:1, allowing the generation of, for example, a stock plate seeded at 25k cells/well and an imaging plate seeded at 12.5k cells/well.

Throughout all steps, the use of pipette tips is managed in such a way as to avoid cross-contamination of samples. Because the liquid volumes for adding trypsin and media to the source plate during harvesting are not especially critical, these dispensing steps are accomplished with the pipette tips hovering above the wells so as to avoid contact with cells and allow a single set of tips to be used on a whole 4-column group. For all steps involving contact with cell suspension, tips are discarded and replaced between plate locations. During seeding of the output plate(s), any media needed for dilution is transferred first, then the same tip is reused for transferring cell suspension.

*ACCS: Open-source cell culture automation system with integrated cell counting for passaging microplate cultures*

### Flow cytometer settings

This section lists the settings we used on the BD FACSymphony A1 when using it as a high throughput cell counter for ACCS seeding performance tests.

| Cytometer parameter | Value |
|---|---|
| Trigger logic | OR |
| Window extension | 0.00 µs |

| High Throughput Sampler parameter | Value | | Detector | Recording channels | Voltage | Trigger Threshold |
|---|---|---|---|---|---|---|
| Loader mode | Standard | | FSC | FSC-H, FSC-W, FSC-A | 350 V | 30000 |
| Sample volume | 60 µL | | SSC | SSC-H, SSC-W, SSC-A | 225 V | |
| Injection rate | 3.0 µL/s | | BB515 | BB515-H | 210 V | 1000 |
| Mixing volume | 75 µL | | BB700 | BB700-H | 285 V | |
| Mixing speed | 180 µL/s | | | | | |
| # of mixes | 5 | | | | | |
| Wash volume | 800 µL | | | | | |

### Supplemental figures

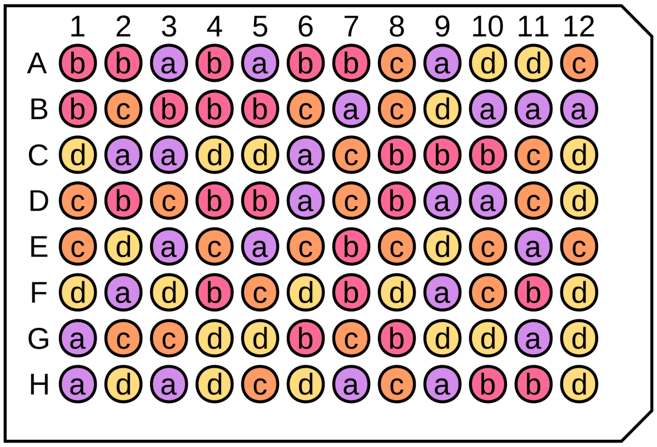


**Supplemental Figure S1. Challenge Plate layout.**
Four groups of 24 wells are randomly distributed on the plate. Each group is assigned a relative seeding density, ranging linearly from 50% for "a" wells to 100% for "d" wells.

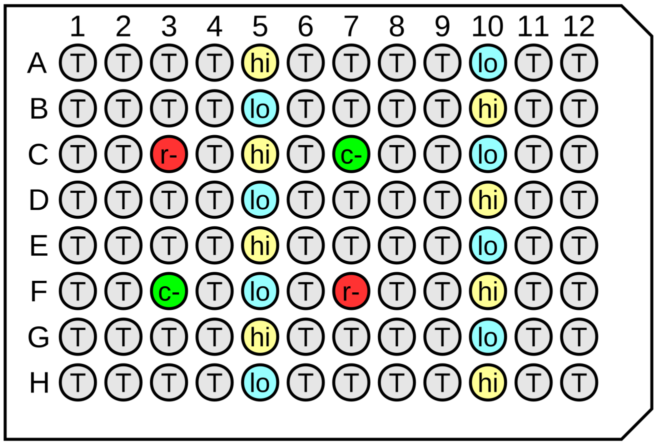


**Supplemental Figure S2. CellTiter-Glo assay plate layout.**
Wells marked "r-" and "c-" are "no reagent" and "no cells" negative controls, respectively. Wells marked "hi" and "lo" are seeded from prepared stocks with 20k and 10k cells, respectively. The 76 remaining wells marked "T" are the sample set for the test.
