## Supplementary material for "Open-source cell culture automation system with integrated cell counting for passaging microplate cultures": B. ACCS Integrator's Manual

241217a

### ACCS Integrator's Manual


Introduction

Hardware components

- Liquid handler robot and accessories
- Operator PC
- Cell Counting Imager (CCI)
- CCI cover
- Tip waste bin (3-0390) and liner
- Tilted heat block adapter (3-0271)
- Tilted plate calibration tool (3-0938)
- Reservoir riser (3-1482)
- Inspection mirror and rear view mirror
- Other tools and facilities

OT-2 setup and modifications

- Pipette modules
- Side / top panels
- Tip waste bin removal
- Ethernet connection
- Software environment

Hardware installation

- Deck layout
- Installation in a biosafety cabinet
- Power and electronic connections

PC software setup

Equipment list

- Core off-the-shelf equipment
- Tools
- Custom components
- Other

Consumables

- Lab plastics
- Cleaning and disposal supplies
- Reagents

**241217a**

### Introduction

This document is designed to provide practical guidance to someone considering implementing ACCS for their own lab. It assumes a basic level of familiarity with hardware prototyping, computer software and wet lab practice.

### Hardware components

#### Liquid handler robot and accessories

ACCS is built around an Opentrons OT-2 liquid handling robot with the following Opentrons accessory equipment:

- 8-channel P300 Gen2 pipette -- used for columnwise processing steps on the source plate (media removal, washing, dissociation) and for loading and flushing the CCI
- 1-channel P300 Gen2 pipette -- used for seeding destination plate
- Temperature Module Gen2 -- temperature-controlled heating/cooling block, used to incubate the source plate prior to and during trypsinization

#### Operator PC

The only strict hardware requirement is that a dedicated gigabit ethernet port must be available for the CCI camera.

We use a Windows 10 system in production. The ACCS and CCI software and their dependencies are cross-platform so in principle a Linux or macOS system could be used instead.

Note that the CCI software as-supplied is designed with the assumption of one CCI per PC, so some slight modifications would be required to operate multiple CCIs simultaneously.

**241217a**

### Cell Counting Imager (CCI)

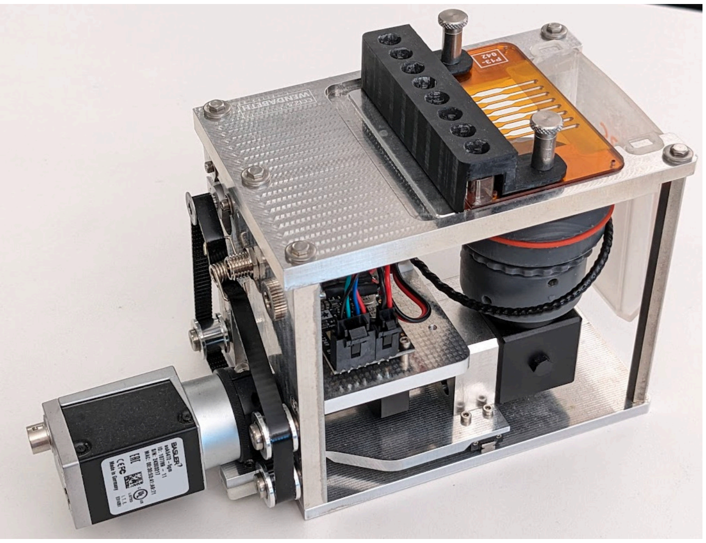

Refer to the separate *ACCS Cell Counting Imager Technical Manual* for information on the instrument itself, the flow cell, and support equipment and consumables related to CCI operation.

### CCI cover

An improvised cover for the CCI can be made by cutting notches out of the corners of an Opentrons tipbox lid to clear the CCI fastener heads, as shown below.

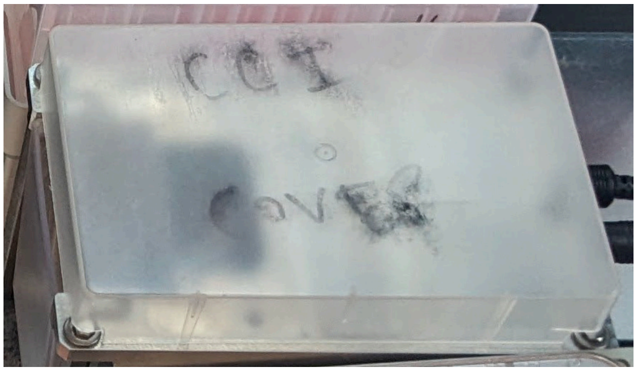


The CCI should be covered when spraying down the deck for disinfection or when the system is not in use, as the objective faces upward.

### Tip waste bin (3-0390) and liner

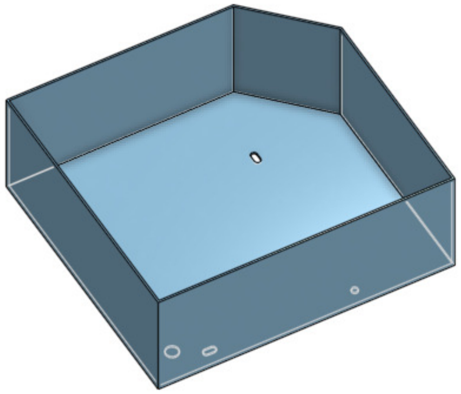

The tip waste bin replaces the small bin built into the OT-2. The tip waste bin is made of 3D printed PLA plastic. The material used for the bin is not critical.

The bin is used with a liner made by cutting a 19" x 23" biohazard bag (Heathrow Scientific HS10322) down to about 10" height. The bin liner is installed by pushing the bottom of the bag into the waste bin, then folding the excess material over the edges and securing with stainless steel binder clips (e.g. McMaster-Carr 12755T81) so that the overhanging material sits flat against the sides of the bin on the two long sides. **It is important to avoid bunching of material on the floor of the bin as much as possible** to reduce the tendency for tips to form tall piles in the drop area. It is also advisable to check on the bin at least once during a long protocol and break up tip pileups if they form.

The bin model includes fastener holes to secure it to the OT-2 deck but in practice we find it more practical to leave it loose so that it can be freely handled to install the liner. The bin is installed by simply placing it on the robot deck as shown:

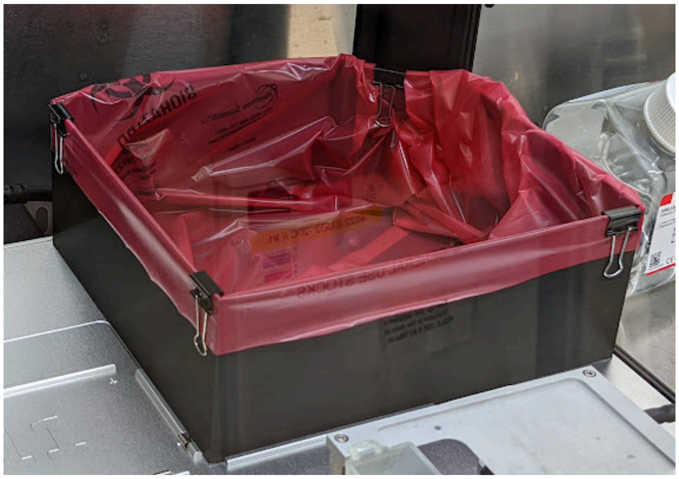

To accommodate the custom bin, the OT-2's built-in bin assembly must be removed as described in Tip waste bin removal.

An alternative single-use tip waste bin design made from laminated cardstock and featuring an inclined bottom is being considered to simplify setup and improve reliability.

#### Tilted heat block adapter (3-0271)

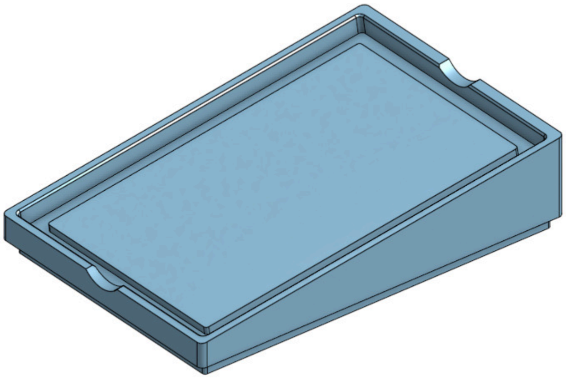

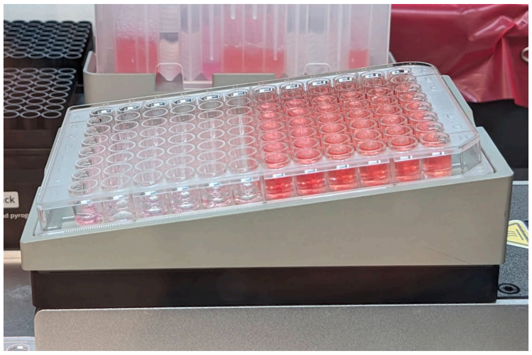

The tilted heat block adapter holds the source plate at an 8° incline while maintaining thermal contact between it and the Temperature Module. Tilting the plate is necessary in order to allow the pipettes to access the full volume of liquid in the wells. The narrow aspect ratio of the wells on a 96 well plate gives a generous allowance for tilt compared to plate types with wider wells which typically must be laid flat between pipetting steps to prevent the well bottoms being uncovered.

We find that a setpoint of 41°C on the Temperature Module results in peak well bottom temperatures just below 37°C and an acceptably uniform temperature (within 2°C) across the plate.

Because the Opentrons software does not currently support the concept of labware being mounted in a non-flat orientation, a special "tilted plate" labware definition file is used. A special tool to aid with labware position check / pipette calibration because the typical strategy of visually lining up the pipette tips to the rims of wells is not reliable.

The tilted heat block adapter is machined from 6061 aluminum and is bead-blasted followed by a Type II anodized finish. The bead blasting is not essential provided that sharp edges are broken by other means but anodizing is recommended for corrosion resistance.

The adapter is installed by simply placing it on the bare platen of the Temperature Module (ensure any included adapter is removed first, revealing the black surface) and is held in place by gravity. Finger notches are provided to make it easier to remove the microwell plate from the adapter.

The adapter is designed so that common plastic 96-well culture plates should drop in without resistance. Our testing was based on Thermo Scientific BioLite plates, cat. no. 130188.

#### Tilted plate calibration tool (3-0938)

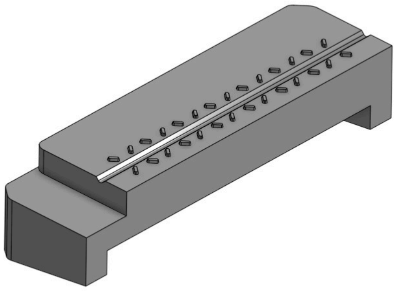

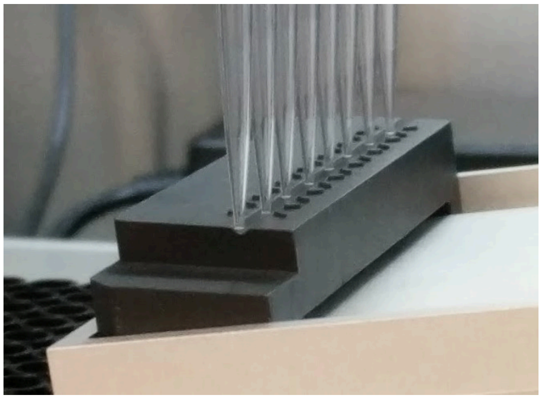

The tilted plate calibration tool is used to achieve non-contact alignment of the multichannel pipette with its calibration position during the Opentrons labware position check / calibration process. It was printed on a Formlabs Form 3 SLA printer using Formlabs Black Resin V4.

It is important that the tool sit in a consistent, stable and level position when placed on the tilted block; depending on the printing configuration, this may require sanding or machining the bottom of the tool. The height of the tool can be adjusted by shimming with Kapton tape or a similar material. The height should be set such that the tips just nearly reach (but never push against) the bottom of the source plate wells after calibration using the tool. A calibration position that is too high may result in under-aliquoting of cells and/or more frequent introduction of air bubbles during mixing. A calibration position that is too low may result in some or all tips being blocked due to being held against the well bottoms, preventing proper aspiration.

Routine usage of the calibration tool is detailed in the *CCI Normalization Sample SOP*.

#### Reservoir riser (3-1482)

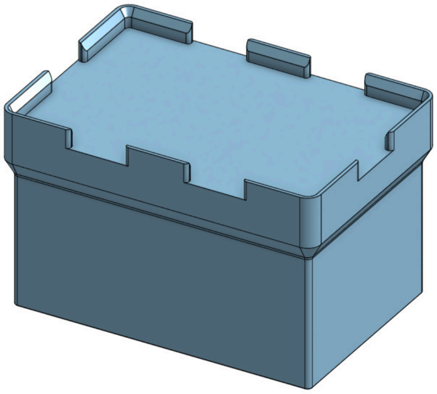

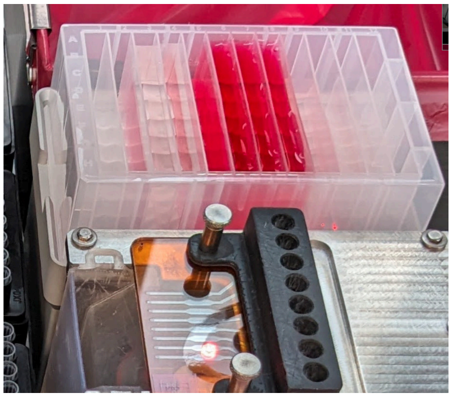

The reservoir riser allows drop-in installation of the reagent reservoir and elevates it by 76.5mm to make it easier for the operator to access it. It is machined from 6061 aluminum. Refer to finishing notes for the heat block adapter.

The reservoir riser was designed to address recurring issues related to mounting the reservoir directly on the deck. In particular, the design of the rails and spring clips on the OT-2's deck allows false placement of the reservoir such that it appears properly installed at a glance and feels secure, but is in fact out of position, leading to crashes and potentially catastrophic outcomes (such as the reservoir being picked up by the pipette and driven into the CCI). The reservoir riser is not an essential component but is recommended for these reasons. Although the solid metal construction is preferred for stability and hygiene reasons, 3D printing this part is a viable lower-cost alternative.

#### Inspection mirror and rear view mirror

Similarly to the reagent reservoir, the Opentrons tip boxes can inadvertently be installed out of position. This can be avoided by ensuring the rails are visible on all sides of each box, but this can be difficult when the OT-2 is installed in a recessed location such as inside a biosafety cabinet.

An inspection mirror with a telescopic handle (e.g. McMaster-Carr 1017T25) is useful for verifying that tipracks are seated properly. Additionally, we have installed a convex mirror (United Pacific 43001) at the back of the robot to provide a rear view of the deck as shown below. The improvised mounting solution consists of a modified piece of aluminum angle attached to the robot using a set of small panel clamps (AES Industries AES-20410). The mirror is positioned such that the bottoms of the rearmost tipboxes are visible from the operator's normal seated eye level during setup.

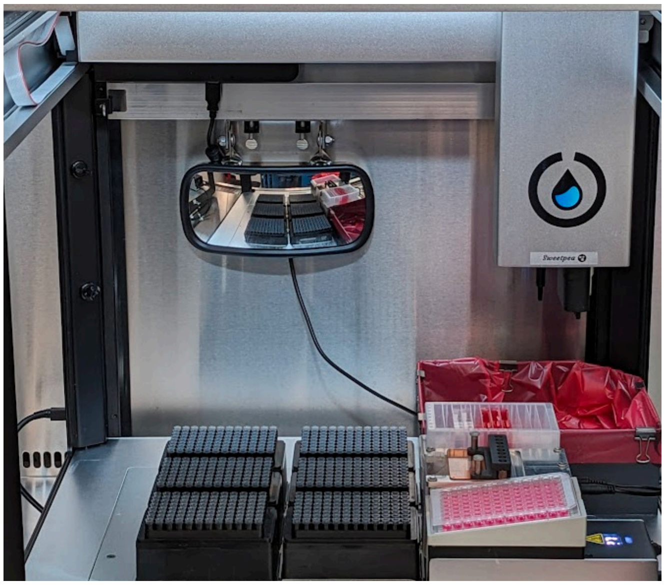

#### Other tools and facilities

It is recommended to keep a multi- and a single-channel manually operated P300 pipette that fit universal tips (allowing use with the Opentrons tips) with the system.

In addition, particularly if the system is installed in a BSC, it is very helpful to have a suction line and waste trap available for emptying the CCI waste cup and reagent reservoir in-place.

For clearing liquid from the CCI flow cell (see *CCI Normalization Sample SOP*), we currently use a pneumatic blow gun (McMaster-Carr 5186K81) with a non-filter 300uL pipette tip attached using tape. The blow gun is supplied by house compressed air via a regulator and particulate filter; pressure is set to ~0.2 bar. This step could also be performed with a manual pipette with potentially lower aerosol generation risk but we have found the compressed air method to yield the most consistent results in terms of the flow cell being properly primed by the robot.

### OT-2 setup and modifications

#### Pipette modules

Pipette modules should be installed as follows, following the Opentrons documentation:

- Left mount: P300 single-channel Gen2
- Right mount: P300 8-channel Gen2

#### Side / top panels

Assuming the system is to be used in a biosafety cabinet, the clear side and top panels should be removed from the robot to allow for free air flow.

#### Tip waste bin removal

To accommodate the custom tip waste bin, the built-in bin assembly must be removed from the OT-2. To do this, first take the black plastic insert and detach the bin frame by removing four screws in the locations indicated below:

241217a

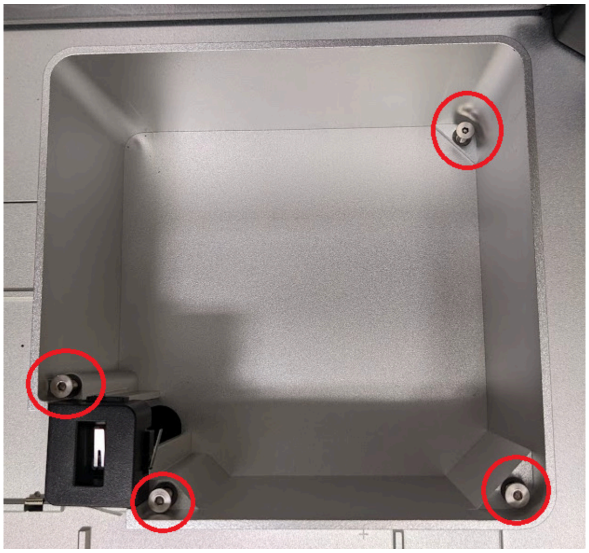

With the bin frame loose, lift it up and disconnect the electrical connector before fully removing it from the deck. Feed the end of the cable down through the passthrough hole and tape it to the underside of the deck to keep it out of the way.

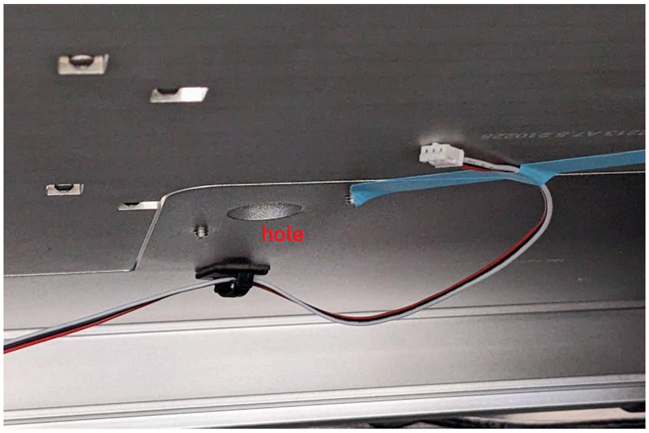


#### Ethernet connection

Operation of the CCI relies on a consistent network connection between the OT-2 and the PC running the CCI software. Ideally the PC and robot(s) in use should share an ethernet network and each have fixed IP addresses.

The version of the robot currently sold (sometimes called OT-2R) has an ethernet connection available and can be used as-is. Early versions of the OT-2 provided a USB port as the primary hardwired connection for controlling the robot. Internally this is wired to a USB ethernet adapter which in turn is connected to the Raspberry Pi that runs the robot. This can be removed to use a direct ethernet connection instead.

Remove the left hand frame panel from the OT-2 to expose the compartment with the ethernet and power connections in it. Unplug the USB-ethernet adapter from the PC board and the internal ethernet patch cable. Thread a 1.5ft male-female ethernet extension cable (e.g. Cable Matters 160024-BLK-1.5) through the bottom of the robot frame into the compartment and connect it to the internal patch cable with a female-female coupler (e.g. Monoprice 107297).

Ready-to-use ethernet jack on OT-2R:

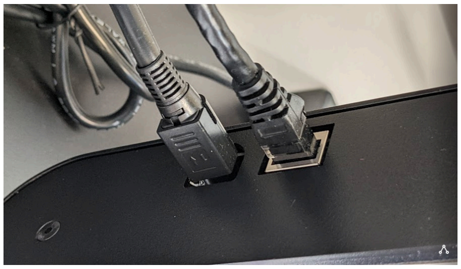

Old style OT-2 with USB ethernet adapter:

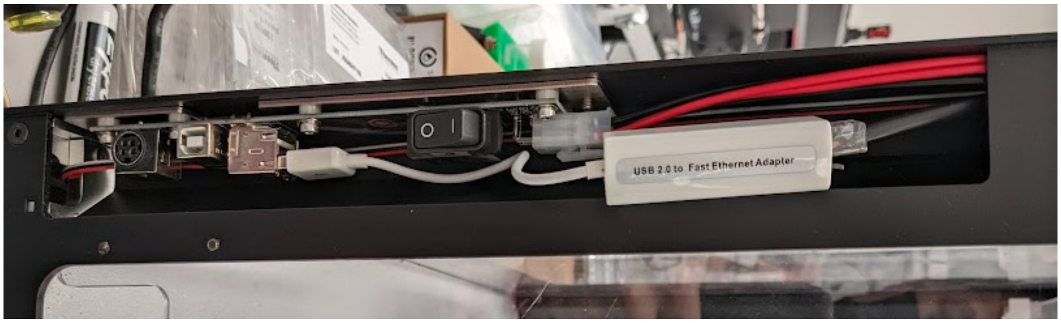


USB ethernet adapter removed and replaced with coupler and extension:

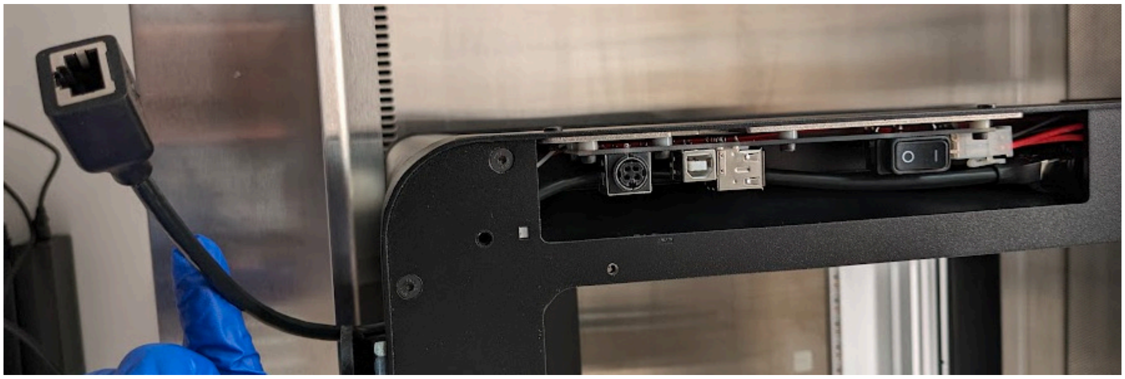

**241217a**

#### Software environment

The software environment on the robot is based on Opentrons OT-2 4.7.0 software image. The system has not yet been tested and validated with later Opentrons software versions. A new OT-2 must be downgraded by following the instructions on the Opentrons site.

The Opentrons app and matching robot software image version 4.7.0 can be found here: https://github.com/Opentrons/opentrons/releases/tag/v4.7.0

The ACCS protocol framework repository includes `ot2logbot` as well as instructions for obtaining and installing the appropriate Opentrons software version on the OT-2 and setting it up for ACCS operations:
https://github.com/czbiohub-sf/accs-protocol-framework-pub

After installing the 4.7.0 software image, the only further change that is strictly required is to install the `requests` Python package, which is used to communicate with the CCI server. Optional components such as the ACCS Slack alert service (`ot2logbot`) can be installed by following the accompanying READMEs.

The above can be achieved using the file upload facility and terminal emulator available in the onboard Jupyter environment, accessible via a link in the Opentrons app. For more convenient access, particularly for transferring files, it is helpful to set up SSH access to the OT-2 by following the instructions supplied by Opentrons.

### Hardware installation

#### Deck layout

The photo below shows a typical layout of equipment installed on the OT-2 deck for the cci_normalization protocol.

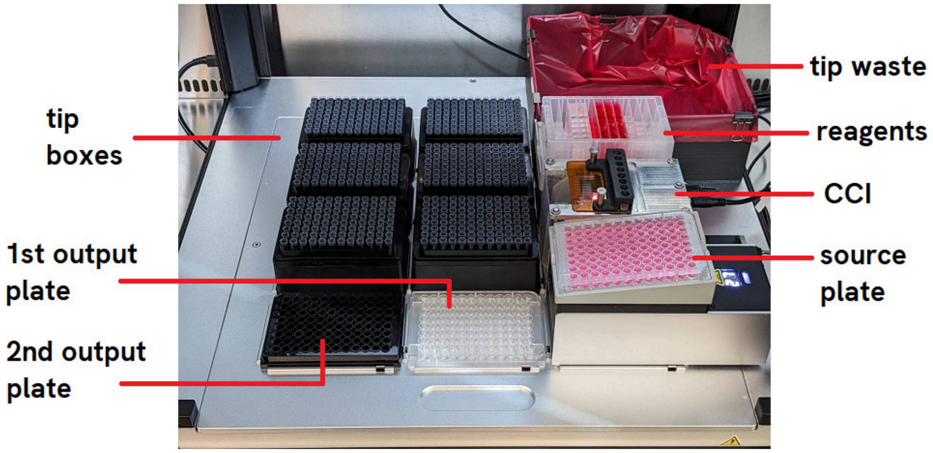


See the *CCI Normalization Sample SOP* for more details on installing items on the deck for each run.

The CCI, reservoir riser and Temperature Module are normally always left on the deck unless running protocols that require other items to be installed in their corresponding deck slots.

#### Installation in a biosafety cabinet

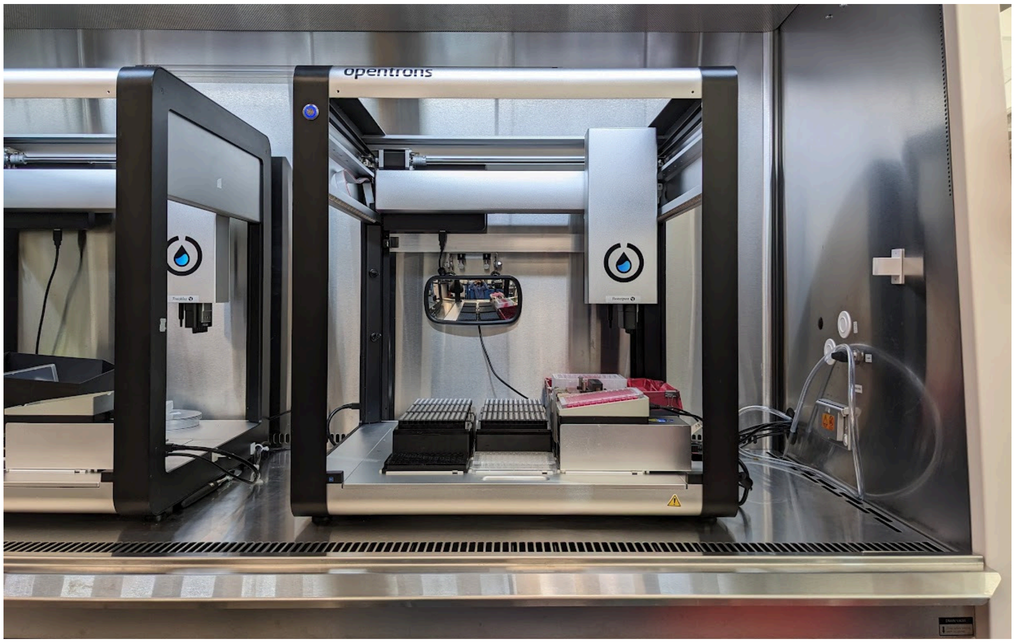


A single ACCS setup can comfortably occupy a typical 4-foot biosafety cabinet with enough workspace left on the side for manual prep tasks such as filling the reagent reservoir. Two

ACCS setups can fit side-by-side in a 6-foot cabinet, but using the same cabinet for prep work becomes less practical, especially when both robots are in use. Additionally, the work surface must be rated (or appropriately reinforced) to support the weight of the two robots and all other equipment.

Note that the OT-2 is heavy (48 kg), so manipulating it in and out of the cabinet should be done using at least two people.

Also note that reaching the back of the OT-2 deck in a BSC with the sash at normal operating height may be challenging depending on the length of the operator's arms. In this case the operator may consider temporarily opening the sash fully to pre-install the trash bin and (lidded) rear row tip boxes.

#### Power and electronic connections

The following wiring must be routed into the BSC, per setup:

- OT-2 power cable
- Temperature Module power cable
- OT-2 ethernet cable
- CCI camera ethernet cable
- CCI stage power cable
- CCI stage serial cable

Wiring and support equipment for the CCI is covered in the *ACCS Cell Counting Imager Technical Manual*.

To save space and reduce airflow obstructions in the biosafety cabinet, we locate the power supplies for the OT-2 and Temperature Module outside the cabinet. Both devices use identical 36V power supplies whose DC output leads terminate in a Kycon KPPX-4P connector. We use off-the-shelf 4ft extension cable assemblies (GlobTek KPPX4124642M0KPJX4(R)) to add the necessary length to the output leads to route them into the BSC.

**241217a**

### PC software setup

The first software to install is the Opentrons app, specifically version 4.7.0, which is available on GitHub:
https://github.com/Opentrons/opentrons/releases/tag/v4.7.0

The ACCS software suite includes the ACCS protocol framework itself, the user-facing UI for generating protocol scripts, and tools for protocol development. The latest version of the software and instructions for setting up the system can be obtained from the Github repository: https://github.com/czbiohub-sf/accs-protocol-framework-pub . This repository also includes the optional Slack alert bot that can be installed on the OT-2 (`ot2logbot`).

Refer to the *CCI Technical Manual* for the setup steps for the CCI server, camera drivers, etc.

### Equipment list

#### Core off-the-shelf equipment

- Opentrons OT-2 pipetting robot, 999-00111
- Opentrons 8-channel P300 Gen2 pipette, 999-00006
- Opentrons 1-channel P300 Gen2 pipette, 999-00003
- Opentrons Temperature Module Gen2, 991-00350-0
- Operator PC with at least 1 gigabit ethernet port

#### Tools

- Universal fit P200 or P300 single-channel manual pipette
- Universal fit P200 or P300 8-channel manual pipette
- Telescopic inspection mirror, e.g. McMaster-Carr 1017T25
- Pneumatic blow gun -- see Other tools and facilities

#### Custom components

See corresponding sections under Hardware Components for detailed information

- 3-0390 Tip waste bin
- 3-1482 Reservoir riser
- 3-0271 Tilted heat block adapter
- 3-0938 Tilted plate calibration tool
- Cell Counting Imager and accompanying components:
  *see ACCS Cell Counting Imager Technical Manual*

#### Other

- 2x DC power extension lead for OT-2 and Temperature Module, GlobTek KPPX4124642M0KPJX4(R)

### Consumables

#### Lab plastics

- Pipette tips: OT-2 200µL Filter Tips, Opentrons 999-00081
- Culture plate: BioLite 96-well plastic microwell plate, Thermo-Fisher 130188

#### Cleaning and disposal supplies

- Tip trash liner: 19" x 23" biohazard bag (modified as described in the Hardware Components section), Heathrow Scientific HS10322
- CCI flow cell external cleaning agent: Blue Ribbon Products Plexi-Clean
- Wipes for CCI flow cell external cleaning: 4x4" polyester cleanroom wipes, Texwipe Absorbond TX404

#### Reagents

Refer to the *CCI Normalization Sample SOP* for more context for the following.

- Buffer for cell wash and CCI flow cell flush: Dulbecco's Phosphate-Buffered Saline, Gibco 14190-144
- Dissociation agent: 0.25% Trypsin-EDTA, Gibco 25200-056
- Culture media (example as used for routine culture of HEK293):

- DMEM, high glucose with GlutaMAX, Gibco 10566-016
- + 10% v/v fetal bovine serum
- + 25 mM HEPES, Gibco 15630-080
- + 100 U/mL penicillin-streptomycin, Gibco 15140-122

- CCI flow cell deep clean solution: 10% v/v in DI water of Contrad 70, Fisher Scientific 04-355
- Deionized water for final CCI flow cell rinse

Note in particular the addition of HEPES to the media as a proactive measure to stabilize culture pH as plates may spend multiple hours outside of the incubator.
