## Supplementary material for "Open-source cell culture automation system with integrated cell counting for passaging microplate cultures": C. ACCS Cell Counting Imager Technical Manual

Intro, scope and first steps
Bill of Materials
- Custom machined / printed parts
- Off-the-shelf hardware

CCI instrument machined / printed parts
- Chassis parts (3-1239, 3-1240, 3-1241, 3-1242)
- Optics platform parts (3-1243, 3-1246, 3-1247, 3-1250)
- Motor pulley (3-1263)
- Pipette tip guide / flow cell clamp (3-0885)
- Darkfield illuminator parts (3-1253, 3-1254, 3-1261)
- Focus locking sleeve and focus knob (3-1252, 3-1260)
- Waste trough (3-1273)

CCI instrument hardware build guide
- Flow cell fasteners
- Belt clamp
- Focus locking sleeve
- Motor controller board
- USB serial communications cable
- Wiring harness prep
- Darkfield illuminator collar
  - Threaded insert
  - LED installation and wiring
- "Middle section" and motor test
- Linear guide mounting
- Optics platform
- Turret and camera tube assembly
- Optical system
- Drive train
- Install focuser and illuminator
- Top plate

Commissioning
- Software setup
- Basic stage function test
- Camera test
- Focus adjustment
- Top plate position adjustment
- Software stage alignment

CCI flow cell design and fabrication
- Overview
- Top (3-0352) and bottom (3-0305) plates
- Spacer layer (3-0302)
- Pipette tip interface (3-0303)
- Waste nozzle strip (3-0377)

General operating procedures
- Using the CCI
- Flow cell storage and working life
- Flow cell cleaning
- Cleaning the CCI instrument

### Intro, scope and first steps

The Cell Counting Imager is an open-source imaging-based cell counter designed to facilitate automated cell culture with a pipetting robot. It is part of a project called the Automated Cell Culture Splitter (ACCS), which uses an Opentrons OT-2 to passage (harvest and transfer a portion to a new vessel with fresh media) human cells cultured in 96-well plates. This document is one of several resources being released alongside the manuscript and its purpose is to be the primary source of information pertinent to building and operating the CCI hardware.

There is no specialized experience strictly required to build the CCI, assuming the fabrication of the custom fabricated parts is outsourced. However, it should be noted that this is an experimental, unpolished design, and someone who intends to build a CCI using this information would benefit greatly from a basic foundation of general hardware prototyping and troubleshooting experience. This document does not instruct on general skills such as soldering or splicing and terminating cables. The builder should seek out appropriate resources and assistance according to their experience level in order to complete the work in accordance with relevant best practices.

It is recommended to review the main manuscript and the ACCS Integrator's Manual in addition to this document for context. Links to the preprint, written supplements and other resources such as a CAD model of the instrument can be found here:
https://github.com/czbiohub-sf/2024-accs-pub

### Bill of Materials

This section enumerates all of the physical components and some specific tooling involved in constructing the CCI instrument. It assumes a ready supply of general electronics prototyping supplies such as hookup wire, heat shrink tubing, solder, etc. as well as access to basic soldering equipment and hand tools.

### Custom machined / printed parts

See the following section (Custom hardware components) for further detail on each part.

*Glossary: FDM = Fused Deposition Modeling (filament based 3D printer), SLA = Stereolithography (resin based 3D printer)*

| Part no | Description | Process | Material |
|---|---|---|---|
| ***Chassis*** | | | |
| 3-1239 | Frame bottom plate | CNC milling | Aluminum 7075-T651 |
| 3-1240 | Frame vertical plate | CNC milling | Aluminum 6061-T651 |
| 3-1241 | Frame top plate | CNC milling | Aluminum 6061-T651 |
| 3-1242 | Frame shelf | CNC milling | Aluminum 6061-T651 |
| ***Stage*** | | | |
| 3-1243 | Optics platform base plate | CNC milling | Aluminum 7075-T651 |
| 3-1245 | Optics platform belt clamp body | SLS printing | Glass reinforced polyamide |
| 3-1247 | Mirror cube mounting bracket | CNC milling | Aluminum 6061-T651 |
| 3-1250 | Stage homing switch flag | FDM printing | PLA |
| 3-1263 | Stage motor pulley | Manual drill & ream | McMaster # 1375K29 |
| ***Optical system*** | | | |
| 3-1246 | Optics tube spacer ring | Manual turning | McMaster # 8978K937 |
| 3-1252 | Focus locking sleeve body | SLA printing | Formlabs Tough 2000 resin |
| 3-1260 | Focus knob | SLA printing | Formlabs Tough 2000 resin |
| 3-1253 | Illuminator main body | SLA printing | Formlabs Tough 2000 resin |
| 3-1254 | Illuminator shroud, front half | SLA printing | Formlabs Tough 2000 resin |
| 3-1261 | Illuminator shroud, rear half | SLA printing | Formlabs Tough 2000 resin |
| ***Misc*** | | | |
| 3-1273 | Waste trough | SLA printing | Formlabs Clear V4 resin |
| ***Flow cell*** | | | |
| 3-0302 | Flow cell spacer layer | CO2 laser cut | (see notes) |
| 3-0303 | Flow cell pipette tip interface | SLA printing | Cast PUR |
| 3-0305 | Flow cell bottom slide | CO2 laser cut | Cell cast PMMA sheet |
| 3-0352 | Flow cell top slide | CO2 laser cut | Cell cast PMMA sheet |
| 3-0377 | Flow cell waste nozzle strip | CO2 laser etch & cut | Adhesive backed PTFE film |

### Off-the-shelf hardware

| Qty | Vendor | Part number | Description | Drive Size |
|---|---|---|---|---|
| ***Flow cell fasteners*** | | | | |
| 2 | McMaster-Carr | 91125A613 | SS round F-F standoff, 4-40, 1/4" OD | |
| 2 | McMaster-Carr | 91830A101 | SS thumbscrew, 3/8" head dia, 1/8" shldr ht, 1/4" lg | |
| 2 | McMaster | 92785A507 | SS cone point set screw, 4-40, 3/4" lg | 0.050" hex |
| ***Chassis*** | | | | |
| 2 | McMaster-Carr | 1873N51 | Nylon cable tie anchor, 0.13" tie wd, #4 screw hole | |
| 2 | McMaster-Carr | 91075A228 | SS M-F hex standoff, 4-40, 3.75" lg | 1/4" ext hex |
| 5 | McMaster-Carr | 91525A314 | SS washer, #4, 0.344" OD | |
| 2 | McMaster-Carr | 92196A107 | SS socket head screw, 4-40, 5/16" lg | 3/32" hex |
| 3 | McMaster-Carr | 92196A109 | SS socket head screw, 4-40, 7/16" lg | 3/32" hex |
| 2 | McMaster-Carr | 92314A110 | SS hex head screw, 4-40, 1/2" lg | 3/16" ext hex |
| 3 | McMaster-Carr | 92314A112 | SS hex head screw, 4-40, 5/8" lg | 3/16" ext hex |
| 2 | McMaster-Carr | 92949A106 | SS button socket head screw, 4-40, 1/4" lg | 1/16" hex |
| 2 | McMaster-Carr | 96246A094 | Heli-Coil insert, 4-40, 0.224" lg, dry film lubricated | |
| ***Stage mechanical*** | | | | |
| 1 | McMaster-Carr | 1679K626 | MXL timing belt, 1/4" wd, 135T | |
| 3 | McMaster-Carr | 3693N14 | HTD idler pulley, 15mm OD, for 5mm shaft | |
| 6 | McMaster-Carr | 91292A004 | SS socket head screw, M2x0.4mm, 4mm lg | 1.5mm hex |
| 10 | McMaster-Carr | 91292A340 | SS socket head screw, M2.6x0.45mm, 6mm l | 2mm hex |
| 2 | McMaster-Carr | 91585A093 | SS dowel pin, 1mm dia, 20mm lg | |
| 2 | McMaster-Carr | 92141A005 | SS washer, #4, 0.312" OD | |
| 1 | McMaster-Carr | 92196A022 | SS socket head screw, 4-40, 9/32" lg | 3/32" hex |
| 2 | McMaster-Carr | 92210A107 | SS countersunk socket head screw, 4-40, 5/16" lg | 1/16" hex |
| 1 | McMaster-Carr | 93615A111 | SS low profile socket head screw, 4-40, 3/8" lg | 0.050" hex |
| 2 | McMaster-Carr | 93615A317 | SS low profile socket head screw, 8-32, 1/4" lg | 5/64" hex |
| 1 | McMaster-Carr | 96246A046 | Heli-Coil insert, Nitronic 60, 4-40, 0.168" lg | |
| 3 | Misumi | DBTS3-11-6-PC | Low hd shldr screw, M3, 5mm x 11mm shldr, 6mm thd | 2mm hex |
| 1 | Misumi | SSE2B6-85-MC | Mini linear guide assy, 85mm lg, dual carriage | |
| 1 | Misumi | SSELB6-55-MC | Mini linear guide assy, 55mm lg, long carriage | |

| | | | |
|---|---|---|---|
| 3 | Misumi | WSSS8-5-1 | Cylindrical spacer, 8mm OD, 5mm ID, 1mm ht |
| 1 | Partsbuilt 3D | R2C2-SPRING-BELT | Belt tensioner spring |

*Stage electronics*

| | | | | |
|---|---|---|---|---|
| 1 | Pololu | 3138 | Tic T249 stepper motor controller | |
| 9 | TE Connectivity | 1-104480-6 | AMPMODU crimp socket contact for 22-26AWG | |
| 4 | TE Connectivity | 1-104481-3 | AMPMODU crimp socket contact for 28-32AWG | |
| 2 | TE Connectivity | 104257-1 | AMPMODU locking connector housing, 2 pos | |
| 1 | TE Connectivity | 104257-3 | AMPMODU locking connector housing, 4 pos | |
| 1 | TE Connectivity | 104257-9 | AMPMODU locking connector housing, 10 pos | |
| 2 | TE Connectivity | 5-103908-1 | Shrouded locking vertical header, 2 pos | |
| 1 | TE Connectivity | 5-103908-3 | Shrouded locking vertical header, 4 pos | |
| 1 | Tensility Intl | 54-00063 | Panel mount barrel jack, 5.5x2.1mm | |
| 1 | TE Connectivity | 6-104935-0 | Shrouded locking right angle header, 10 pos | |
| 3 | McMaster-Carr | 91116A240 | SS washer, M2, 7mm OD | |
| 3 | McMaster-Carr | 91292A833 | SS socket head screw, M2x0.4mm, 10mm lg | 1.5mm hex |
| 2 | McMaster-Carr | 92095A104 | SS button socket head screw, M2x0.4mm, 10mm lg | 1.3mm hex |
| 2 | McMaster-Carr | 92196A082 | SS socket head screw, 2-56, 7/16" lg | 5/64" hex |
| 2 | McMaster-Carr | 92320A453 | SS unthreaded spacer, #2, 1/8" OD, 5/32" lg | |
| 1 | Omron | D2F-01L3-D | SPDT microswitch | |
| 1 | Soyo | SY20STH42-0804A | NEMA 8 stepper motor, 5.4Ω, 4.2 oz in | |
| 1 | TE Connectivity | T4070014041-001 | M8 connector, 4 pos rear mount, wire leads | |

*Optical system*

| | | | | |
|---|---|---|---|---|
| 1 | Edmund Optics | 49-030 | Longpass filter 700 nm 12.5mm | |
| 1 | Edmund Optics | 67706 | 4x finite conjugate RMS | |
| 1 | Basler AG | acA5472-5gm | 20MP 1" monochrome GigE camera | |
| 4 | McMaster-Carr | 92196A106 | SS socket head screw, 4-40, 1/4" lg | 3/32" |
| 1 | McMaster-Carr | 99091A190 | Black nylon hex head screw, 8-32, 1/4" lg | 1/4" ext hex |
| 1 | Thorlabs | CCM5-G01 | ThorLabs CCM5-G01 protected turning mirror | |
| 1 | Thorlabs | SM05A1 | C-mount camera to SM05 male threads adapter | |
| 1 | Thorlabs | SM05L05 | SM05L05 lens tube, 0.5" thread dp | |
| 1 | Thorlabs | SM05M20 | SM05 Lens Tube Without External Threads, 2" Long | |
| 2 | Thorlabs | SM05RC | Slip mount for SM05 lens tube | |

| | | | |
|---|---|---|---|
| 1 | Thorlabs | SM05V05 | Ø1/2" Adjustable Lens Tube, 0.31" Travel |
| 1 | Thorlabs | SM1A1 | SM1 to SM05 adapter |
| 1 | Thorlabs | SM1A3 | Internal RMS to external SM1 adapter |
| 1 | Thorlabs | SM1L05 | SM1 lens tube, 0.5" lg |

***Illuminator and focus lock***

| | | | | |
|---|---|---|---|---|
| 8 | Marktech | MTE1074N1-R | NIR T1-3/4 LED, 4mW, 740nm peak | |
| 1 | LEDynamics | 4006-030 | Inline current regulator module, 30mA | |
| 1 | McMaster-Carr | 1173N031 | Silicone O-ring, 1/16" wd, 1.88" OD | |
| 2 | McMaster-Carr | 90669A074 | SS Brass-Tip Set Screw, 2-56, 1/8" lg | 0.035" hex |
| 3 | McMaster-Carr | 92785A053 | SS cone point set screw, 2-56, 3/16" lg | 0.035" hex |
| 5 | McMaster-Carr | 92395A311 | Flanged split brass threaded insert, 2-56, 0.136" lg | |

***Specialty material stock or parts to be modified***

| | | | |
|---|---|---|---|
| 1 | McMaster-Carr | 1375K29 | 10 tooth MXL pulley, for motor pulley 3-1263 |
| 1ft | McMaster-Carr | 6720T12 | Bend-and-stretch-rated cable stock for illuminator |
| 1 | FTDI Chip | TTL-232RG-VSW5V-WE | USB to TTL serial cable, 5V, bare wire ends |
| 1 | Tensility Intl | 10-05920 | M8 cable assy, 4 pos A key socket, 3 m, bare wire |

***Support equipment***

| | | | |
|---|---|---|---|
| 1 | Globtek | TR9KI2700LCPCIMR6B | 24V 65W power supply |
| 1 | TP-Link | TL-PoE160S | 30W PoE+ injector |
| | * | * | Computer with gigabit ethernet port |
| | * | * | Add'l cabling as necessary for pwr, ethernet, etc |

***Utility / misc.***

| | | | |
|---|---|---|---|
| 1 | ThorLabs | SM05L30 | Stackable 1/2" lens tube, 3" internal thread length. Used as an alignment tool. |
| 1 | McMaster-Carr | 3009A118 | (or similar) 1/8" drill rod for use as press tool |
| 1 | Henkel | 680522 | Loctite 454 (or similar) thixotropic CA adhesive |
| 1 | Henkel | 2765219 | Loctite 7452 (or similar) cyanoacrylate accelerator |
| | * | * | General essentials and consumables not listed: Hookup wire, heat shrink tubing, etc. |

### CCI instrument machined / printed parts

This section briefly describes the individual machined and/or printed parts used in the construction of the CCI instrument. The CCI flow cell components are covered separately in the CCI flow cell fabrication section.

Refer to the corresponding Onshape CAD models for part geometry for manufacturing. In cases where the solid model itself does not fully describe the part requirements a 2D drawing or other supplemental files(s) will be included in the Onshape document. When uploading models to a prototype manufacturing service it is important to make sure relevant details such as threaded holes are communicated correctly, and the process differs between vendors.

Refer to the Hardware build guide section for more context regarding the function of each part and order of assembly.

#### Chassis parts (3-1239, 3-1240, 3-1241, 3-1242)

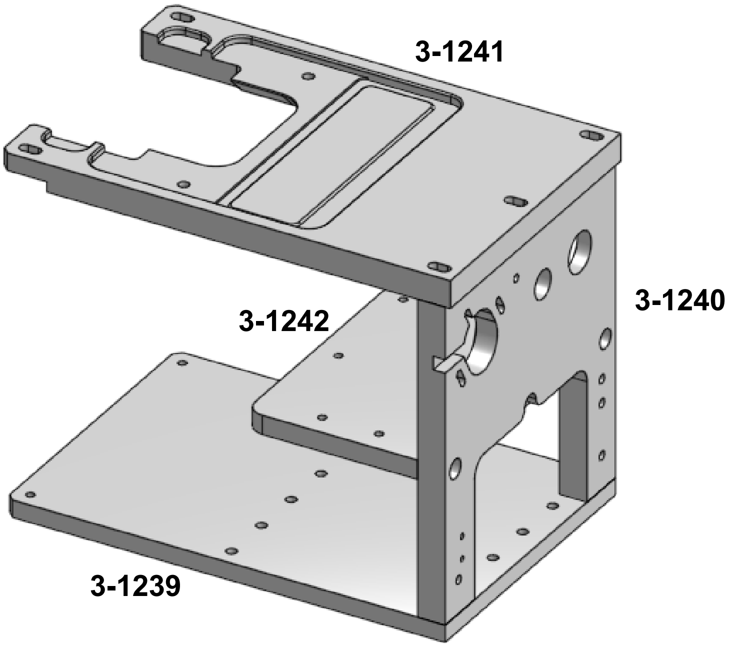

The structural frame includes four CNC machined aluminum plates:

| # | Description | Material |
|---|---|---|
| 3-1239 | Bottom plate | Aluminum 7075-T651 |

| | | |
|---|---|---|
| 3-1240 | Vertical plate | Aluminum 6061-T651 |
| 3-1241 | Top plate | Aluminum 6061-T651 |
| 3-1242 | Shelf | Aluminum 6061-T651 |

The 7075 alloy was selected for the bottom plate on our prototypes for increased stiffness, however the corrosion resistance of 6061/6063 may be preferable due to likely exposure to spilled media, etc.

The shelf (3-1242) primarily serves as a platform for the motor control board and attachment point for the illuminator power cable and is not a critical structural component, so a printed part could be substituted to save cost.

#### Optics platform parts (3-1243, 3-1246, 3-1247, 3-1250)

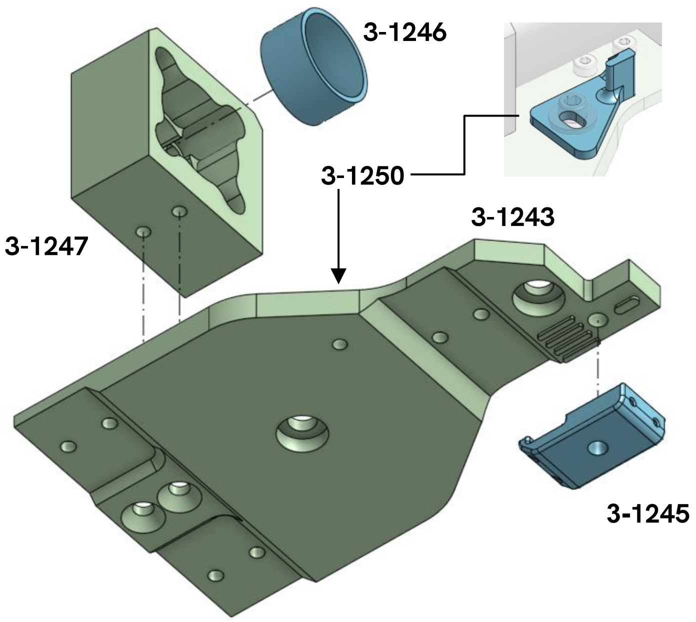

The optics platform / carriage structure consists principally of two machined aluminum parts:

| # | Description | Material |
|---|---|---|
| 3-1243 | Optics platform base | Aluminum 7075-T651 |
| 3-1247 | Mirror cube bracket | Aluminum 6061-T651 |

7075 alloy was selected for the base plate for its mechanical properties but 6061 could be substituted.

The spacer ring (3-1246) is made by cutting a 0.31" length from 5/8" OD x 0.058" wall 6061-T6 aluminum tubing (e.g. McMaster-Carr 9390N15).

The belt clamp body (3-1245) is made from SLS-printed glass-reinforced Nylon and reinforced with stainless pins (see Hardware build guide). This approach was motivated by a significant cost savings compared to ordering a CNC machined version of this small part in prototype quantities. Barring cost concerns, a solid machined part would be a simpler solution.

The homing switch flag (3-1250) was made by FDM printing in PLA plastic but the choice of material is immaterial for this part.

#### Motor pulley (3-1263)

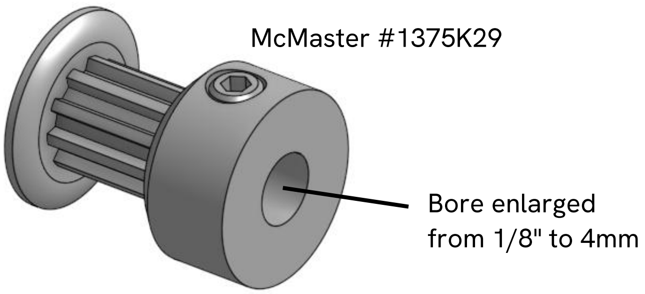

The motor pulley is made from a commercial 10-tooth, 1/8" bore aluminum MXL pulley (McMaster-Carr 1375K29), modified to fit on the 4mm diameter shaft of the stepper motor by re-drilling and reaming the bore on a lathe.

This process has proved to be troublesome and has a poor yield. The pulley is a press-fit assembly of three pieces. Enlarging the bore must be done carefully as it leaves very little material thickness in the middle piece where it mates with the hub and the end flange. In addition, any eccentricity becomes problematic for proper operation of the stage drive mechanism.

#### Pipette tip guide / flow cell clamp (3-0885)

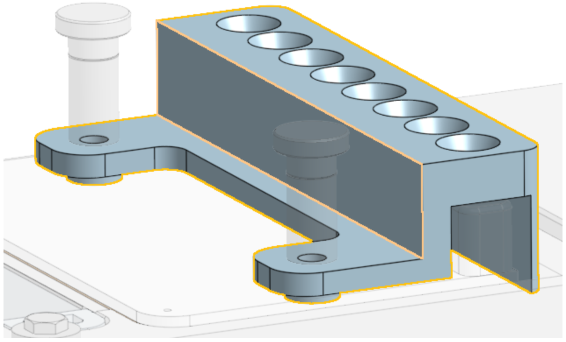

This piece is installed on top of the flow cell and held down with the thumbscrews. It was originally intended to guide the pipette tips into the relatively small holes on the pipette tip interface. The current design of the pipette tip interface itself incorporates lead-in cones to guide the tips into the openings, so now the primary purpose of the tip guide is to provide a bearing surface for the thumbscrews and provide backup protection against the pipette tip interface being stabbed by grossly misaligned pipette tips.

The pipette tip guide was made using SLA printing in Formlabs Black V4 resin. The choice of material is not critical but a dark non-reflective surface is preferable.

#### Darkfield illuminator parts (3-1253, 3-1254, 3-1261)

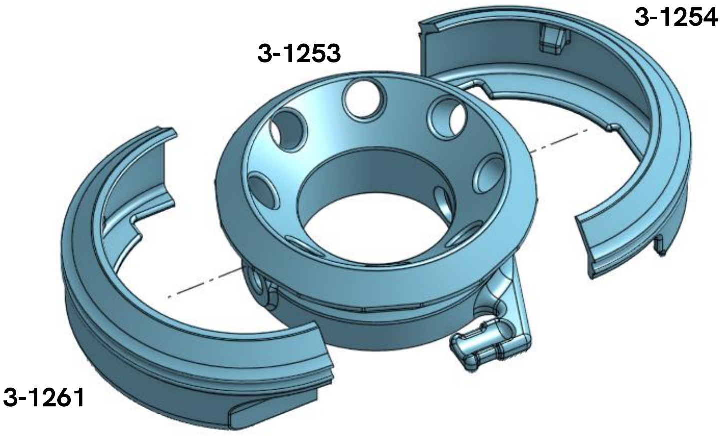

The illuminator main body (3-1253) forms the primary mechanical structure of the illuminator; the illuminator shroud front half (3-1254) and rear half (3-1261) conceal the LED wiring and provide a small amount of shielding from incidental liquid exposure.

All three parts are made using SLA printing in Formlabs Tough 2000 resin.

#### Focus locking sleeve and focus knob (3-1252, 3-1260)

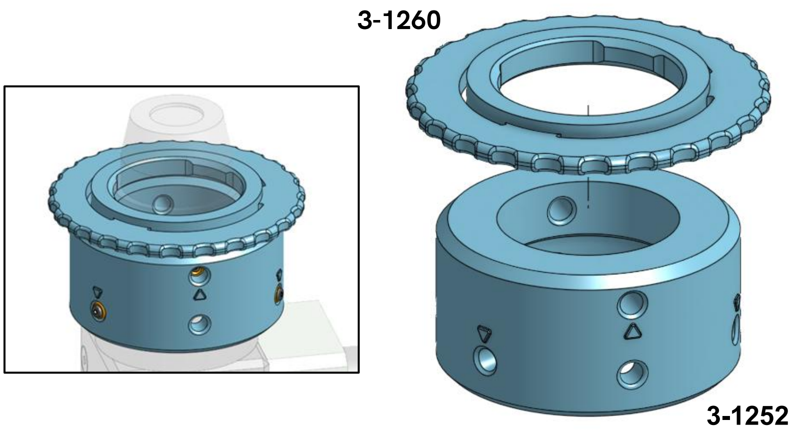

The focus locking sleeve body (3-1252) is part of a simple focus adjustment system wherein the height of the objective can be adjusted by screwing an adaptor  in and out of a threaded tube and locked by jamming a set screw against the side of the objective. The part is finished by installing threaded inserts which hold 3 cone-tipped (for securing the sleeve in place on a tube) and one brass-tipped (for jamming against the objective) set screws. An access hole opposite each insert location acts as a drill guide for cleaning up the printed bore, then a press tool (a short length of 1/8" drill rod) is inserted through the access hole to press the insert into place.

The focus knob (3-1260) simply press-fits onto the outside of the objective and provides a way to rotate it with one hand while the other hand keeps the illuminator from spinning (there is no clearance for fingers to reach the objective itself in this situation).

As with the darkfield illuminator parts, both of these parts are made using SLA printing in Formlabs Tough 2000 resin. Using a material with different properties may require slight design adjustments. As discussed in the Hardware build notes section, the focus locking sleeve may require some additional post-machining.

#### Waste trough (3-1273)

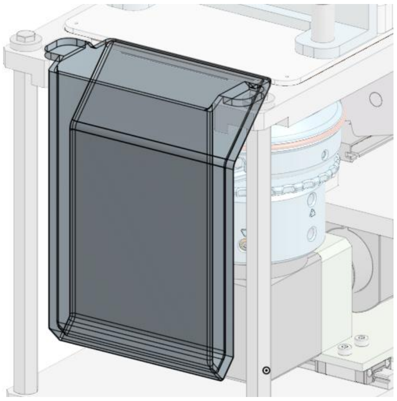

The waste trough hangs from the top plate of the CCI frame underneath the flow cell and collects the liquid that flows out of the drain ports.

The waste trough is made by SLA printing in Formlabs Clear Resin V4. It is important to use a non-opaque material so that the liquid level is readily visible, reducing the risk of overflow due to failure to empty waste from a previous run.

The waste trough could be considered a semi-disposable item, although in practice the material appears to hold up well to long term use and repeated disinfection with bleach.

A modified design with a drain port plumbed to an external waste trap could hypothetically remove the need to manually empty the trough between runs.

### CCI instrument hardware build guide

The order of operations presented here is somewhat arbitrary.

#### Flow cell fasteners

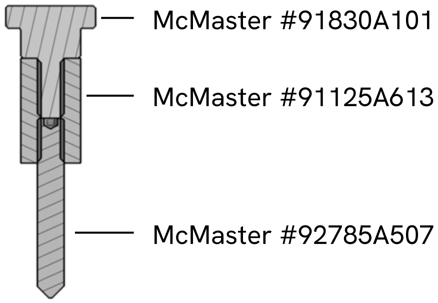

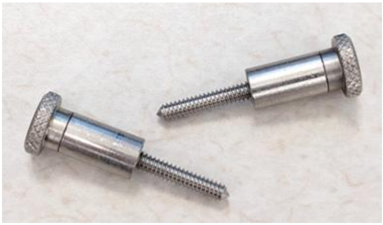

Assemble two flow cell fasteners by firmly threading together the three component parts and securing with permanent threadlocker or epoxy.

#### Belt clamp

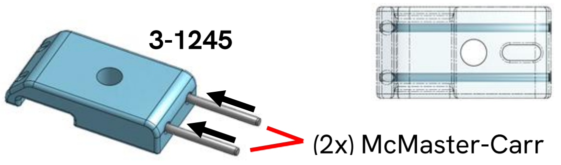

The belt clamp is completed by simply inserting two 1mm x 20mm stainless steel pins (McMaster-Carr 91585A093) into the 3D-printed belt clamp body (3-1245) using a small arbor press. If necessary, a bench vise or other tool could potentially be used instead, or the holes could be drilled for a clearance fit and the pins glued in place.

#### Focus locking sleeve

Completing the focus locking sleeve (3-1252) involves installing a total of four threaded inserts. Essentially the installation process is the same as for the illuminator collar but without the need to brace the part to keep it upright.

After setting the four inserts, install three cone point set screws (McMaster-Carr 92785A053) in the "lower" holes (actually on top in the pictures below) and a brass-tip setscrew (90669A074) in the "upper" hole.

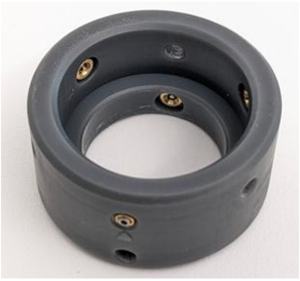

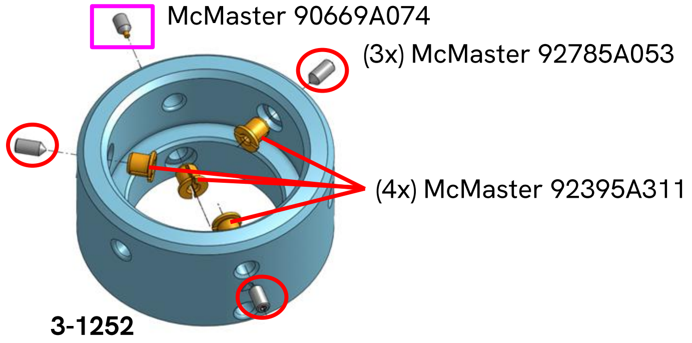

### Motor controller board

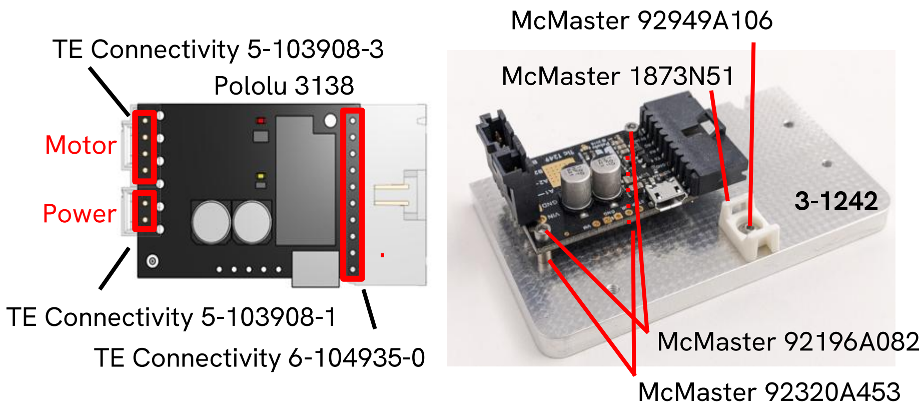

Solder three connectors onto the Tic board as indicated, then mount the board on the shelf plate 3-1242  using the specified screws and spacers.

Also mount a cable tie anchor at the indicated location.

### USB serial communications cable

This cable connects the host PC to the serial interface for the motor controller on the CCI instrument. It is made by splicing an M8 socket pigtail cable (Tensility 10-05920) to an adapter cable with a TTL-level USB serial port interface in it (FTDI TTL-232RG-VSW5V-WE).

This combination gives a maximum reach of up to 15 ft if using the full length of both half cables. Cut down the length of the M8 pigtail before assembly to produce a length that is not excessive for your installation.

Only three connections are used -- the two serial data lines and signal ground. Unused wire ends on the FTDI cable side should be dealt with appropriately to ensure that they don't contact each other.

| Wire color,<br>FTDI cable end | Function | Wire color,<br>M8 cable end |
|---|---|---|
| ~~Red~~ | ~~+5V~~ | ->(don't connect) |
| **Black** | GND | **Black** |
| **Yellow** | RxD (CCI->PC) | **White** |
| **Orange** | TxD (PC->CCI) | **Brown** |

| | | |
|---|---|---|
| ~~Green~~ | ~~RTS~~ | ->(don't connect) |
| ~~Brown~~ | ~~CTS~~ | ->(don't connect) |
| (don't connect)<- | n/a | ~~Blue~~ |

#### Wiring harness prep

The **I/O harness** connects the homing switch and serial port to the 10-pin header on the Tic board. Start by wiring the N.O. side of the homing switch to positions 6~7 of the 10-position connector with at least about 7cm of movable wire length, as illustrated below.

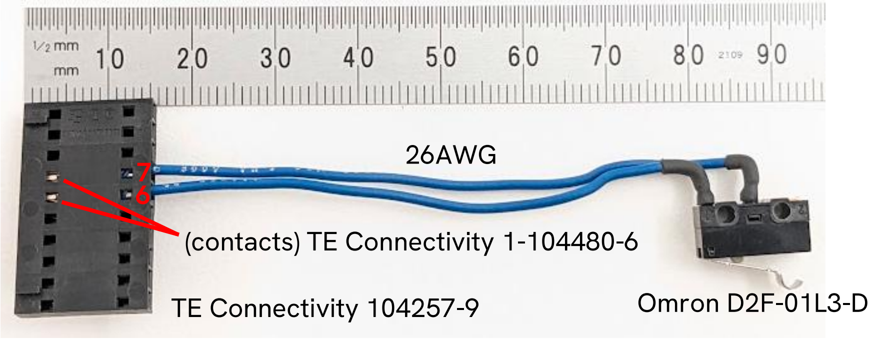

Next add the serial communications lines from the M8 bulkhead connector (TE Connectivity T4070014041-001): cut off the blue wire from the M8 connector close to the base, trim the remaining leads to about 12cm long and terminate them in the 10-position connector per the table below to complete the I/O harness.

| M8 plug position no. | Wire color | Function | Header position no. |
|---|---|---|---|
| 4 | **Black** | Signal ground | **1** |
| 1 | **Brown** | Serial data PC→Tic | **4** |
| 2 | **White** | Serial data Tic→PC | **5** |
| n/a | | Switch COM* | 6 |
| n/a | | Switch NO* | 7 |
| | | | * interchangeable |

Next wire the **power harness** with the barrel jack and two two-pin connectors in parallel as shown below.

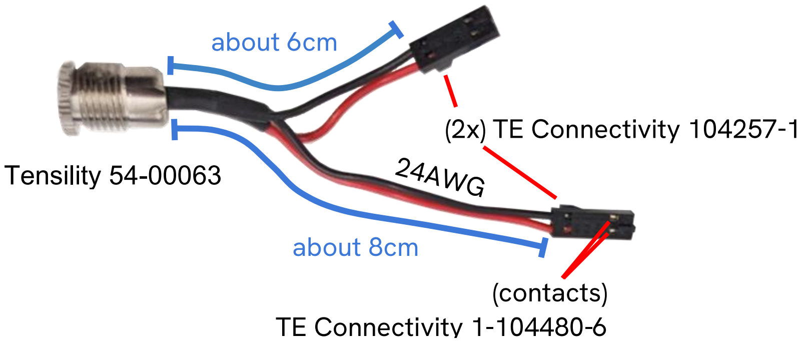

| Barrel jack terminal | Function | Header position no. |
|---|---|---|
| Outer | Ground | 1 |
| Inner | +24V | 2 |

Note that although the barrel jack is a front-mount design, the 2-pin connectors will fit through the mounting hole for the jack, so the harness does not have to be assembled in-place.

Finally, terminate the **motor cable** included with the stepper motor with a 4-pin connector as shown:

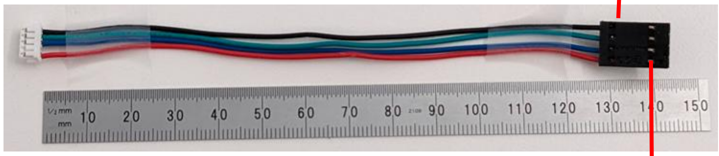

| Wire color | black | green | blue | red |
|---|---|---|---|---|
| Header pos. no. | 1 | 2 | 3 | 4 |

Note that the motor leads are 28AWG which requires different contacts than the previous 3 connectors.

#### Darkfield illuminator collar

**Note**: Before proceeding to build the illuminator, check the fit of the LED holes by inserting an LED into each one and verifying that the LED passes freely into the hole and that the flange of the LED bottoms out squarely on the flat land surrounding the hole. This should be done first in case any rework of the part is needed.

##### Threaded insert

The illuminator collar slips over the end of the objective and is secured in place by bearing down on it with a soft-tipped setscrew. The setscrew is installed in a flanged threaded insert which is installed from the inside of the bore. The access hole opposite the threaded insert provides access for a press tool but the protruding rim of the part presents a challenge when attempting to apply force in a controlled manner. The design includes a small flat boss around the through hole which can be used to brace the part against a suitable object such as a square wooden bar as illustrated below.

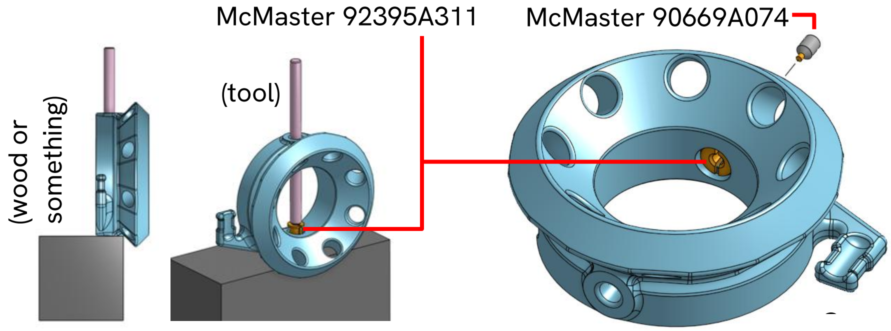

Before attempting to install the insert, run a 1/8" drill through the two opposing holes in the base of the part as a "clean-up pass" in case they are undersized or distorted.

With the part properly supported, the insert should be able to be pressed in easily using a piece of 1/8" drill rod held in a drill press or whatever method is readily available. After installing the insert, drive in a sacrificial #2-56 cap screw (from the outside hole) to expand it. Make sure a cap screw can be threaded all the way through the insert without binding up before attempting to drive in a setscrew with a hair-thin hex key. The inserts are easily damaged by mis-installation, so it is suggested to order extra stock to avoid having to try to work with a compromised insert.

After a successful function test, install the brass-tipped setscrew (90669a074) in the insert from the outside, so that the tip faces the center bore.

#### LED installation and wiring

Electrically, the illuminator consists of a loop of 8 LEDs broken in the middle by an inline current regulator and supplied with 24V DC, as illustrated below.

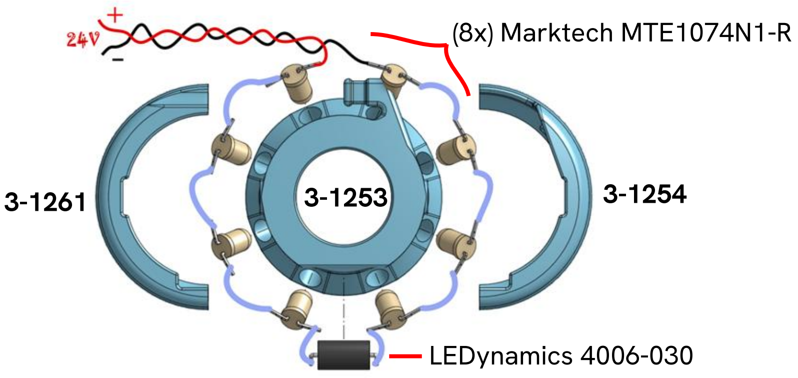

Physically, the LEDs are inserted into angled holes and held in place with glue, then the leads are bent and soldered together. Power is supplied from the instrument's power input harness via a flexible cable that moves with the stage.

We have found that the installation process is easiest when the LEDs have a free fit or a very light interference fit in the holes. If the fit is slightly too tight, the part may be adjusted by running a #8 or #7 drill through all the holes. We use a gel-type cyanoacrylate glue (Loctite 454 or similar), so a slightly loose fit is tolerable and less risky than a too-tight fit.

For each LED, apply a small blob of adhesive on top of the rim, insert the LED into the hole, and rotate it so that the flat side of the rim faces the counterclockwise direction (when viewed from above). After all 8 LEDs are placed, use tweezers or another pointed tool to poke the base of each LED and make sure it is fully seated. Once satisfactory placement of all LEDs is confirmed, apply a spray of accelerator (e.g. Loctite 7452). Additional glue can then be added on for security if needed.

Bend the leads of the current regulator at a 90 degree angle close to the body and then glue it into the curved recess in the orientation shown below, making sure the end marked "+" is facing the *clockwise* direction (opposite to the flats on the LEDs).

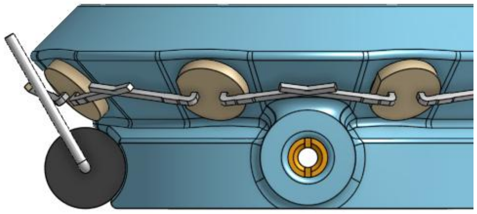

Manipulate the leads of the current regulator and its neighbor LEDs into contact and trim away as much excess material as possible. Repeat this down the chain but stop at the last pair of neighbor leads, as this is where the power cable will be connected. Test-fit the shroud halves (3-1261, 3-1254) periodically in the process in order to solve clearance issues as they arise.

Make the power cable for the illuminator by cutting a 180mm length of flex cable (McMaster 6720T12), removing about 20mm of outer jacket at each end, and terminating one end with a 2-position header (TE Connectivity 5-103908-1), following the polarity convention of the power harness made earlier (i.e. positive on pin 2). Finally, solder all of the connections around the loop and connect the power cable (red wire to the curved side of the first LED, green wire to the flat side of the last LED). Use a cable tie to secure the outer jacket of the cable to the arm for strain relief as shown.

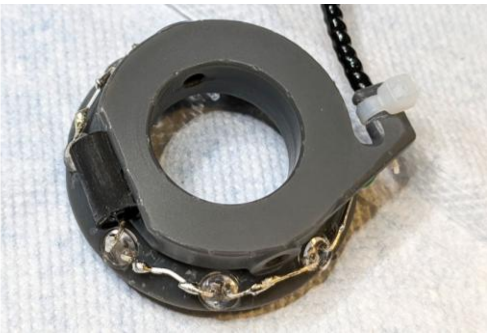

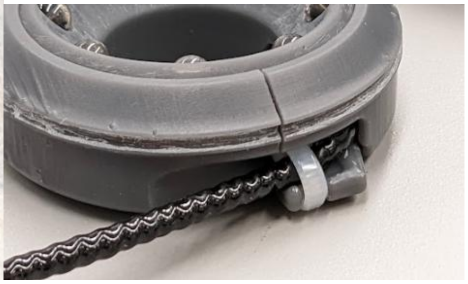

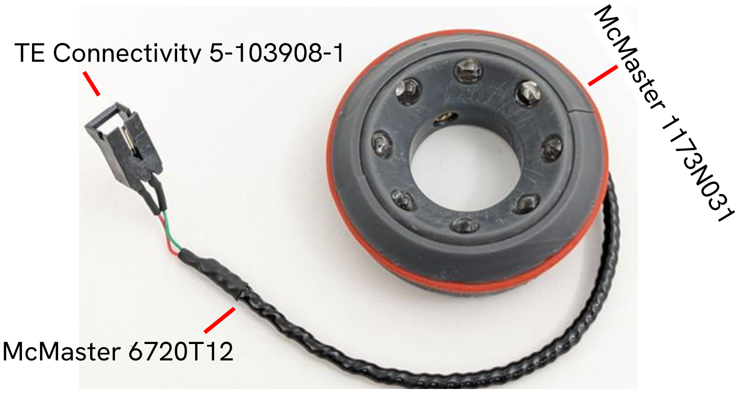

After fitting the shroud halves, the silicone O-ring (McMaster 1173N031) can be hooped around the shell halves to hold them together. Test with power to verify that the LEDs light up (the output will be visible to human vision as a dim red glow), then set the completed illuminator aside.

#### "Middle section" and motor test

First install three idler assemblies on the vertical plate 3-1240 as illustrated. Note that there are four holes where the idlers can be mounted; the correct three locations are indicated in the diagram below.

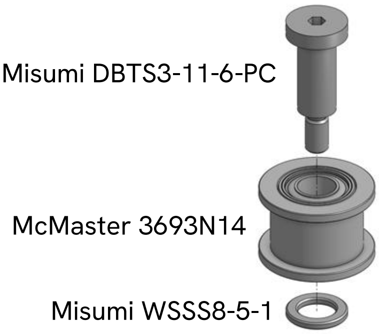

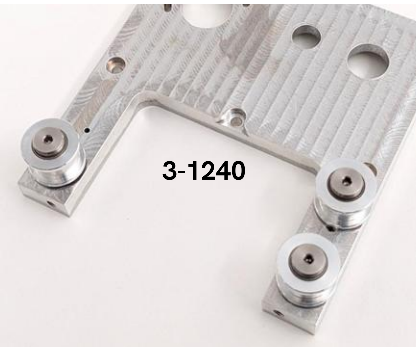

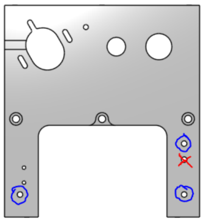

Next install the serial connector, the power connector and the homing switch on the vertical plate as shown:

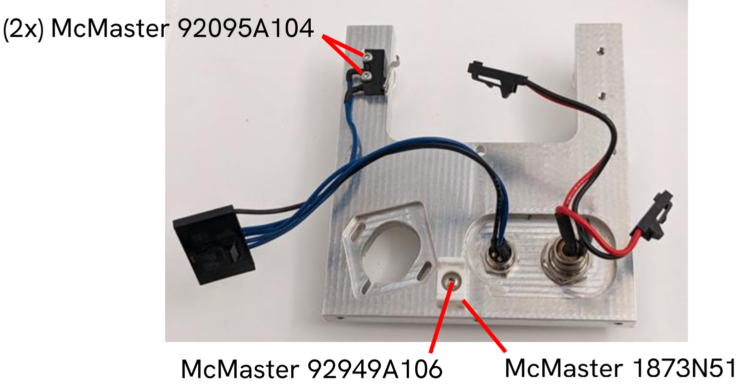

Attach the shelf plate to the vertical plate with 3 screws as indicated, taking care to route the switch wires through the notch in the shelf plate. The I/O and and motor power connectors can then be plugged in.

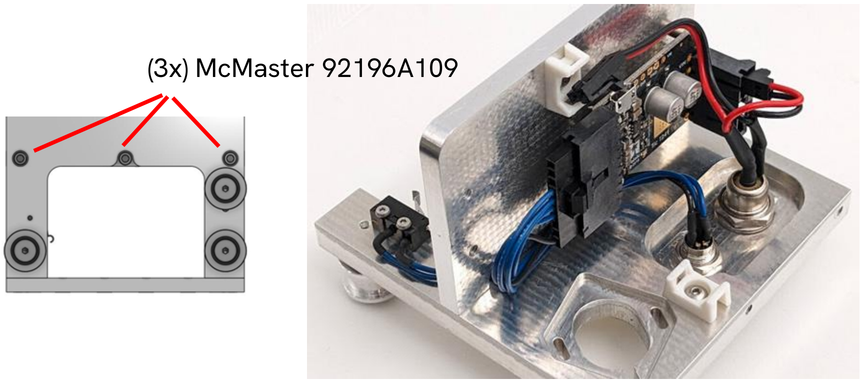

Attach the motor pulley to the motor shaft loosely (only screw in the set screw enough to key the pulley to the flat on the shaft, then back off). Install the motor on the vertical plate as shown, with the screws installed loosely so that it can move along the slots. Plug in the motor cable and secure it to the upper cable tie anchor.

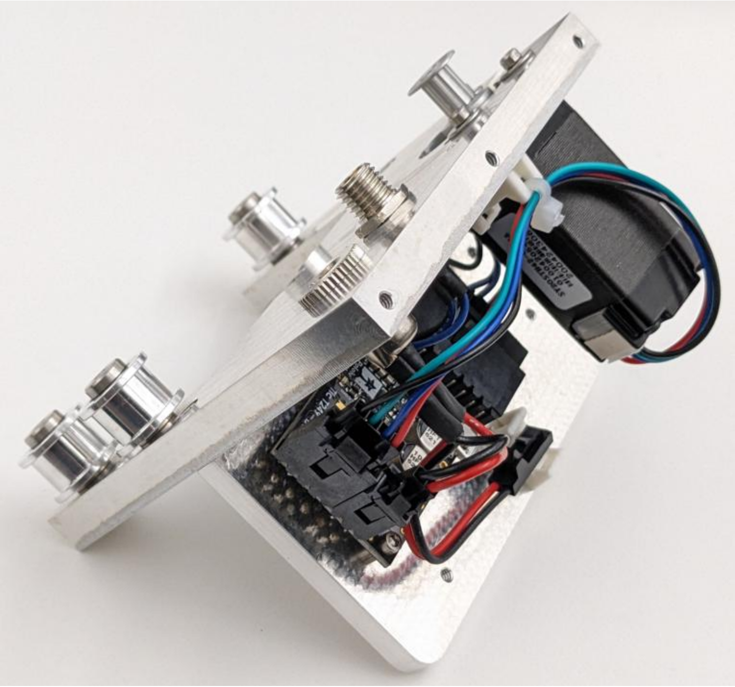

It is highly recommended to do initial setup and testing of the Tic board at this point. Find and install the appropriate software download for your OS from the manufacturer's website: https://www.pololu.com/docs/0J71/all

Connect the Tic board to a PC via its MicroUSB connector and plug a 24V power supply (e.g. Globtek TR9KI2700LCPCIMR6B) into the power connector on the bulkhead. Run the Tic Control Center software. It should immediately connect to the board and start displaying status information:

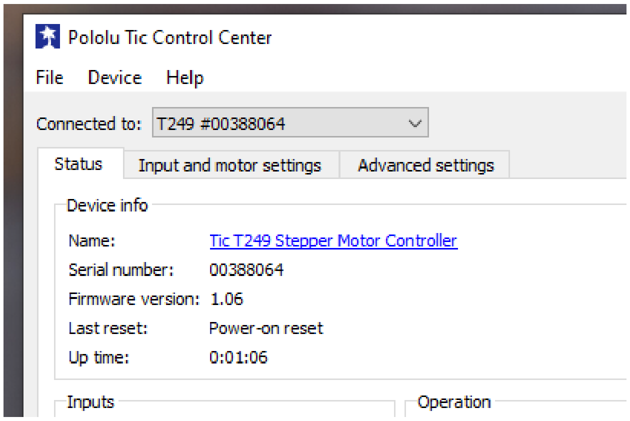

To initialize the Tic board with the correct configuration for the CCI, select "File" -> "Open settings file...", open the tic_settings_cci2.txt file obtained from the CCI GitHub repository, then use the "Apply Settings" button to send the new settings to the board.

Press the limit switch manually and you should see an indicator appear:

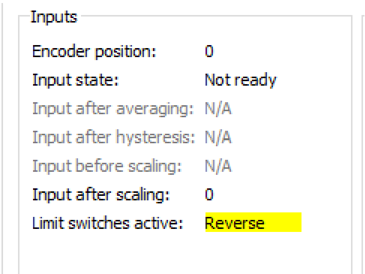

You can simulate homing the stage using "File" -> "Go home reverse". The motor should start slowly turning. Push the homing switch and it should reverse direction, then stop when you let go of the switch. Finally, you should be able to use the "Set position" and "Set velocity" controls at the bottom of the window to manually make the motor move.

#### Linear guide mounting

**Warning:** Without endstops in place it is easy to inadvertently let the linear guide carriages run themselves off the rails when handing the bottom plate assembly, especially once the weight of the optics platform is attached. The carriages coming off the rails is not necessarily a catastrophic event, but the optical system meeting the floor could be.  Until the "Middle Section" is attached to the bottom plate, consider using tape to avoid unexpected movement.

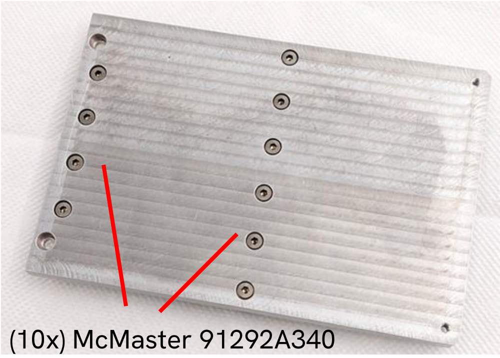

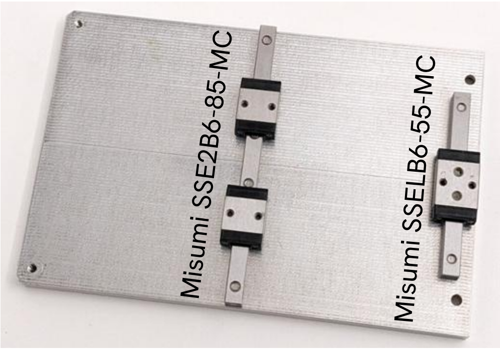

Line each linear guide rail up with its corresponding holes in the bottom plate and install all of the screws as indicated but only take up the slack, do not tighten. Looking at the bottom of the bottom plate, manually manipulate the longer rail so that all of the screw heads for that rail appear centered in the counterbores, then gradually tighten them in multiple passes from the center outward.

The shorter rail should be left slightly loose for now; it will be aligned and fastened in a later step.

#### Optics platform

Install the Heli-Coil insert (McMaster-Carr 96246A094) at the indicated location on the optics platform base plate (3-1243) if this operation was not ordered as part of the manufacturing of the part.

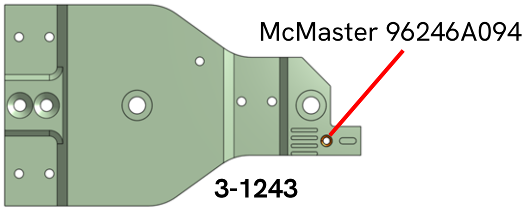

Attach the homing flag and tube clamps as shown below, but *do not fully tighten the tube clamps down to the plate yet* -- they should be able to rotate slightly but should not wiggle freely.

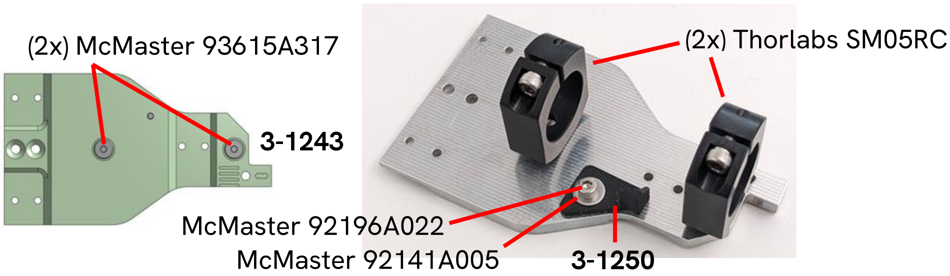

Slip a Thorlabs SM05L30 tube through the clamps to align them and tighten the clamps until they both grip the tube, then fully tighten the mounting screws for the clamps. Open the clamps and remove the tube.

#### Turret and camera tube assembly

Install the RMS-to-SM1 adapter (Thorlabs SM1A3) on the objective (Edmund Optics 67706). Tighten firmly and consider mounting the adapter permanently using a threadlocker.

Place the 700nm longpass filter (Edmund Optics 49-030) in the Thorlabs SM05L05 lens tube and secure it using the locking ring included with the tube.

Install the black nylon screw (McMaster 99091A190) in the open mounting hole on the housing of the turning mirror (Thorlabs CCM5-G01).

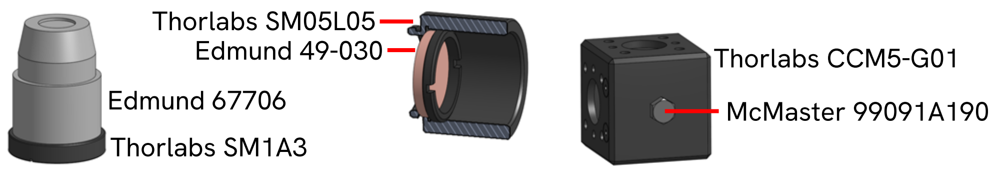

Facing the side of the mirror cube you just installed the screw in, with the openings facing up and left, screw the Thorlabs SM05-to-SM1 adapter (SM1A1) into the top hole followed by the 0.5" SM1 tube (SM1L05). Make these connections as tight as reasonably possible.

Install the mirror cube bracket (3-1247) on the side of the mirror cube with 4 screws as shown; *leave the screws loosened by 1/4 turn for now*. *Make sure the two screw holes on the outside of the bracket are facing down* (opposite from the direction the objective is pointing).

Screw the SM05L05 with the filter in it onto the end of the SM05 "adjustable" lens tube (SM05V05). Slip the spacer ring (3-1246) over the external threads and then screw the tube into the mirror block. The tube should bottom out on the spacer ring, such that the mirror cube bracket still has a small amount of translational freedom.

Thread the objective in its adapter partway down into the top tube. Do not bottom it out. The SM1 adapter and the tube are used as a crude focus adjustment mechanism.

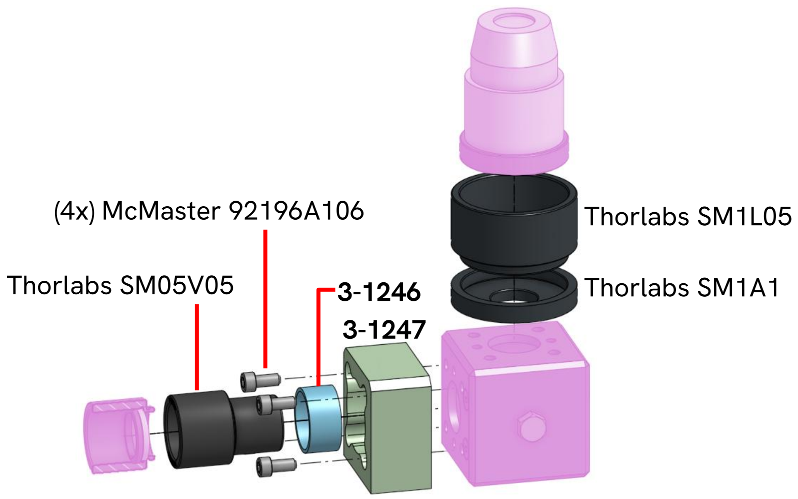

Install the C-mount adapter (Thorlabs SM05A1) tightly into the 2" SM05 tube (SM05M20). Install the adapter onto the camera (Basler acA5472-5gm).

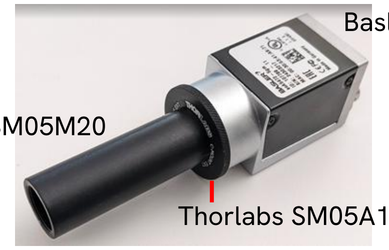

#### Optical system

Temporarily install the long SM05 tube (Thorlabs SM05L30) onto the open leg of the turret assembly to use for alignment. Insert the tube through the clamps. Tighten the clamps just until the tube has absolutely no wiggle but can still move axially without needing excessive force. Line up the mirror cube bracket mounting holes with the corresponding countersunk holes on the base plate. Install and tighten two screws to attach the bracket to the base plate as indicated below.

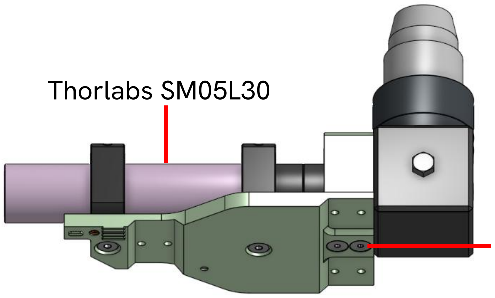

Viewing the assembly from the side opposite the long tube as shown below, make sure the turret is pointing straight up (perpendicular to the base plate) and adjust the angle if necessary. Ensure the opposite side of the cube is pushed all the way up against the cube bracket.

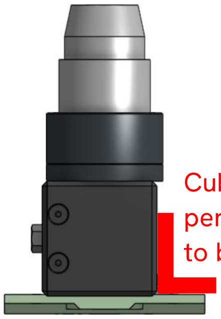

Tighten the 4 screws inside the mirror cube bracket using the short leg of a 3/32" hex key. Loosen both tube clamps and unscrew and remove the long tube.

Position the carriages of the linear guides to line up with the mounting holes on the optics platform baseplate and place the optics platform onto the linear guides. Install all 6 screws as indicated below, without tightening.

(6x) McMaster 91292A004

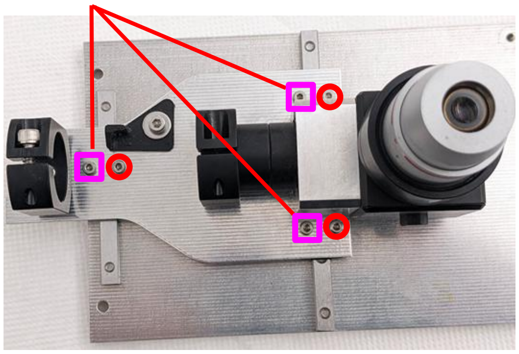

Turn the three screws designated by magenta squares until lightly snug. Move the stage back and forth from one end of travel to another. If there is excessive drag, re-loosen the three screws and try re-snugging them with less force than before while gently wiggling the stage back and forth to feel for any change in friction. Once the stage moves smoothly in this configuration, snug the screws indicated by the red circles, then fully tighten the magenta square screws followed by the rest.

Move the stage to one extreme of travel and gently tighten the outermost screw on the short rail on that side. Repeat with the stage at the other extreme. If increased drag develops at this point, try releasing and re-fastening the carriage mounting screws for the short rail. Once smooth movement is achieved, tighten all screws on the short rail in a few passes moving outward from the center.

Install the camera by sliding the attached tube through the clamps until it stops. Orient the camera as shown (silver bottom facing the same direction as the nylon screw on the mirror box) and secure it by tightening the two clamps.

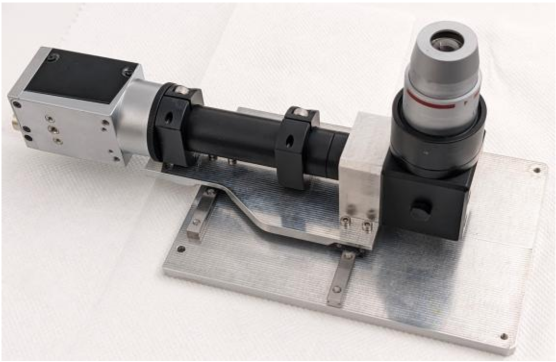

#### Drive train

Attach the "middle section" to the bottom plate with two screws, making sure the edges of the two parts are aligned, and then install the two long standoffs as indicated below.

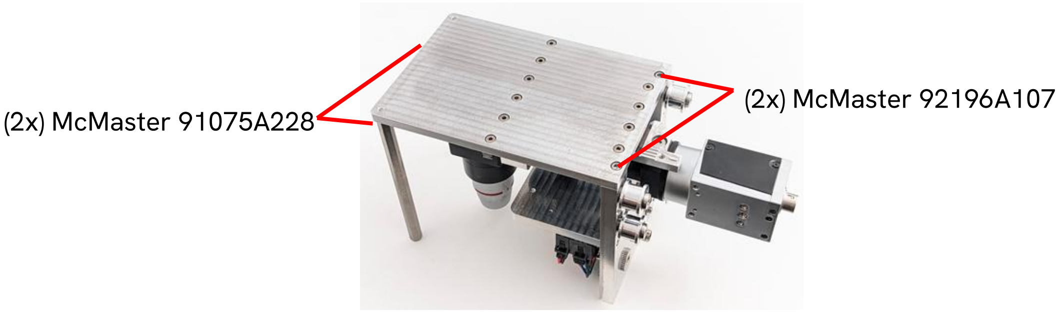

Attach the tensioner spring (Partsbuilt 3D R2C2-SPRING-BELT) to the drive belt (McMaster 1679K626) as shown. Slide the motor in its mounting slots towards the center of the vertical plate. Install the drive belt by hooking around the motor pulley and two of the idlers, then slipping it over the edge of the third roller.

Facing the outside face of the vertical plate, with the instrument upside-down, move the stage all the way to the right as shown below. Keeping the stage in this position, lift the belt off of the teeth on the optics platform base plate and shift it so that the spring is in the position shown. Reengage the belt on the teeth and install the belt clamp with the screw and washer shown.

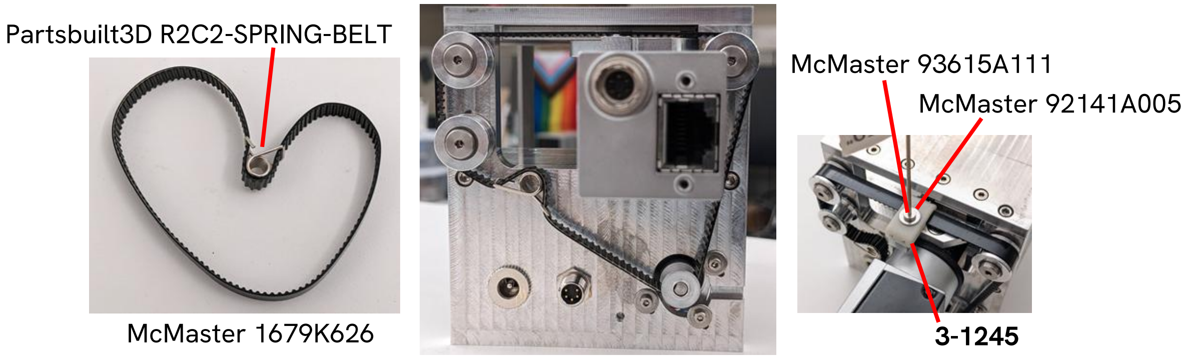

Push the motor fully back to the outermost position (you will encounter resistance from the spring) and tighten the mounting screws. Rotate the pulley so that the set screw is lined up with the slot in the plate as illustrated below. Adjust the axial position of the motor pulley to agree with the belt and the idler pulleys, then tighten the setscrew.

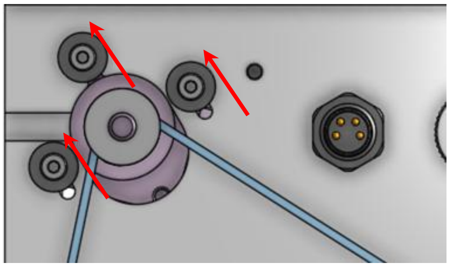 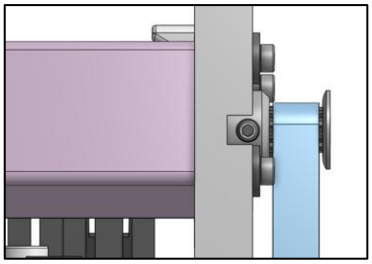

With the drive train now assembled, move the stage back and forth by hand from stop to stop to verify that it moves smoothly over the range of travel. Also verify that the homing switch activates slightly before the stage reaches the hard stop, and adjust the flag on the optics platform if necessary.

#### Install focuser and illuminator

Fit the focus locking sleeve assembled earlier onto the top tube on the turret and rotate it so that the upward facing arrow mark is on the same side as the printed markings on the mirror cube. Fasten it by *gently* tightening the three bottom set screws arranged around the bottom, just until the sleeve is immobilized.

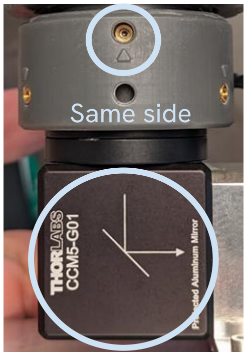

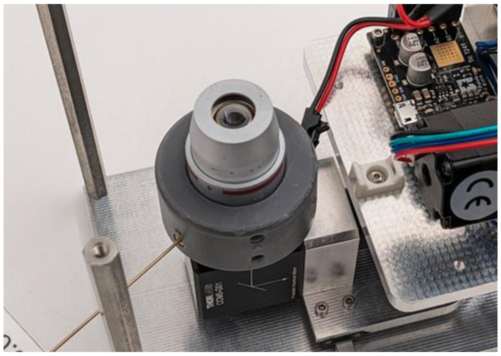

Check whether the objective can rotate freely. Overtightening the set screws that hold the collar onto the focusing tube will distort the tube and cause the threads to bind up. We have also found in multiple cases that the RMS-to-SM1 adapter ring on the objective seems to run slightly eccentric in the tube which causes the objective to interfere with the focus locking sleeve. If the latter is the case, it may be necessary to bore out the opening slightly to compensate for the eccentricity with extra clearance.

Install the focus knob by simply pressing it over the objective with the raised boss facing up as shown. The top of the boss should be slightly below the step on the objective.

Place the illuminator assembly built earlier onto the objective in the orientation shown below. It should rest on the step on the objective. Carefully secure the cable and header to the cable tie anchor with cable ties as shown. Plug the remaining connector from the power input harness into the header. Tighten the setscrew in the illuminator collar to secure it to the objective by inserting the hex key through the small hole in the shroud. If it is difficult to blindly align the hex key with the set screw this way, remove the shroud to access the set screw.

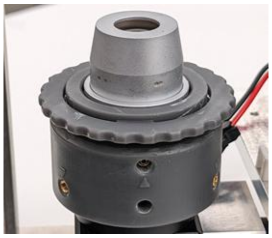

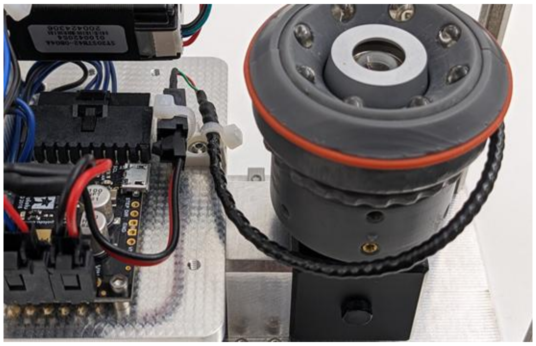

#### Top plate

Install the two Heli-Coil inserts (McMaster-Carr 96246A046) at the indicated locations from the top of the frame top plate (3-1241) if this operation was not ordered as part of the manufacturing of the part.

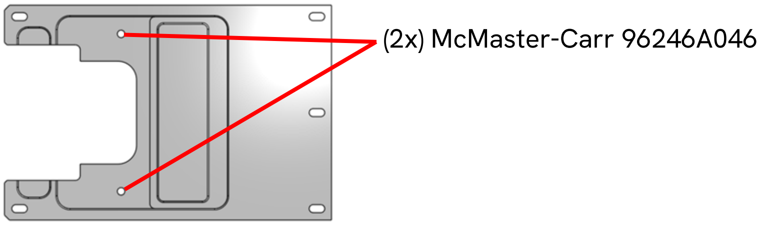

Place the top plate on the instrument and loosely install the 5 mounting screws and washers as indicated below. You will fasten the top plate after aligning it using the camera in a later step.

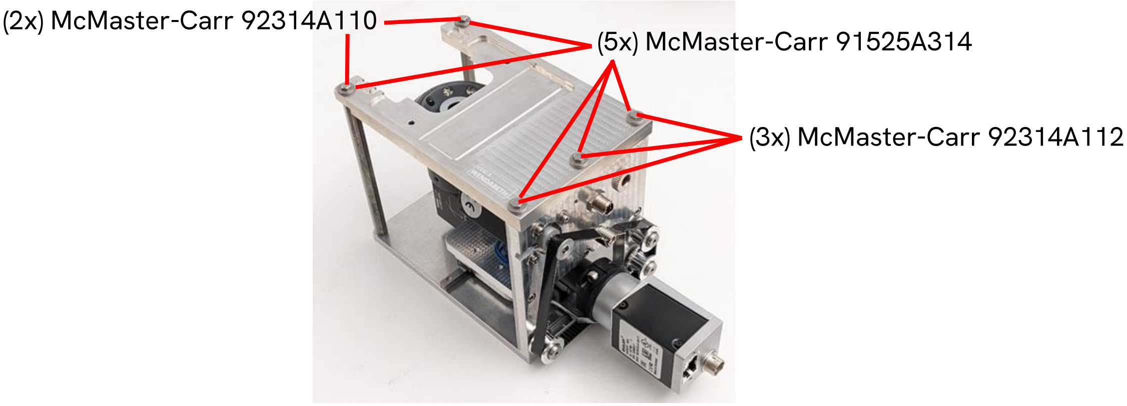

### Commissioning

#### Software setup

The Basler Pylon Camera Software Suite is not a strict prerequisite to run the Cell Counting Imager software, but it includes a camera viewer tool which is useful for testing and adjustment, so it would be useful to install this first.
https://www.baslerweb.com/en-us/downloads/software/

**Note!** This document covers installation and usage of the CCI software at a surface level in order to support initial commissioning of the CCI. The primary practical documentation for the CCI software is packaged with the source code to ensure it is up to date as the software evolves.

The CCI software and documentation can be obtained from the GitHub repository here:
https://github.com/czbiohub-sf/accs-cell-counting-imager-pub

The Cell Counting Imager software is a Python package that should be installed in a dedicated Python virtual environment. How you choose to do this may depend on your platform, preferences and existing practices. For example, we use Anaconda so the process to install the CCI software from scratch would look like this:

```
cd /dir/where/you/downloaded/it/cell-counting-imager-0.2.0/
conda create -n accs-cci python=3.12
conda activate accs-cci
pip install .
```

Refer to the CCI software documentation for further information.

#### Basic stage function test

Connect the 24V power supply to the CCI. The LEDs on the illuminator should appear to light with a dull red glow and there should be some indicator light activity from the TIC board. The stepper motor should immediately energize and start holding the stage in place, which you can confirm by gently pushing on the stage.

Connect the USB serial cable to the CCI and then to the PC. Once you have the CCI software installed, you can run the included cci_test command, cci_test --stage-auto

which will attempt to find which port the stage is plugged into and then run a movement test routine to make sure the stage functions correctly. If there is more than one FT232R based USB serial adapter connected then the program will list them for you and exit. If you are using a different type of cable or detection fails for some reason, you will need to check manually what port name your operating system assigned the virtual port to. You can specify the port to run the movement test with this variant of the command:

cci_test --stage COM4

You can also verify that the software can communicate with the camera with the following command, which simply captures an image and saves it in a JPEG file:

cci_test --camera

See the CCI software documentation for further details.

#### Using the Pylon camera viewer

As part of the CCI software setup process you will have installed the Basler Pylon Camera Software Suite, which includes the Pylon Viewer application. Review the online documentation to become familiar with how to operate the software:
https://docs.baslerweb.com/overview-of-the-pylon-viewer

Connect the CCI camera to the gigabit ethernet port on the PC via the PoE injector (TP-Link TL-PoE160S). Run Pylon Viewer, connect to the camera, and apply the following settings:

| | |
|---|---|
| Pixel format | Mono12 |
| Auto gain | Off |
| Raw gain | 0 |
| Gamma | Off |
| Binning factor, horizontal | 2 |
| Binning factor, vertical | 2 |
| Binning mode | Sum |
| Exposure time | 120 ms |

Enter Continuous Shot mode and you should see an image from the camera, which can be used for focusing and position adjustments as described in the following subsections.

#### Focus adjustment

If an actual flow cell and a sample of cells is not available, a convenient target for rough focus adjustment is an unassembled flow cell bottom plate (3-0305) or simply a piece of 1.5mm thick PMMA cut to the same shape without fiducials, with some fine scratches on the top surface.

To adjust the focus, loosen the setscrew in the illuminator collar until it can rotate freely, then loosen the upper setscrew in the focus locking collar and use the knob to rotate the objective. Note that this system is not very precise and the act of re-tightening the set screw tends to cause a significant shift in the focus. One can compensate somewhat by applying a counter-offset before tightening the screw and/or continuing to make adjustments while tightening the screw in small increments.

#### Top plate position adjustment

Mount a flow cell to the top plate, position the stage so that the fiducial mark in the first lane is vertically centered in the image, and adjust the alignment of the top plate so that the mark is not vignetted. Check against the fiducial in the last lane as well; if the fiducial is not in the same place in the X-axis then the top plate is not aligned squarely with the axis of travel of the stage.

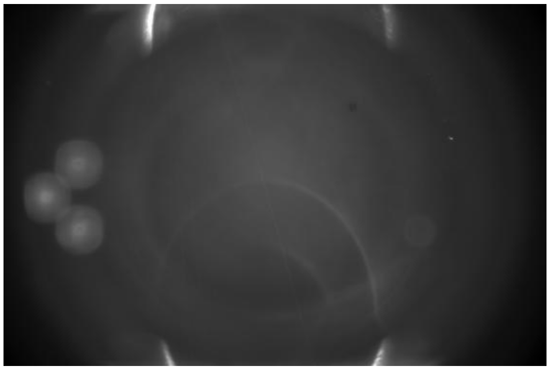

Once the result seems satisfactory, tighten the mounting screws to secure the top plate. Continue to observe the camera view while doing so as tightening the screws will likely affect the alignment.

#### Software stage alignment

The CCI software uses a configuration file to remember what positions to move to on a given instrument. Normally this only needs to be set up once for a given instrument unless something is changed mechanically or there is significant manufacturing variation in the lane positions between batches of flow cells or between individual flow cells.

Install a flow cell on the CCI, plug the USB serial cable into the instrument and the PC, connect power to the CCI, and follow the directions in the CCI software documentation to measure the lane positions and create the configuration file.

### CCI flow cell design and fabrication

#### Overview

An exploded view of the CCI flow cell is shown below.

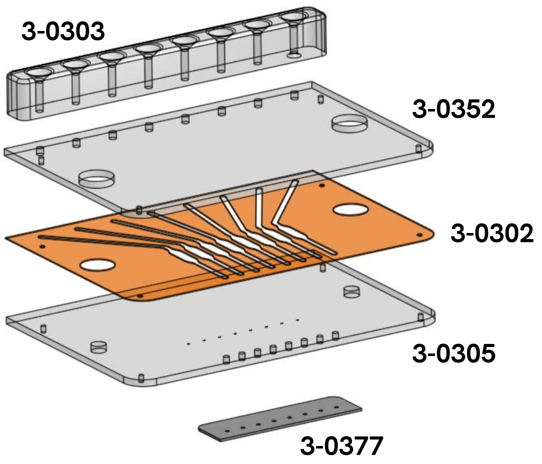

The core of the flow cell is a pair of PMMA plates (top plate, 3-0352; bottom plate, 3-0305) sandwiching an adhesive spacer layer (3-0302) which creates 8 fluid channels. A soft elastomer part on the top of the flow cell with chamfered openings (3-0303) forms a seal with the robot's pipette tips for injection of liquid. An etched PTFE strip with small exit holes covers the waste ports on the bottom of the flow cell and allows fluid to be pushed out by the pipette but inhibits wicking of fluid out of the channels which otherwise happens when the area around the waste ports becomes wet.

The flow cell was originally designed to be possible to fabricate without specialized equipment beyond a $CO_2$ laser cutter. Files used for cutting the flat parts can be found as attachments to the Cell Counting Imager Onshape document. The pipette tip interface was originally designed for "DIY" manufacturing as well but this was abandoned as discussed below.

#### Top (3-0352) and bottom (3-0305) plates

The top and bottom plates are cut from 1.5mm-thick cell cast PMMA material. The bottom plate has a set of etched fiducial marks which are used for image registration and for recognizing the center of the channel during manual calibration of stage positions.

#### Spacer layer (3-0302)

The "in house" version of the spacer layer was cut from Grace Bio-Labs SecureSeal adhesive sheet, 0.24mm thickness (SA-S-2L) using a $CO_2$ laser cutter.

#### Pipette tip interface (3-0303)

The pipette tip interface is cast from a Shore 40A hardness, water-clear polyurethane rubber material by an online prototype manufacturing service.

The pipette tip interface was originally produced in-house by casting PDMS in a custom mold made of machined aluminum plates and dowel pins but this was eventually abandoned due to the constraints it posed on the design of the part and the unreasonable amount of effort and mess involved required to produce the part this way compared to simply ordering elastomer castings from a prototype manufacturing service based on a solid model file.

The pipette tip interface is bonded to the flow cell using an adhesive gasket made from the same material as the spacer layer.

#### Waste nozzle strip (3-0377)

The waste nozzle strip is produced from 0.02" thick, adhesive-backed PTFE sheet (McMaster-Carr ) by etching an extremely fine checkerboard pattern before cutting out the drain holes and the outline of the part.

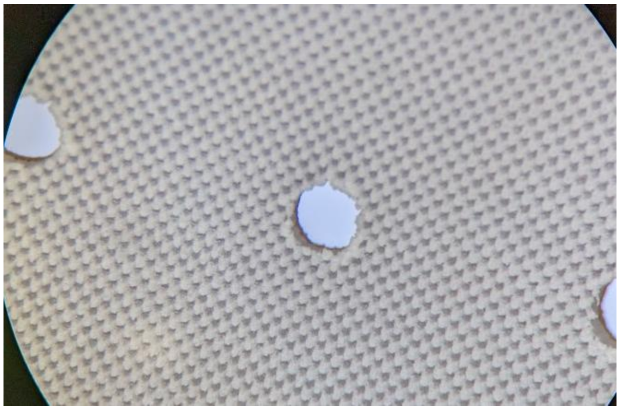

The combination of the small exit holes and the superhydrophobic laser-treated surface surrounding them are designed to force exiting liquid to break away and fall in small drops rather than wet the surface and form a wicking path into the waste trough or large hanging drops that pull liquid out of the channels.

### General operating procedures

This section contains a few key points about general handling of the instrument and flow cells. Users will need to develop their own detailed SOPs as appropriate to their situation.

#### Mounting consumables

The waste trough and flow cell are mounted on the CCI as illustrated below. The waste trough must be placed first as the flow cell blocks it from being removed. Both items can be left in place for multiple experiments if the waste trough is emptied by aspiration as necessary. After use the consumables should be removed to allow removal of any spilled liquid that may have seeped into gaps, and all items should be cleaned as described below.

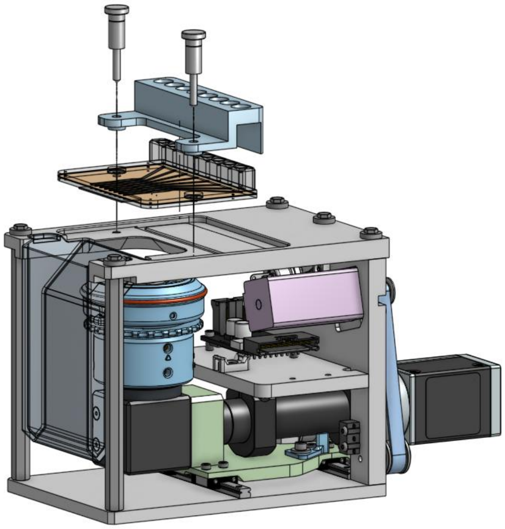

#### Automated operation

The CCI is normally mounted on the deck of a pipetting robot and controlled using simple HTTP requests. In the context of ACCS these requests come from the protocol script running on the robot. Refer to the CCI software documentation for more details.

Information pertaining to use of the CCI from the operator's perspective, in the context of an automated cell culture workflow, can be found in the *CCI Normalization Sample SOP*, available separately as a supplement to the ACCS manuscript.

#### Empirical calibration

Currently the CCI software uses a single calibration factor, supplied as part of the configuration file, to convert cell counts to concentration. This allows us to account for a variety of non-ideal factors (software counting sensitivity, manufacturing variation of the flow cells and the instrument itself, light source aging, etc.). Because of the relatively frequent turnover of flow cells and the labor involved in making a quality ground truth measurement, we do not re-calibrate for each individual flow cell, and the software currently does not use separate calibration values for each channel on the flow cell (though adding support for this would be relatively straightforward). In practice we find that any systematic inter-channel variation in readings for a given flow cell (e.g. due to variations in material thickness, channel geometry, reflective debris, etc) is generally obscured by other sources of noise and bias.

Our typical procedure to calibrate the CCI is to prepare linear dilutions of stocks of the relevant cell type, spanning the expected concentration range, then use repeatedly measure these with the CCI (with the help of the OT-2) as well as with a reference instrument, and use a linear regression to generate the calibration factor. For full details, refer to the procedure described in the ACCS manuscript and/or the Supplementary Materials and Methods, as well as the analysis code provided in the main repository for the publication. Of course, for other use cases, samples could be loaded manually and the CCI scans performed with (e.g.) a simple Python script.

The CCI software distribution includes an example protocol script for the Opentrons OT-2 to load a series of samples for CCI calibration measurements. The default CCI configuration also includes a generic calibration factor that should be "close enough" for basic use if a reliable reference instrument or appropriate cell stock is not available for calibration purposes.

#### Quality checks

When ACCS is being used in production, we perform a weekly Monday morning quality check by preparing a cell dilution with a particular nominal density (e.g. 250k cells/mL) and running a short OT-2 script that loads the same stock and takes measurements with the CCI several times in a row. The stock is prepared by dissociating HEK293T cells grown over the weekend, taking a concentration measurement with a Countess II cell counter, and diluting accordingly. We plot the resulting data and flag any statistical anomalies based on arbitrary alert thresholds, e.g. any outliers not explained by air bubbles picked up by the pipette, >12% CV between channels on a given scan, >15% CV across cumulative readings from all 6 measurements, or an average measured concentration further than ±25% from the expected value.

The CCI software distribution includes an example protocol script to run the aforementioned QC measurement protocol.

#### Flow cell storage and working life

No special storage conditions are observed for new, unused flow cells or flow cell components.

Once a flow cell has been exposed to biological materials it is kept in a dated plastic container (e.g. a 150mm round culture dish) under refrigeration. Used flow cells are discarded after at most 31 calendar days as a proactive contamination control measure. We therefore do not have an empirical basis to predict a practical maximum service life for a CCI flow cell.

#### Flow cell cleaning

The exterior surfaces are wet cleaned before and after use with a commercial plastic cleaning product (Blue Ribbon Plexi-Clean) and soft polyester cleanroom wipes (Texsorb Absorbond TX404). Exposing the PMMA surfaces to alcohols should be avoided as it induces stresses in the material and can cause crazing under some conditions.

Cleaning of the internal channels is an automated process built into the ACCS protocols. The robot fills the flow cell with a strong detergent solution (Decon Laboratories Contrad 70, diluted to 10% v/v in DI water) and soaks it for 60 seconds, then flushes three times with 200 µL of water.

#### Cleaning the CCI instrument

The open-frame design of the CCI poses some limitations on cleaning methods. It should not be sprayed down with liquid. Exposure to incidental mists of alcohol solution while cleaning the biosafety cabinet and robot deck is likely to be harmless.

We normally soak a laboratory wipe in 70% ethanol and spot clean the top of the instrument as well as any surfaces exposed to drips or spills. Cleaning the linear guide rails should be avoided if possible. In the case of a spill reaching the guide bearings, proactive replacement of the affected linear guide should be considered as there is not a practical way to decontaminate the internals and it may eventually start to fail.

The waste trough is normally disinfected with a 10% bleach solution. The trough should be emptied of bleach solution and allowed to dry before mounting it back on the CCI.
