## Supplementary material for "Open-source cell culture automation system with integrated cell counting for passaging microplate cultures": D. Sample CCI Normalization SOP

### ACCS CCI Normalization SOP

*Uses Opentrons software version 4.7, protocol framework* `t220929a` *or later and the CCI2.5 hardware.*
*DO NOT ACCEPT ANY DIALOG OFFERING TO UPGRADE THE OPENTRONS APP OR ROBOT SOFTWARE.*

*Protocol Description:*
This protocol takes a 96 well source plate containing adherent cells and splits it out to a propagation plate, using the CCI to normalize to specified seeding targets (cells/well) for each well. Optionally it then distributes cells to a second plate at the same density (but optionally a different amount). Each destination plate can be either a standard 360uL 96-well plate or a 440uL glass-bottom imaging plate. By default a single plate will be filled to 200uL or two plates to 150uL each.

1. Initial prep
    a. Start thawing/warming up reagents to 37°C
        i. At least 8mL of 0.25% trypsin-EDTA dissociation reagent
        ii. At least 60mL of suitable growth medium including 10% FBS and 25mM HEPES
    b. Open BSC sash and start fan
    c. Wipe down surfaces in the BSC
    d. Take a dish with a flow cell from the refrigerator and move it to the BSC (it needs a while to warm up to avoid condensation)
    e. Install tip trash liner:
        i. Cut new bag to half height and discard upper half
        ii. Tuck bag into bin and secure with binder clips (you can take the bin off the deck to make this easier). Make the bag as flat as possible against the floor and inside walls of the bin, especially around the front-left corner. Make sure the overhanging plastic on the outside is kept neat and does not encroach over neighboring deck slots (see example below).

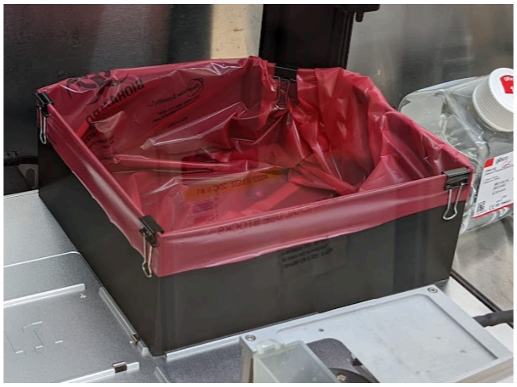

        iii. Put tip trash bin back in place on robot deck

2. <u>CCI Prep:</u>

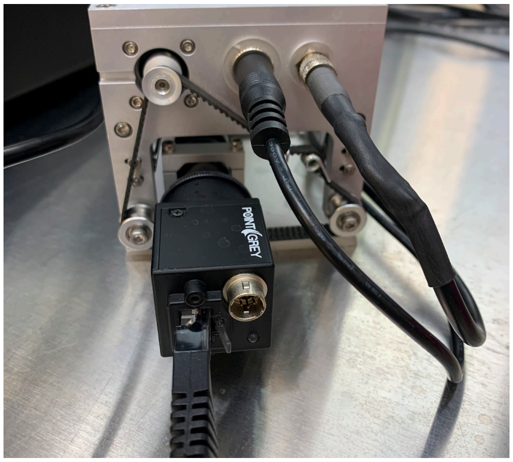

    a. If the CCI is not in place already, install it on Slot 6 of the OT2, with the waste cup on the left side and the connectors on the right.
    b. Make sure the three cables to the CCI are plugged in as shown
    c. Make sure the ethernet cable plugged into the camera is not obstructed by the other cables and has room to move back and forth

3. <u>Flow cell prep:</u>
    a. If you are using a brand new flow cell, make sure it has received the Pluronic F-127 pre-treatment as described in the Cell Counting Imager Technical Manual.
    b. Check the flow cell for debris and/or residue on both sides, particularly over the wide part of each channel (where imaging takes place).
        i. Clean with Plexi-Clean and a lint-free cloth as necessary. Do not use alcohol.
        ii. Use compressed air as necessary to blow off any dust and/or lint left after wet cleaning.
        iii. If the flow cell is still cold from the refrigerator it may form condensation. Defog the surface with a gentle stream of compressed air if necessary.
    c. Put the flow cell in place on the CCI
        i. Handle the flow cell by the edges as much as possible and avoid touching the top and bottom surfaces near the imaging area
        ii. Place the flow cell with the mounting holes lined up with the holes on the CCI deck and drop it into place.
        iii. Push the flow cell to the left (towards the waste trough) to make sure the rounded end is pushed up against the wall.
        Push from the mounting holes (again, avoid touching the imaging area).
        *iv. Do not install the tip guide and thumbscrews yet!*
    d. Clear the flow cell channels of liquid
        i. Place a cloth over the opening of the waste cup on the left side of the CCI to mitigate flying droplets
        ii. Line the nozzle of the compressed air gun up to each of the holes on the clear rubber tip interface on the flow cell and apply just enough flow to blow out any liquid from storage or the previous run. A slight fog or micro-droplets may remain on the inside surfaces.

4. Robot startup
    a. Turn on the robot, tempdeck, CCI camera and CCI stage using their designated switches on the outside of the BSC.
    b. If the robot was not turned on already, it will take a few minutes to start up. When the robot is ready the blue light will stop flashing and you will a few mechanical noises.
    c. Open the Opentrons app. Connect to the robot by clicking on the toggle next to the relevant robot name (e.g. OT2-SWEETPEA). Make sure you select the correct robot! Each robot has a name label attached on the front beneath the logo.

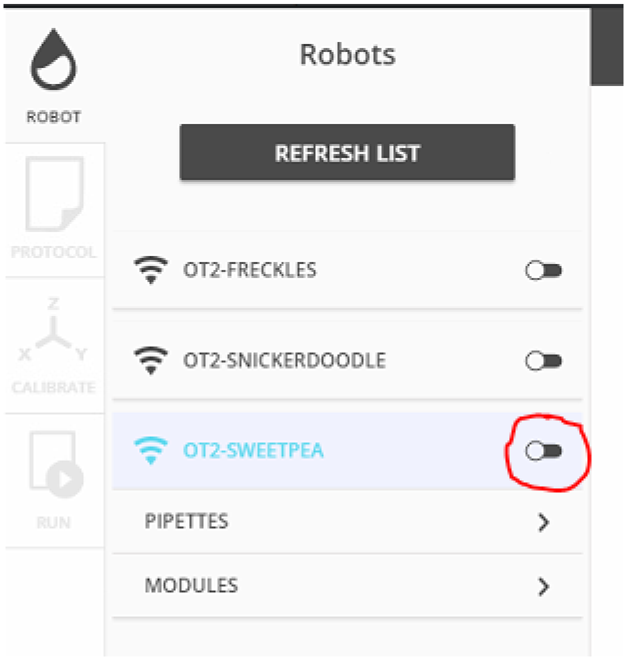

5. Prepare the protocol script
   a. Open the "Splitting software" folder on the desktop, then "Protocol setup" and then finally run "Setup CCI Normalization (NPF)". A browser window should pop up and show a Jupyter notebook called "CCI Normalization Protocol Setup".
   b. Click the ⏩ button as instructed to initialize the form
   c. Fill out the form fields
   d. Click the "Generate script" button and the path to the generated protocol script should appear along with a confirmation message.

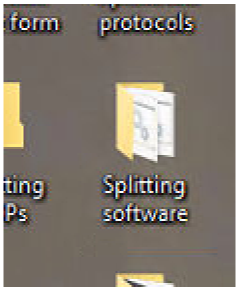

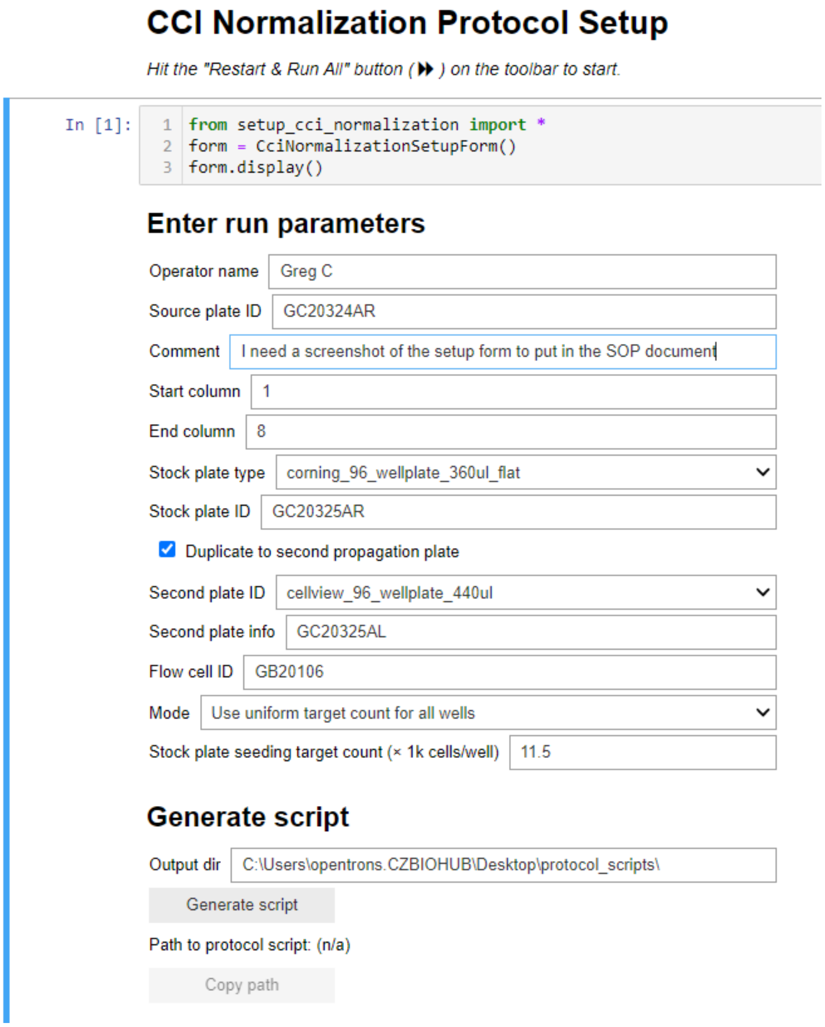

6. (if needed) Labware calibration

This does not need to be done for every run! Skip this section unless the robot has been moved or there is some other reason to believe the calibration needs to be checked/adjusted.

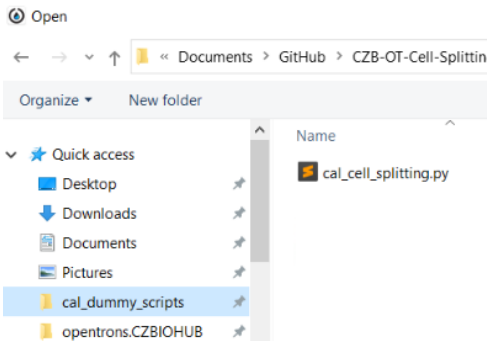

   a. In the Opentrons app, open the protocol called "`cal_cell_splitting.py`". You can find it in the dialog via the "cal_dummy_scripts" shortcut in the Quick Access pane.
   b. Select the "Calibration" tab in the app, ensure the tempdeck is powered up, and click "Continue to labware setup" if shown
   c. Set up labware for calibration
      i. Refer to the deck layout on the "Calibration" tab of the Opentrons app and install labware as applicable.
         - See the "Tip for installing labware" inset below this section if you are unfamiliar with attaching labware to the OT-2 deck.
         - You can use spare / nonsterile labware (96 well plates and 12-well reservoir) for the calibration process.
         - Only install the labware types you are actually using (i.e. if you are not using a microscopy plate, you don't need to put anything in slot 1).
         - Put each applicable type of labware in the specific location indicated, regardless of what arrangement of plates you are using for your actual run
         - Do not place your source plate on the tempdeck yet.

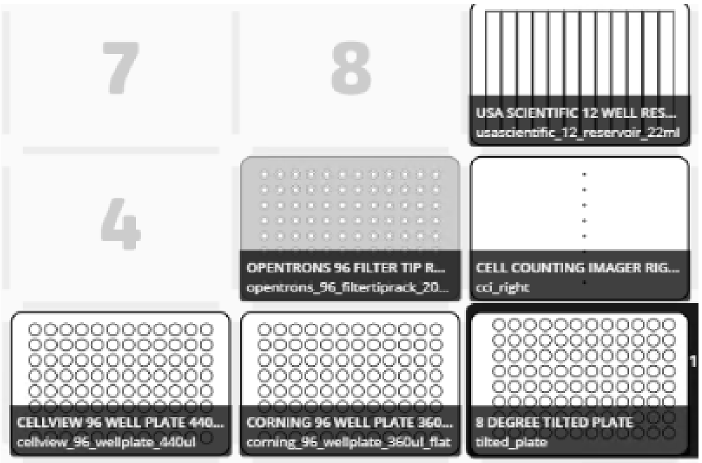

      ii. Install a box of tips on slot 5 for calibration. You only need the leftmost column filled with tips. It is recommended to use a leftover open box for this.
      iii. Install the tilted plate calibration tool on the tempdeck.

d. Review the following notes, then follow the steps on the Calibration tab of the Opentrons app to proceed through labware calibration.
   Note: After the tip box is done, you can choose the order to calibrate labware in by manually selecting from the deck diagram instead of using the "next" button. It may be beneficial to go in the order listed below.
   i. Tip box (slot 5)
      - Do not change the calibration unless the alignment is visibly off and/or tip pickup fails. Pick up tips, save, and continue.
   ii. Trough (slot 9):
      - Make sure the tips are roughly centered over the first well of the reservoir. The ends should be level with the top surface of the labware and not below it.
   iii. CCI (slot 6):
      - Make sure the flow cell is seated over the window on the CCI.
        The black tip guide should not be in place yet!
      - The ends of the tips should be (on average) centered in the entry cones and should dip about 1/8" below the top surface of the pipette tip interface, down to the bottom of the entry cones where the narrow channels start.
   iv. Tilted plate (slot 3)
      - Verify that the tilted plate calibration tool is in place on the heat block before sending the robot to that slot.
      - Z position should be such that the ends of the tips are in plane with the top surface of the tool or a little above -- verify by looking with your eyes level with the top of the tool (see photo). Ensure the tips are not down inside the groove at all; err on the "too high" side.

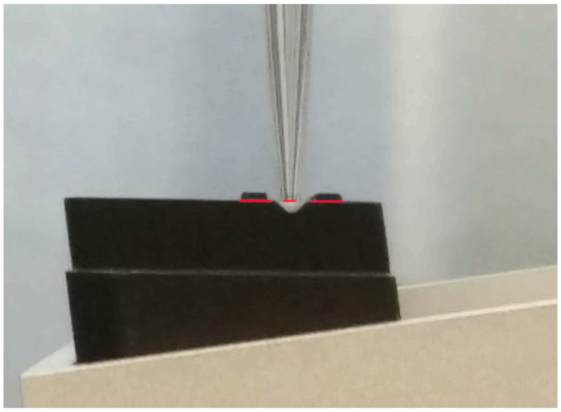

      - X and Y position should be such that the tips *(on average)* line up with the raised crosshairs. It will never be perfect; it is normal for all 8 tips to be *slightly* off in different directions.

v. Destination plate(s) (slots 1 and 2)
- You will probably want to use an identical spare plate for calibration instead of your sterile propagation plate
- Make sure there is no lid on the plate before sending the robot to that slot
- If you will not be using the type of plate assigned to a given slot, you can just omit the plate, accept the calibration and move on without changing it.
- Center the tips on the wells, making sure the ends are level with the top rims and not poking down into the wells.

e. After you have calibrated every item, click "Return tips and proceed to run" (but do not actually proceed to run yet).

f. Remove the tilted plate calibration tool and any non-sterile labware used for calibration. Wipe down the deck again if appropriate before proceeding to install your sterile labware.

**Tip for installing labware**

1. Put the front right corner of the labware against the deck
2. Push it towards you and to the right until it fully compresses the spring clips

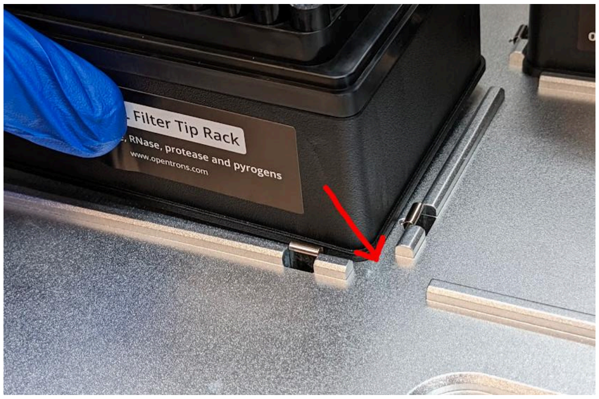

3. While holding the labware against the clips, push it down flat to seat it
4. Release and visually confirm proper position. Rails should be visible on all 4 sides.

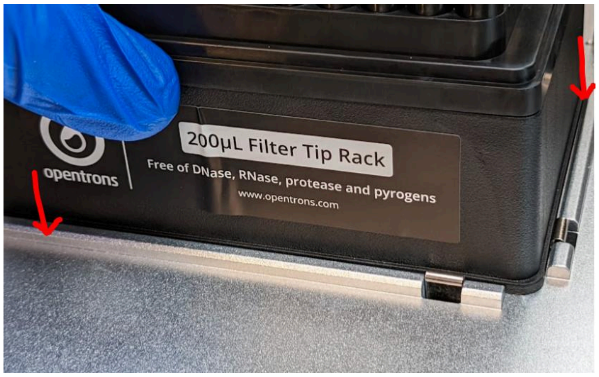

7. Set Up
   a. Go to the Run tab in the app and set the tempdeck to 41°C so it has time to stabilize
   b. In the Opentrons app, open the protocol script you generated in the previous step. The simulation process will run while you complete the rest of the preparations.
   c. Load reagents in trough in the arrangement shown below. Consider adding the trypsin last.

| | | | | | | | | | |
|---|---|---|---|---|---|---|---|---|---|
| 1. Waste (empty) | 2. Waste (empty) | 3. PBS (20mL) | 4. PBS (20mL) | 5. Trypsin (8mL) | 6. Media (20mL) | 7. Media (20mL) | 8. Media (20mL) | 9. DI water (8mL) | 10. Cleaning soln* (4mL) |

*10% Contrad 70 in distilled water. Provided in a tube labeled "CCI CLEANING SOLN".

   d. Ensure your source plate contains no more than 200µL of media per well. Remove some liquid if necessary.
   e. Finish loading labware on the deck.
      i. You can check the Calibration tab in the Opentrons app to determine how many tip boxes to install and where. Install remaining labware according to the layout below.
      ii. When mounting your source plate on the tempdeck, push down to make sure it is fully seated. The fit may be snug on some plates.
      iii. When loading tip boxes, go in order from right to left and back to front, making sure they are properly seated inside the rails, not overhanging or offset.

| | | |
|---|---|---|
| 10 Tip box? | 11 Tip box | (Trash) |
| 7 Tip box? | 8 Tip box? | 9 Reagent trough |
| 4 Tip box? | 5 Tip box | 6 CCI |
| 1 Propagation plate 2? | 2 Propagation plate 1 | 3 Source plate (on tempdeck) |

8. CCI startup
    a. Install the pipette tip guide onto the flow cell and fasten it to the CCI with the two thumbscrews. Make sure to keep turning the thumbscrews until you feel a positive stop. It is important for the flow cell to be seated properly.

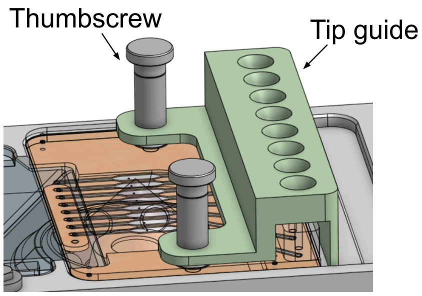

    b. Make sure there is not already a "CCI Server" console window open!
    c. From the "Cell Splitting" folder on the desktop, double-click on "Run CCI".
        i. A console window titled "CCI Server" will open. You should eventually see something like this:

```
======== Running on http://0.0.0.0:80 ========
(Press CTRL+C to quit)
```

        ii. If you see an error instead, seek assistance from a support contact before proceeding
    d. Turn off the light in the BSC

9. Final check

   Always double-check before starting the robot:

   a. Did you remove the tilted plate calibration tool and install your source plate?
   b. Did you replace the labware used for calibration on slot(s) 1/2 with your actual sterile plate(s)?
   c. Are the tip boxes and trough aligned and seated properly in their slots?
   d. Are the lids off of all plates and tip boxes?
   e. Do you have the CCI software running?

10. Run the protocol

    a. Go to the Run tab in the Opentrons app and start the protocol
    b. Make sure you have joined the relevant Slack channel (e.g. `#ot2_error_sweetpea`) to receive notifications
    c. IMPORTANT:
       Check on the tip waste bin occasionally to make sure tips are not stacking up
       i. You can use an aspirating pipette or other convenient object to gently rake the tip pile to the rear and right sides of the bin, away from the "drop zone".
          Be very careful not to knock or launch tips out of the box in the process, and be aware of the robot's movements. It may be prudent to pause the robot.
    d. IMPORTANT:
       If you are splitting to two destination plates, you will need to replace the tip box on Slot 5 once all the tips in it have been used.
       i. Pause the protocol from the Opentrons app (it is recommended to wait until the robot goes to the trash to discard a tip). You can proceed without pausing the robot but this is at your own risk.
       ii. You may want to cover your destination plate(s) before reaching over them to swap the tip box
       iii. Make sure the new tip box is properly seated and you haven't bumped any neighboring boxes and unseated them

11. Clean up
    a. Bleach source plate
    b. Incubate propagation plate(s)
    c. Consolidate leftover tips in boxes marked "non-sterile" and save for future use. Recycle empty boxes and extra lids.
    d. Aspirate all liquid in the reagent trough and dispose of it
    e. Empty and clean the CCI waste cup
        i. You can free the waste cup from the CCI by lifting up and then sliding it out to the left
        ii. Aspirate all liquid contents
        iii. Disinfect by washing down the inside with 30% bleach solution and then aspirating the excess
    f. Put the flow cell away
        i. Undo the thumbscrews, remove the tip guide, then remove the flow cell from the CCI
        ii. Dab away any excess liquid on the outside of the flow cell with a Kimwipe, then place the flow cell back in its dish, upside-down with the orange drain gasket facing up.
        iii. Add a tally mark to the lid of the dish and place it back in the fridge.
    g. Clean up the CCI
        i. Remove any excess liquid on the top plate of the CCI, then wipe it down with a Kimwipe moistened with 70% ethanol. Avoid letting liquid get on the objective and do not spray anything directly onto/into the instrument.
        ii. Cover the CCI by placing a tipbox lid on top of it
    h. Unclip the liner from the trash bin and dispose of the used tips. Check around the work surface and robot deck for any loose tips.
    i. Disinfect the OT2 deck, trash bin and BSC work surface with 70% ethanol. Avoid spraying liquid on the CCI or into the air vents on the tempdeck.
    j. Disinfect the CCI tip guide, thumbscrews and trash liner clips with 70% ethanol and place back in the storage dish
    k. Shut the BSC sash
    l. Turn off the CCI, OT2 and tempdeck from the switches on the power strip outside the BSC
