## Supplementary material for "Open-source cell culture automation system with integrated cell counting for passaging microplate cultures": E. Growth and Viability

### Supplement E: Further investigation of effects on growth and viability

#### Abstract

Here we further explore potential impacts on cell viability and morphology from passaging cells using ACCS as compared to manual methods, including under extended incubation periods of up to 3 days. We seeded sets of plates with a fixed splitting ratio, both by hand and with the robot, then performed ATP viability assays at approximately 1, 2 and 3 days from seeding. For each condition we also captured 10X phase contrast images from preselected random sites on the plate before the terminal assay to provide examples of morphology. We also investigate possible impacts of the timing of seeding individual wells within a given ACCS passaging run.

#### Results and discussion

##### Viability over various incubation intervals

We measured total ATP to assess overall viability of plates seeded by the robot after incubation periods of 1, 2 and 3 days, as compared to plates seeded manually. For the robot-passaged plates we observe well-to-well variation similar to or better than their manually passaged counterparts and no detriment to viability attributable to handling by the robot.

Figure S3 illustrates CellTiter-Glo 2.0 assay results obtained from test plates at roughly +1, +2 and +3 days since seeding. The test plates were seeded with a fixed 1:6 split factor from nominally uniform source plates and cultured in parallel. Each point on the plot is the mean value over 76 test wells, except that 10 wells are excluded from the second manually-seeded plate in Fig. S3(A) due to the corresponding row on the hand-seeded source plate being inadvertently double-seeded. Figure S4 shows the distribution of scaled assay values from each read.

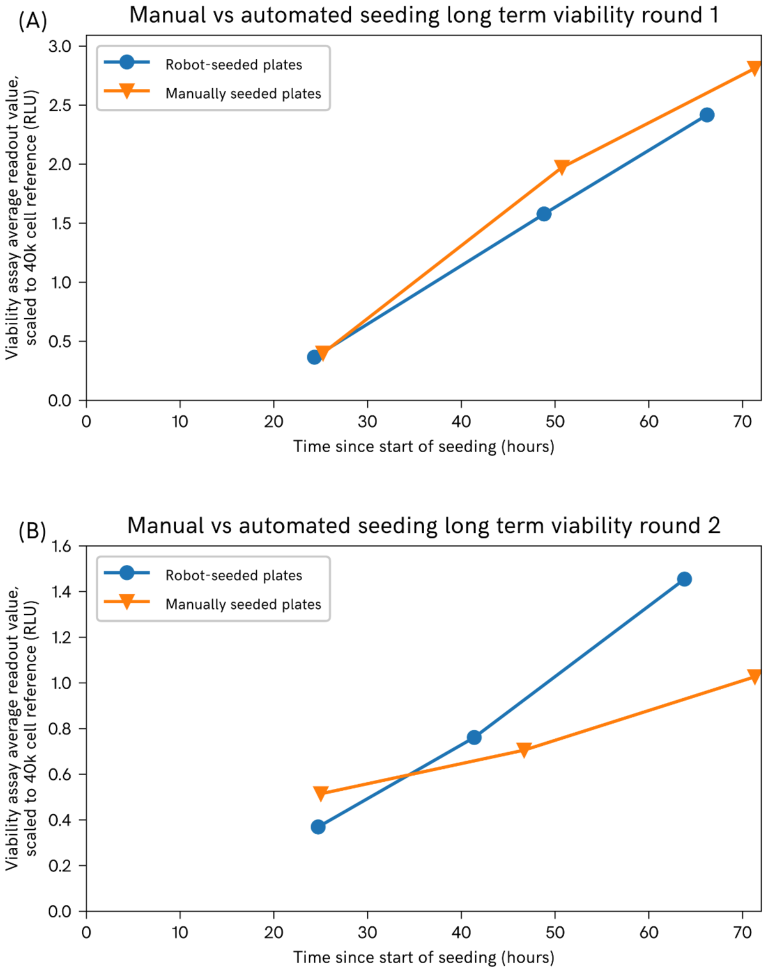


**Fig. S3.** Average of ATP assay readout values for each test plate, scaled to a reference sample on each plate with 40k cells/well, plotted against age of the plate since seeding. To be clear, the assay is terminal and each point represents a different plate. Panels **(A)** and **(B)** represent experiments carried out on separate weeks.

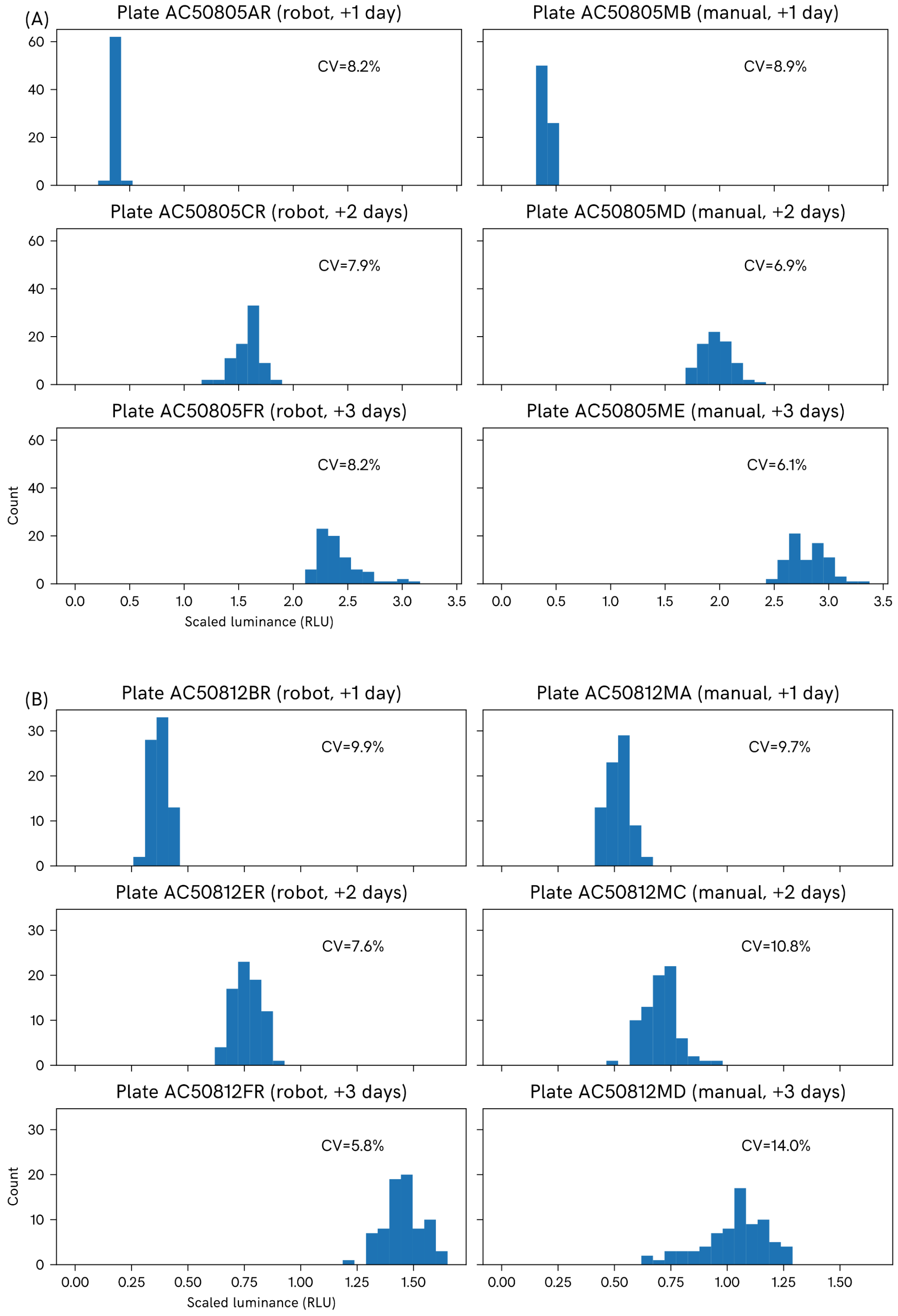


**Fig. S4.** Distribution of individual well values for each test plate. Subfigures (A) and (B) correspond to the data in **Fig. S3(A)** ("round 1") and **S3(B)** ("round 2") respectively.

### Effects of seeding order

Using the same datasets we also investigated whether the order wells are seeded in by the robot affects recovery and growth (wells seeded earlier in the process). We used the ACCS protocol logs to determine the time when each well on a plate was seeded and analyzed the assay scores against well seeding time.

As illustrated in Figure S5, on 5 of the 6 plates we observe a possible positive trend in viability scores with respect to the time the well was seeded. However, the effect is very small compared to other, uncorrelated variation in the data, and all but disappears at +2 days and beyond (in fact, in one case the trend is actually slightly negative).

Assuming we are not being confounded by some other, downstream phenomenon correlated to position on the plate, we speculate that such an effect could be a consequence of cells being subjected to varying lengths of ambient exposure immediately post-seeding according to their position on the plate and the order the wells are seeded in.

In our use cases we find a systematic effect of this magnitude is of virtually no practical significance and makes up a small fraction of the overall error. However, we also note that it would be straightforward to decouple seeding time from well location by adding a randomization strategy to the protocol script if desired. Further, any effect found to be due to temperature stress could be mitigated by adding a second heat block to keep the destination plate contents at an appropriate temperature throughout the protocol.

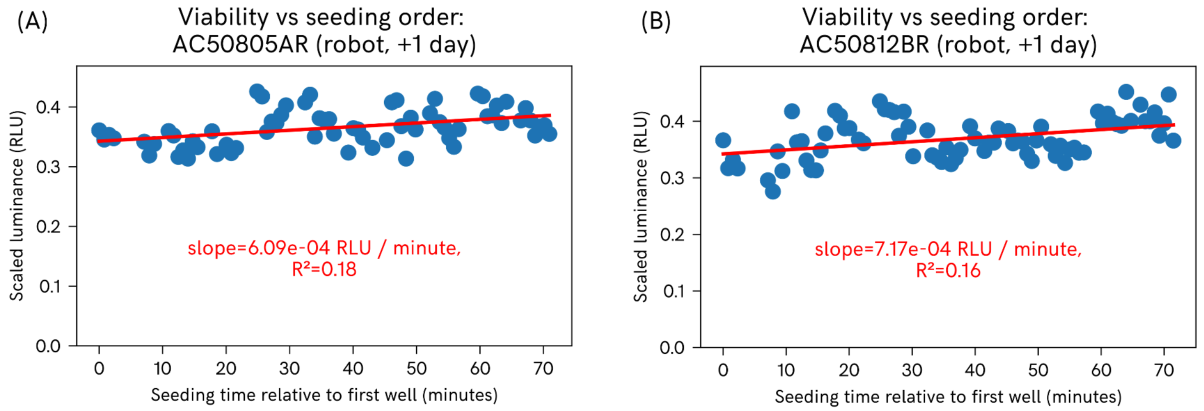

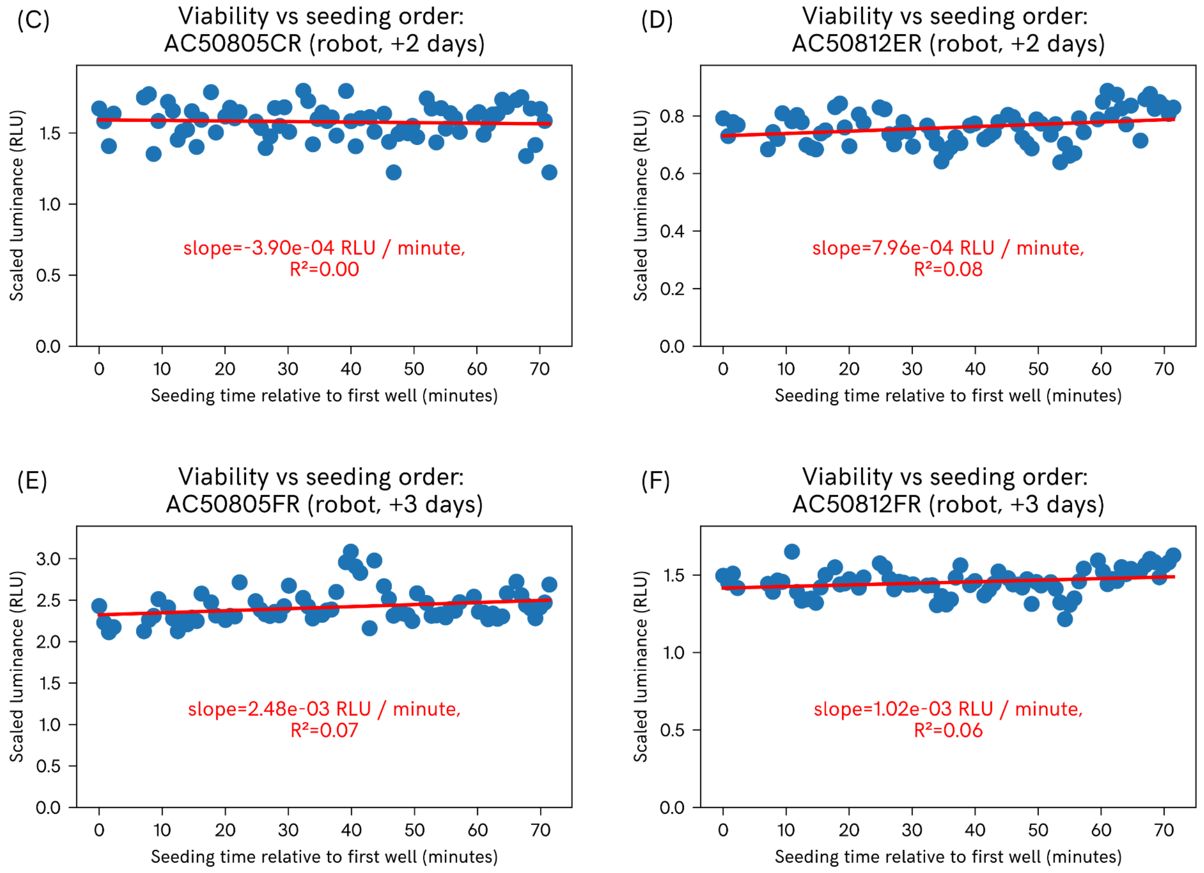


**Fig. S5.** Viability assay results plotted with respect to relative point in time when each well was seeded, with fit lines obtained by linear regression. Assay values are scaled to a reference sample on each plate with a suspension of 40k cells / well.

### Morphology images

This section presents a set of representative morphology images taken at 10X. The location references and imaging procedure are explained in the Materials and Methods section.
Overall we observe healthy cell growth and no systematic abnormalities in the robot-passaged plates compared to the manually-passaged plates.
Full size images and original raw files are available to download from the public repository:
https://github.com/czbiohub-sf/2024-accs-pub/accs24-pub-aux-data/supplement_e/morphology_pics.

#### 1-day-old plate

| | Robot-split plate | Manually split plate |
|---|---|---|

| | | |
|---|---|---|
| B2 upper middle | 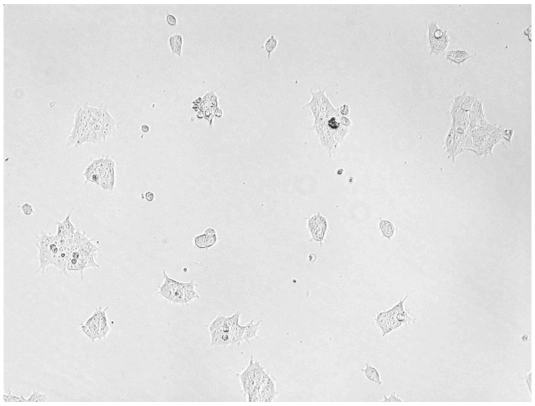 | 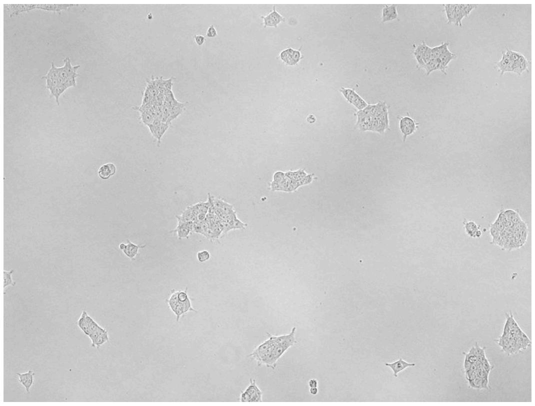 |
| C9 lower middle | 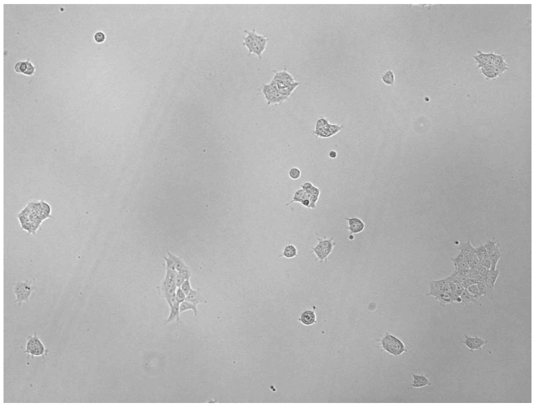 | 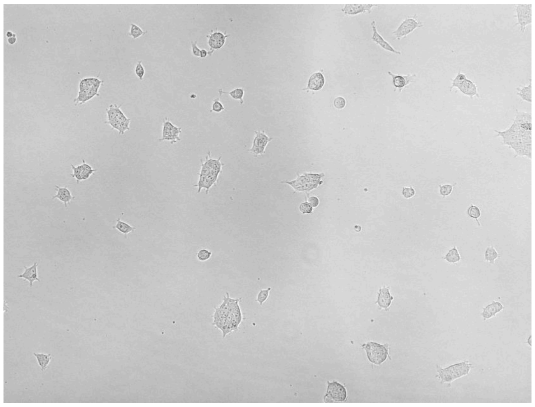 |
| E8 lower right | 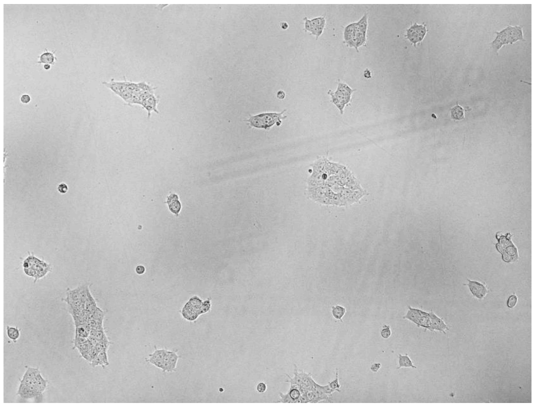 | 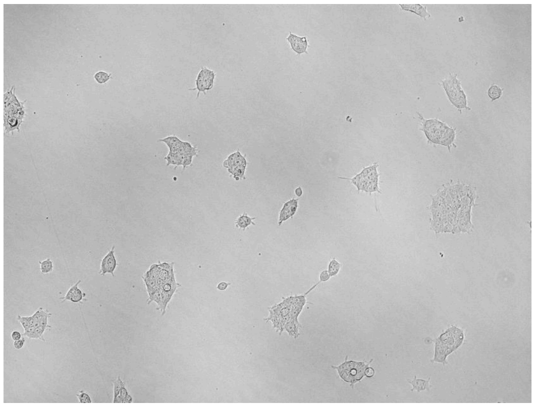 |

| | | |
|---|---|---|
| F8 right middle | 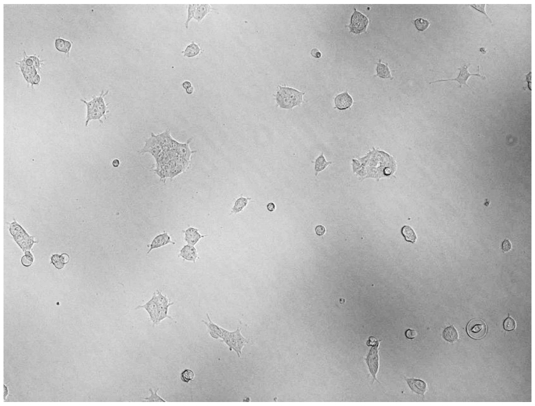 | 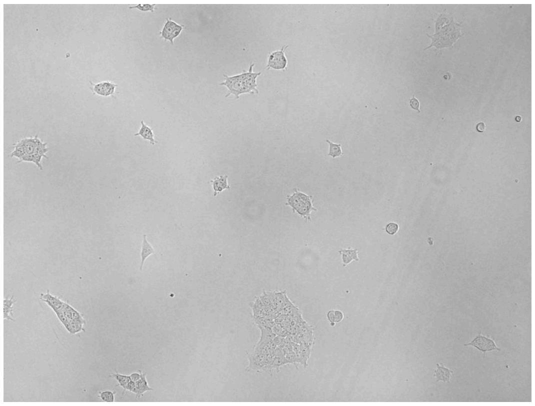 |
| G3 center | 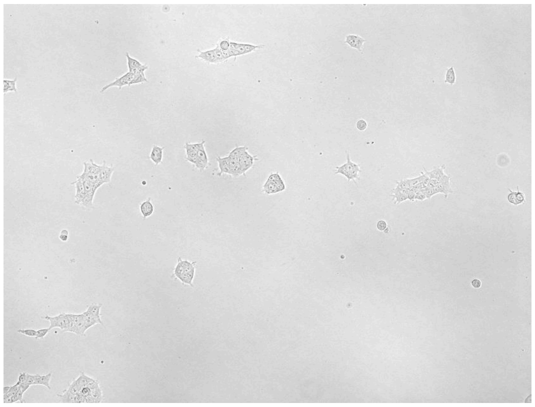 | 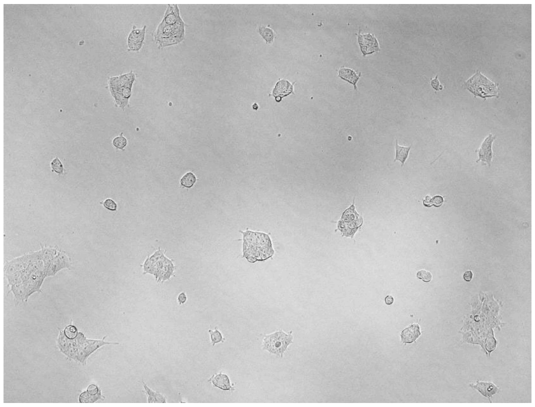 |
| G4 upper left | 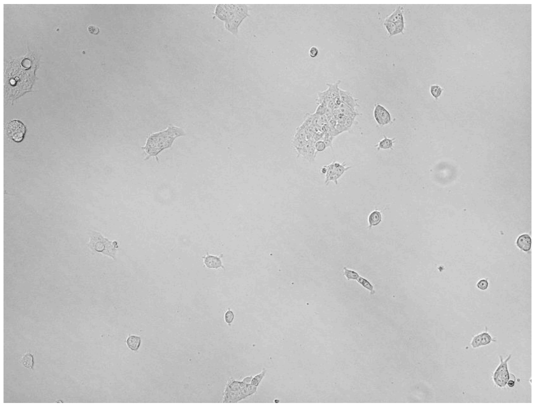 | 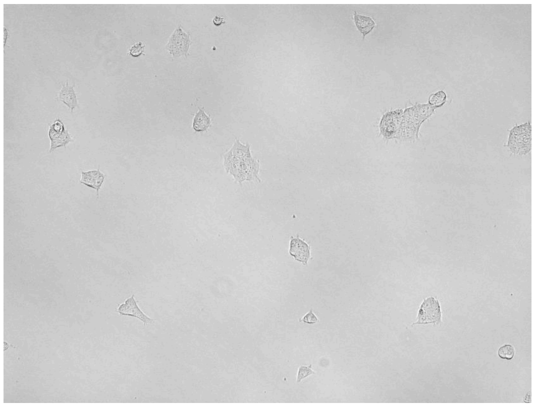 |

#### 2-day-old plate

| | Robot-split plate | Manually split plate |
|---|---|---|

| | | |
|---|---|---|
| B2 upper middle | 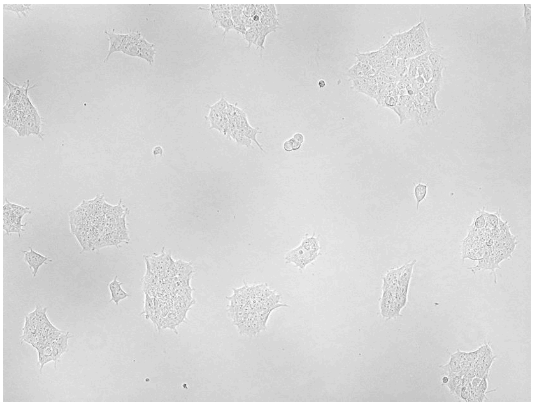 | 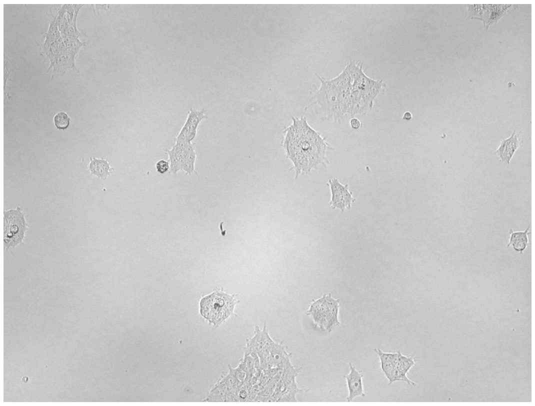 |
| C9 lower middle | 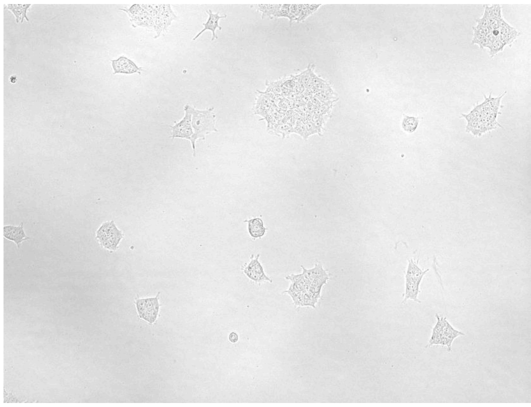 | 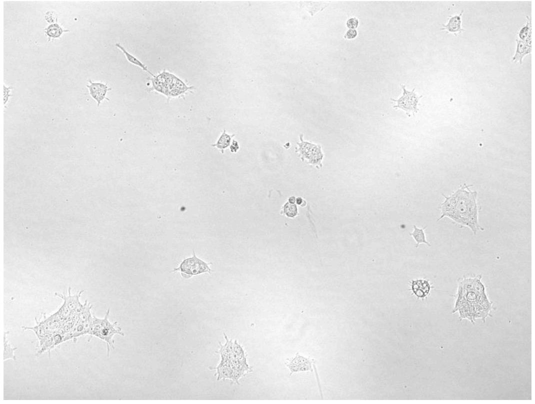 |
| E8 lower right | 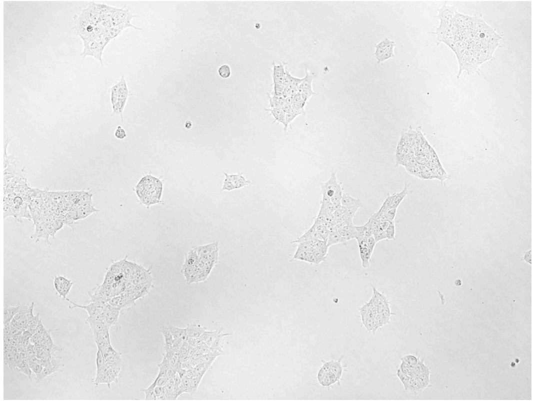 | 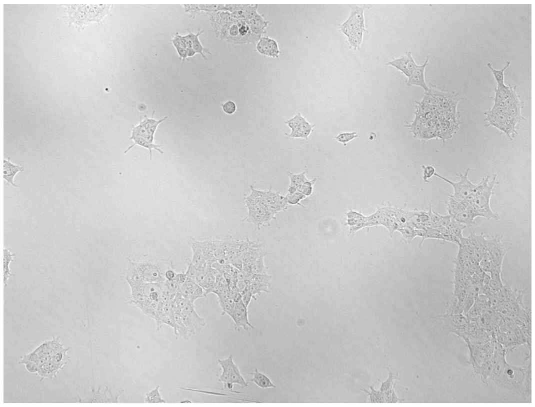 |

| | | |
|---|---|---|
| F8 right middle | 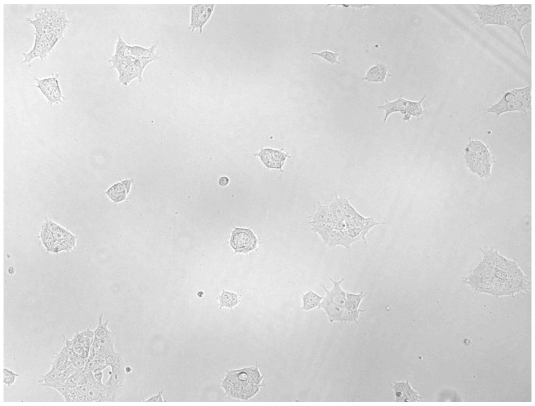 | 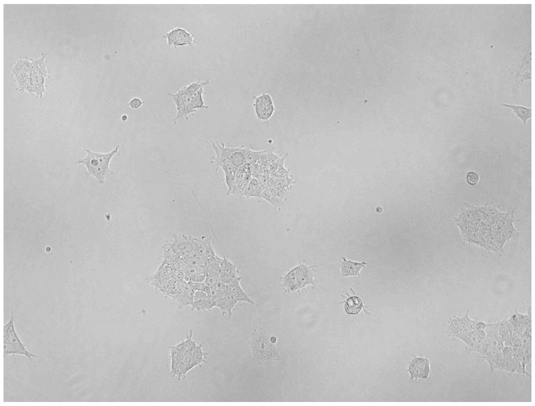 |
| G3 center | 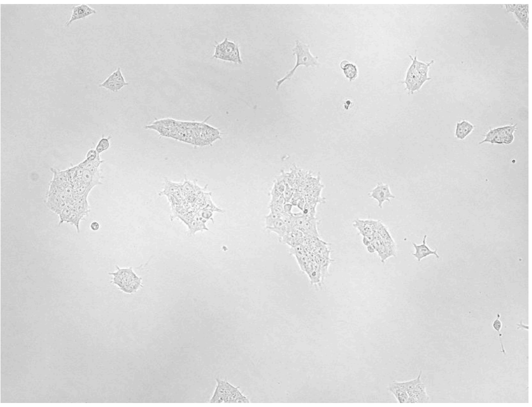 | 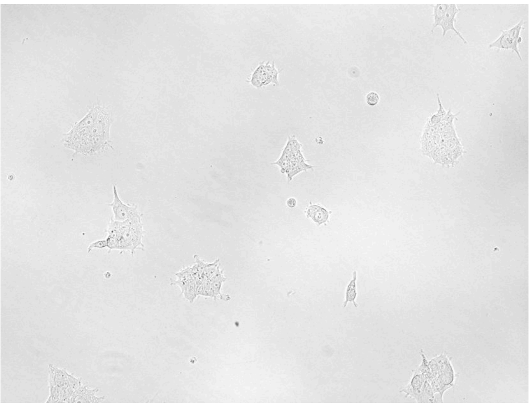 |
| G4 upper left | 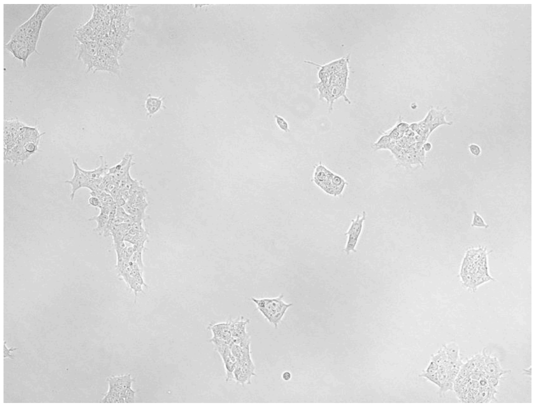 | 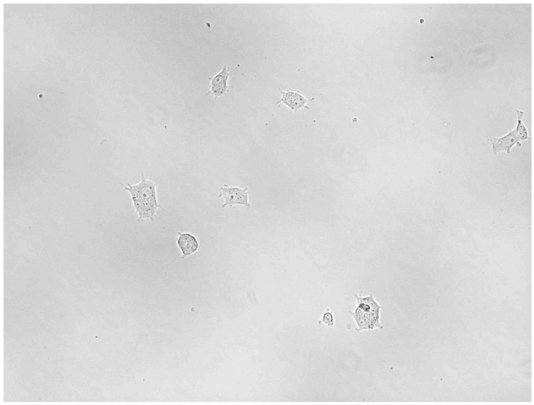 |

#### 3-day-old plate

| | Robot-split plate | Manually split plate |
|---|---|---|

| | | |
|---|---|---|
| B2 upper middle | 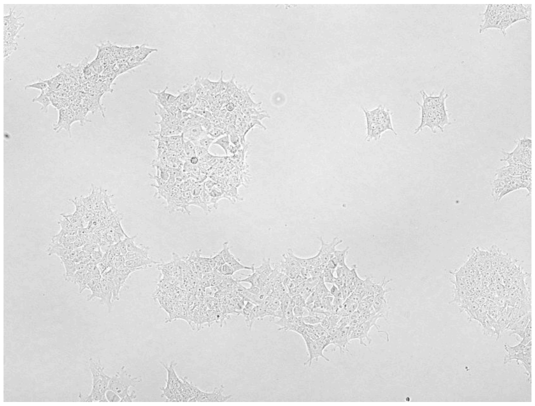 | 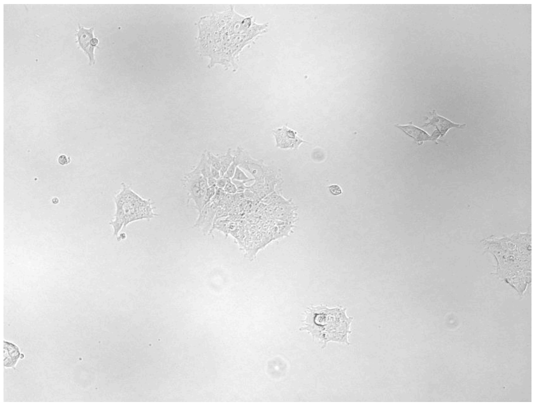 |
| C9 lower middle | 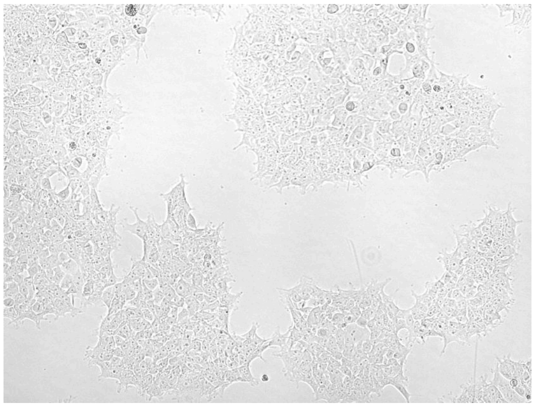 | 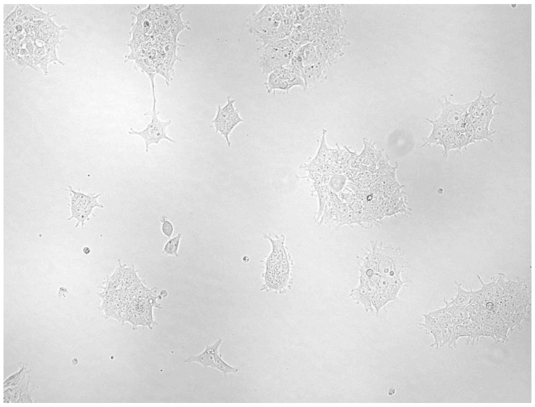 |
| E8 lower right | 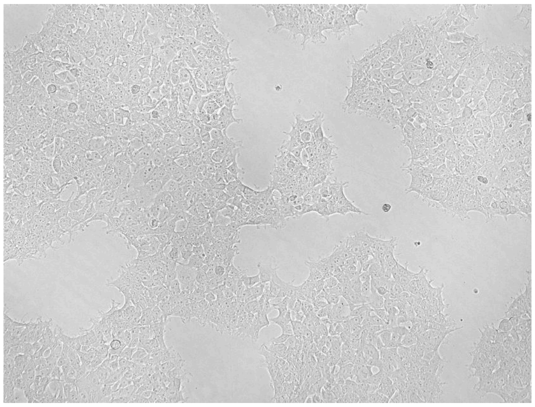 | 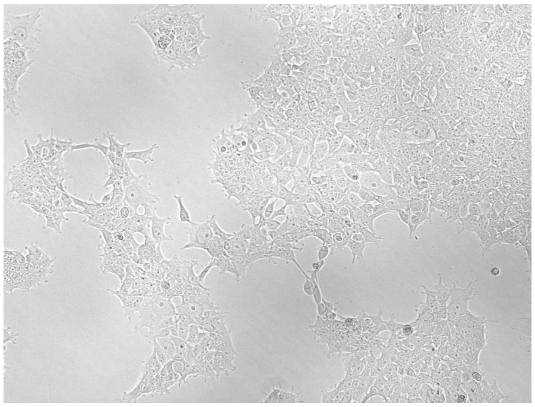 |

| | | |
|---|---|---|
| F8 right middle | 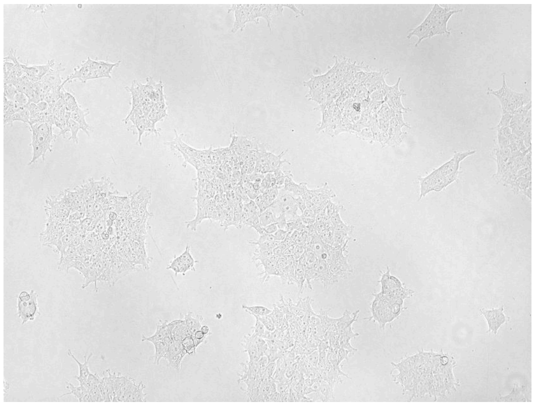 | 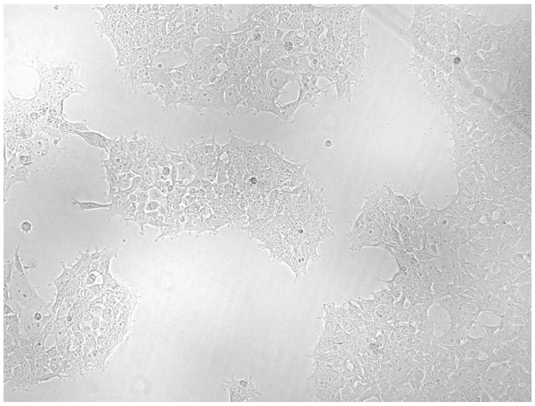 |
| G3 center | 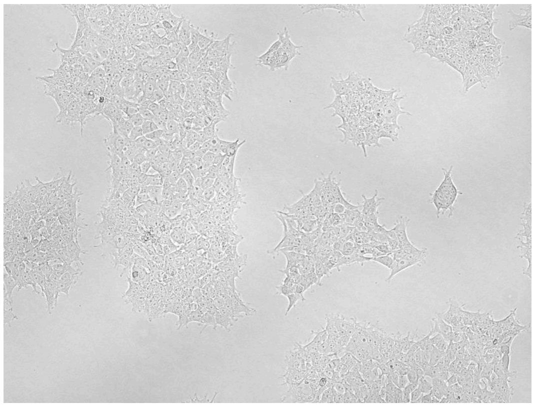 | 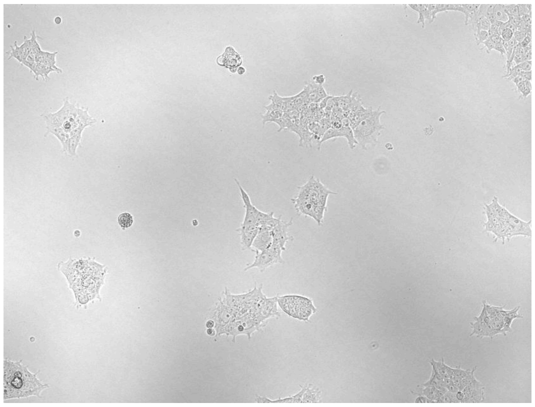 |
| G4 upper left | 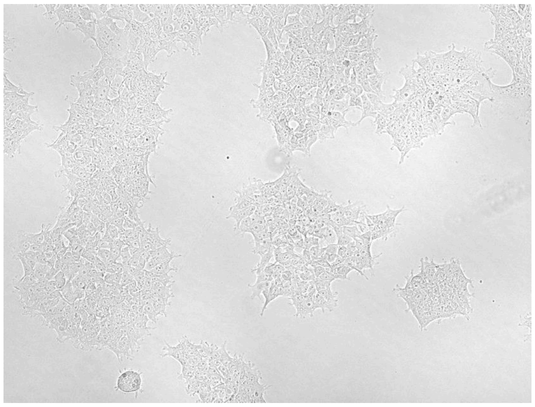 | 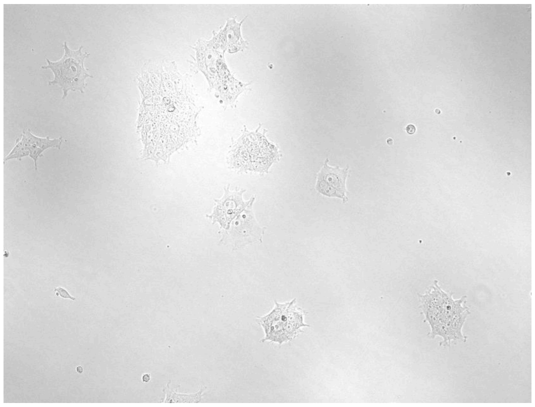 |

### Materials and Methods

#### Reagents and cultures

Cells (HEK293T) and media used for this experiment are the same as described in the main text, Materials and Methods, subheading "Reagents and cultures".

#### Instruments

ACCS system, as described elsewhere in this publication, without Cell Counting Imager
Spectramax i3x plate reader with Minimax 300 Imaging Cytometer
EVOS FL digital microscope, using 10X objective
Invitrogen Countess II FL automated cell counter

#### Consumables

Corning 3610 white wall, clear bottom, TC-treated 96-well culture plates
Bio-Rad Microseal 'B' self-adhesive optical plate seals
Thermo Scientific #236272 white vinyl plate bottom seals
Invitrogen # C10228 Countess disposable counting slides

#### Test plates

On Monday, 6 source plates for the test plates are prepared by manually seeding 30k cells/well into all 96 wells of a Corning 3610 plate. Stocks for seeding are prepared by diluting a stock of freshly dissociated HEK293T cells according to a concentration measurement obtained with an Invitrogen Countess II cell counter.
On Tuesday, 6 test plates are made by passaging the source plates from Monday with a fixed split ratio of 1/6. Specific wells are left empty: C3, C7, F3, F7 for negative controls; columns 5 and 10 for reference wells.
3 of the source plates are seeded by hand according to the general manual tissue culture practices described in the main text ("Passaging of adherent HEK cells in 96 well plates") except that the dilution ratio is fixed, cell suspension is transferred by rows using a multichannel pipette, and wells are left empty as described above. 3 more source plates are seeded using an ACCS passaging protocol script designed to do the same thing but seeding one well at a time with the single-channel pipette. The protocol takes approximately 1.5 hours to run.

#### Viability assay

On each day Wednesday through Friday, one manually-seeded plate and one robot-seeded plate is assayed using CellTiter-Glo 2.0. An 11mL aliquot of thawed CellTiter-Glo 2.0 reagent and a 50mL tube of growth medium are moved from the refrigerator to ambient conditions on the morning of the measurement (at least 1 hour in advance) to allow them to equilibrate to room temperature. After imaging in the Spectramax is done, a white bottom seal is applied to the plate.

To prepare the plate: 100 µL of media is removed from all wells except for C3, C7, F3, F7. 100 µL of fresh media ("cells-" negative control) is added to C3 and F7 (round 2: F3 and C7). A stock is prepared from freshly dissociated cells, the concentration is measured with a Countess II cell counter, and dilutions are made at nominally 200k, 400k, 600k, and 800k cells/mL. Columns 5 and 10 are seeded with 100 µL of suspension each to create reference wells containing 80k, 60k, 40k, and 20k cells per well in a repeating pattern. Once all these additions are made to the plate, it is allowed to rest for 30 minutes to ensure it is in thermal equilibrium.

To initiate the assay, 100 µL of CellTiter-Glo 2.0 reagent is added to all wells except the two from which no media was removed ("reagent-" negative control ). The lid is discarded and an optically clear top seal is applied. A Spectramax i3x plate reader is used to perform both the necessary mixing and the assay readout. The read is carried out in all-wavelength mode with a read height of 6.51mm and an integration time of 500 ms. The plate reader is configured to apply orbital shaking at the "high" intensity setting for 5 minutes (we have found that results are more inconsistent when shaking for only 2 minutes as suggested by the technical manual), then read the plate every 2 minutes for a total of 8 reads. In the analysis presented here we use the read around the 10 minute mark from the start of the run.

The 40k cells/well reference group was chosen as a scale ruler for the assay results as this cell density is well within the range over which CellTiter-Glo 2.0 is advertised to be highly linear. The raw luminance reads are zero-corrected with respect to that plate's "cells-" negative control group, then the resulting net values are scaled such that that plate's 40k cells/well reference group corresponds to a value of 1.0 in order to normalize for overall scale differences due to different source plate densities, etc.

Note that in practice, we find CellTiter-Glo 2.0 responds differently to a fresh cell suspension vs an established adherent culture, even with the increased mixing time applied in this round of measurements, so for this application we do not present the assay values as absolute cell counts.

#### Live imaging

The "round 2" plates were each imaged at 10X in phase contrast at 6 preselected sites just before being assayed. To reduce experimenter bias, the site selection was determined by random draw. Sites were defined by a 3x3 grid circumscribed by the border of each well; for each capture, the microscopist was directed to move the stage to place the FOV center inside the indicated square while viewing under 4X magnification, then switch to 10X and make only minimal position adjustments before taking the image.
In addition to the above, all 12 source plates and all 12 test plates were also imaged in their entirety in brightfield at 4X once per day for diagnostic purposes. 4X imaging was automated using a Spectramax i3x plate reader with MiniMax imaging cytometer. The instrument's chamber was set to 25°C when scanning plates headed for assays that day; for plates still in culture the temperature was set to 37°C.

A square circumscribed by the edge of the well bottom was divided into a grid of 9 squares, numbered 1-9 from left-to-right and top-to-bottom (i.e. 1 is upper left, 7 is lower left). 6 wells were pseudo-randomly selected, then for each well a grid location was selected. Two of the sites were arbitrarily changed due to favoring diversity in locations over true randomness. The technique for positioning the scope at locations other than #5 was to move to the corresponding side or corner of the well and then back towards the center until the well wall was off screen. For #5 the image would just be centered on the bright area in the middle of the well. After the  initial position was acquired, the scope would be switched to 10X and phase illumination, focus was set for best contrast, and the image was taken. Minor repositioning was allowed for technical reasons, e.g. if too close to the opaque wall such that it occluded the illumination and reduced contrast. The goal was not to match exactly the same field of view every time, only to streamline the imaging task and reduce bias from unconscious "cherry picking" of wells/sites.

#### Data analysis

Data analysis and figure generation was performed with Python, Numpy and Matplotlib. See the Data Availability section below.
TIF images from the EVOS microscope were bulk converted to JPG format and individually given linear brightness and contrast adjustments for legibility. Raw images and unretouched JPEG versions can be found in the repository at the location mentioned in the Results section under "Morphology images."

### Data availability

All supplemental materials, code, and documentation related to the ACCS publication can be found by starting at the main public repo:
https://github.com/czbiohub-sf/2024-accs-pub

Copies of images, protocol scripts, logs, and raw data to accompany this supplement can be found at:
https://github.com/czbiohub-sf/2024-accs-pub/accs24-pub-aux-data/supplement_e/
